# Temporary deterioration of health and behavior during pexidartinib-mediated microglia depletion and repopulation in progranulin-deficient mice

**DOI:** 10.64898/2026.04.20.719642

**Authors:** Marc-Philipp Weyer, Lisa Hahnefeld, Luisa Franck, Yannick Schreiber, Carlo Angioni, Michael K.E. Schäfer, Irmgard Tegeder

**Affiliations:** Institute of Clinical Pharmacology, Goethe University Frankfurt, Faculty of Medicine, Theodor-Stern-Kai 7, 60590 Frankfurt am Main, Germany; Fraunhofer Institute for Translational Medicine and Pharmacology ITMP and Fraunhofer Cluster of Excellence for Immune Mediated Diseases CIMD, Theodor-Stern-Kai 7, 60596 Frankfurt am Main, Germany; Department of Anaesthesiology, University Medical Center, Johannes Gutenberg-University, 55131 Mainz, Germany; Focus Program Translational Neurosciences (FTN) of the Johannes Gutenberg-University, 55121 Mainz, Germany

**Keywords:** Progranulin, microglia, synapses, PLX3397, pexidartinib, IntelliCage, lipidome, behaviour

## Abstract

Progranulin (PGRN) is a neurotrophic and anti-inflammatory factor produced mainly by neurons and microglia in the central nervous system. Progranulin haploinsufficiency causes frontotemporal dementia (FTD). In a previous study we showed that transgenic restoration of progranulin in neurons in progranulin knockout mice (NestinGrn KOBG <u>k</u>nock<u>o</u>ut <u>b</u>ack<u>g</u>round) did not prevent the dementia-like phenotype. Here, we assessed if pharmacologic microglia depletion via PLX3397-diet (CSF1R-antagonist) had therapeutic value in these mice. Microglia depletion and spontaneous repopulation was confirmed in immunofluorescence and rtPCR studies. There was no difference in depletion or repopulation efficiency between NesGrn KOBG, PGRN KO and heterozygous (het) PGRN mice, but microglia repopulated faster than in control Grn-flfl mice, and the morphology of primary PGRN deficient microglia during repopulation was closer to homeostatic microglia, and it was accompanied by a remarkable restoration of dendritic spines and synaptic structures. Regardless of these positive effects, NesGrn KOBG and PGRN het mice experienced serious side effects during microglia depletion which peaked around the microglia nadir. Overactivity and excessive grooming escalated and caused serious skin lesions. Bulk transcriptomic and metabolomic studies in the brain taken 8 weeks after the end of PLX-diet clearly revealed differences between genotypes but mostly no lasting impact of PLX-diet, except for a further increase of proinflammatory genes, cathepsins and complement factors in PLX-treated groups. Cell type specific lipidomic studies revealed a time dependent switch not only in microglia but also astrocytes upon PLX3397 treatment. While nadir-microglia were triglyceride-laden, repopulated microglia returned to normal TG levels but were enriched in ether-bound phosphatidylcholines (PC-O) and lysophosphatidylglycerol species which are pro-inflammatory lipids; and astrocytes overtook the TG burden during repopulation. Our data suggest that microglia depletion may cause a deterioration in progranulin-deficiency.

## Introduction

Progranulin (PGRN) is a secreted, pleiotropic protein with neurotrophic and anti-inflammatory properties, produced predominantly by neurons and microglia in the central nervous system [1]. Several receptor systems, including Notch and EphA2 [2, 3] as well as TNFα receptors [4], have been implicated in mediating its extracellular actions, which depend on extracellular PGRN availability, tightly regulated through sortilin-dependent endocytosis and lysosomal trafficking [5]. Beyond these extracellular roles, PGRN has emerged as a key intracellular regulator of phagocytosis, autophagy, and lysosomal degradation [6, 7], particularly by supporting autophagolysosomal flux. How lysosomal PGRN enhances degradative capacity from within the organelle remains incompletely understood [8]. Loss-of-function mutations in *Grn* cause severe neurodegenerative diseases, most prominently frontotemporal dementia (FTD) and neuronal ceroid lipofuscinosis [9–12]. Mouse models lacking *Grn* have therefore been instrumental in demonstrating that restoring or augmenting PGRN expression—genetically or pharmacologically—can ameliorate disease-relevant phenotypes [13–18]. These findings have motivated the development of therapeutic strategies ranging from AAV-mediated local brain or systemic gene delivery [13, 17, 19, 20] to BBB-penetrant PGRN-mimetic peptides [21], and have led to ongoing clinical trials of *Grn* gene therapy (NCT04747431, NCT04408625, NCT06064890) and sortilin-blocking antibodies (NCT04374136) designed to elevate extracellular PGRN [14, 22], the latter so far without success [23].

Most central gene-replacement approaches have relied on AAV9-mediated *Grn* expression, which bypasses sortilin and primarily targets neurons [24]. Although intraventricular AAV9-*Grn* delivery achieves broad CNS expression in knockout mice, supraphysiological levels can provoke region-specific toxicity, as reported in the hippocampus [25], and after brain injury [26]. Other studies have shown partial rescue of behavioral, cellular, and lysosomal phenotypes in *Tmem106b-Grn* double-knockout mice [13] or in heterozygous *Grn*+/− mice [16]. A first-in-human trial of intracisternal AAV9-*Grn* (PR006) demonstrated increased CSF PGRN but also elicited anti-AAV9 immune responses and CSF pleocytosis [24]. These observations underscore that neuronal PGRN restoration is feasible but carries risks and may not be sufficient to achieve a durable therapeutic benefit.

Indeed, our recent work showed that genetic neuronal PGRN restoration alone did not correct dementia-like behavior or gliosis in mice, although it did mitigate synaptic spine loss, a phenotype linked to aberrantly reactive neuron-attacking microglia [27–31]. In contrast, a peripheral, liver-targeting viral approach that used the transferrin receptor as a carrier and restored PGRN in both neurons and microglia was able to nearly eliminate the progranulin-associated pathology in a mouse model [20]. These findings suggest that effective therapy may require not only neuronal rescue but also a resetting of microglial identity and function.

Pharmacological microglia depletion followed by repopulation has emerged as a promising strategy to replace maladaptive microglial states with more homeostatic, neuron-supportive populations [32–34]. Repopulated microglia often exhibit normalized morphology and transcriptional profiles, and they can attenuate pathology in models of neurodegeneration [35–38], traumatic injury [39], and vascular damage [40]. Depletion is typically achieved with CSF1R inhibitors such as pexidartinib (PLX3397) [41], after which microglia repopulate spontaneously. Repopulation arises mostly from residual PLX-resistant microglia, and by few infiltrating monocytes that adopt microglia-like morphology [42] while remaining genetically distinct [43]. This plasticity has opened the door to combining depletion–repopulation paradigms with transplantation of gene-edited microglia or stem-cell-derived precursors. Supporting this concept, a recent study showed that busulfan– and PLX3397-conditioned *Grn*-deficient mice receiving an allogeneic bone marrow transplantation regained microglial PGRN, normalized lysosomal function, and corrected lipid metabolism [44]. These results suggest that microglial PGRN is as essential as neuronal PGRN for disease correction.

In the present study, we therefore investigated whether PLX3397-driven microglia renewal, in the presence or absence of neuronal PGRN restoration, is sufficient to ameliorate the behavioral and pathological consequences of progranulin deficiency in mice.

## Methods

### Animals

PGRN-KO mice [45] (gift from Aihao Ding, Grn^tm1Aidi^, MGI:4421704) were crossed with Gt(ROSA)26Sor^tm1.1(Ubc-Grn)Ite^ (MGI: 6149573), referred to as Grn-flfl mice [7] and with Nestin-Cre mice (B6. Cg-Tg(Nes-cre)^1Kln/J^). To generate the Grn-flfl mice, a targeting vector consisting in the murine cDNA of progranulin, upstream ubiquitin promoter and a loxP flanked STOP codon (fl-STOP-fl) was inserted via homologous recombination into the Rosa26 locus, leading to the generation of Gt(ROSA)26Sor^tm1.1(Ubc-Grn)Ite^ mice (MGI: 6149573) [7]. Subsequently, the transgene expression of progranulin by Cre-recombinase was achieved by crossing Grn-flfl mice with Nestin Cre mice (B6.Cg-Tg(Nes-cre)^1Kln/J^), thereby removing the floxed STOP-codon. The resulting mice were then crossed with progranulin knockout mice (B6.CD1-*Grn*^tm1Ding^) referred to as PGRN KO mice [45]. The breeding strategy is shown in [31].

Triple heterozygous offspring were further crossed to homozygosity for the PGRN knockout allele (PGRN KO) and the loxP-STOP-loxP-mGrn allele (Grn-flfl) and to hemizygosity of Nestin-Cre (NesCre). Cre-positive mice exhibited exclusively neuronal PGRN expression. This line is referred to as NesGrn KOBG. The short name stands for Nestin driven *Grn* expression in <u>k</u>nock<u>o</u>ut <u>b</u>ack<u>g</u>round where the endogenous *Grn* was deleted. Cre-negative animals carry an inactive floxed allele and are therefore PGRN KO mice. A triple genotyping assay consisting of 6 probes was developed and is available from Transnetyx (Line: NesGrn KOBG). All mice were on a C57BL6 genetic background. Suppl. Table S1 provides an overview of sample sizes, ages and sex of mice used for behavioral and biological experiments.

Mice had free access to water and food and were kept in climate-controlled rooms with a 12 h light-dark cycle. The behavioural studies were approved by the local Ethics Committee for animal research (Darmstadt, Germany) under FK1103 and FU2080. The studies adhered to the guidelines of the Society of Laboratory Animals (GV-SOLAS) and were in line with the European and German regulations for animal research and the ARRIVE guideline.

### Treatment

Mice were assigned randomly to PLX3397 diet or standard chow. PLX3397 (MedChemExpress) food pellets were generated by ssniff (ssniff Spezialdiäten GmbH) yielding a concentration of 330 mg/kg standard lab chow. PLX-food was provided ad libitum for 14 days, and control animals received vehicle chow (colored without drug). The efficiency of dietary PLX3397 administration in mice was shown before [46]. Food intake and body weight of mice were determined twice a week during treatment and 1x/week during washout. Mice were euthanized directly at the end of the 14d-depletion diet referred to as “d0 repopulation” and at day-8 (d8) and week-8 (W8) after stopping PLX-diet (d8 repopulation, W8 repopulation). Several cohorts were investigated for behavior, histology, gene expression, lipidomics/metabolomics and primary microglia cultures. Biological analyses (RNAseq, lipidomics/metabolomics) were done in randomized order without knowledge of genotype or treatment.

### Mouse tissue collection: brain and plasma

Mice were euthanized with carbon dioxide and blood withdrawal by cardiac puncture, whereby blood was collected into K3^+^ EDTA tubes, centrifugated at 1300 *g* for 5 min, and plasma was transferred to a fresh tube and snap frozen on dry ice or in liquid nitrogen. The brain was dissected for lipidomic and transcriptomic analyses. Cerebellum and olfactory bulb were removed, and the brain was cut sagittal. Left and right halves were weighed with precision scales and snap frozen on dry ice. Samples were stored at –80 °C until analysis.

### Brain tissue processing and immunofluorescence analysis

Mice were terminally anesthetized with carbon dioxide and ketamine and transcardially perfused with cold phosphate buffered saline (PBS) followed by 2.25% paraformaldehyde (PFA) for fixation. Tissues were excised, postfixed in 2.25% PFA for 2 h, cryoprotected overnight in 20% sucrose at 4°C, embedded in small tissue molds in cryo-medium and cut on a cryotome (12 μm). Slides were air-dried and stored at −80 °C. Immunofluorescence staining was performed with air-dried cryosections. After washing in 1x PBS, sections were blocked in 0.1% Triton X-100/5% BSA/PBS at room temperature (RT) for 120 min and incubated in primary antibody solution (IBA1, CD68, CD11b, GFAP) solution consisting in 0.1% Triton X-100 and 1% BSA in PBS overnight at 4°C. After washing, sections were incubated in secondary antibody for 2 h at RT. The steps were repeated for the second primary/secondary antibody pair (GFAP). Finally, nuclei were counterstained with DAPI for 10 min at RT, washed again and mounted with Aqua-Poly/Mount (Polysciences). Synaptic structures were analyzed accordingly using immunofluorescence studies with anti PSD95 and anti-synaptophysin. Antibodies are listed in Suppl. Table S5.

### Golgi-Cox staining of brain tissue and quantification of spine density

Golgi–Cox staining was performed using the FD Rapid GolgiStain™ Kit (FD NeuroTechnologies) following the manufacturer’s guidelines. After CO₂ euthanasia and transcardial perfusion with PBS and 2.25% PFA, the brain was removed and a ∼5 mm frontal block (Bregma +0.5–2.5) was prepared. Tissue was immersed in the impregnation solution, which was refreshed after 24 h, and kept at room temperature for 14 days in the dark. Samples were then transferred to Solution C for 72 h (with one solution change), embedded in 2% low-melting agarose, and sectioned at 100 µm on a vibratome.

Sections were mounted on Superfrost slides and air-dried for 72 h in the dark. Before staining, slides were rinsed in Milli-Q water and incubated for 10 min in freshly prepared staining solution (Solutions D and E diluted in water). After additional water washes, sections were dehydrated through graded ethanol, cleared in xylene, and coverslipped with Pertex. Slides were dried overnight at 37 °C and stored at room temperature protected from light.

Images of layer II/III pyramidal neurons in the frontal cortex were acquired with a Keyence BioRevo BZ9000 microscope (RRID:SCR_015486) using a 40× objective lens. For each image, one or two dendritic segments of 30 ± 1 µm were analyzed in FIJI/ImageJ to determine spine density, which was normalized per 10 µm dendrite length.

### Image acquisition and analysis

Immunofluorescence images were captured using fluorescence microscopy (BZ-X800, Keyence) or confocal scanning microscopy (Leica Stellaris 8, RRID:SCR_024660). FIJI ImageJ (ImageJ, RRID:SCR_003070) was used for quantitative analysis of the immunoreactive (IR) area relative to the area of the region of interest (ROI) with adequate threshold setting using the particle analyzer plugin of ImageJ. The ROI was identical in size for all mice/sections. Three images of non-adjacent sections of each three to five mice were analyzed per region (motor cortex, hippocampus, thalamus, temporal cortex), resulting in 9-15 data points per genotype per region.

### IntelliCage behavior

Behavioural analyses were done with unbiased IntelliCages. The IntelliCage (TSE, Berlin Germany) consists of four operant corners, each with two water bottles, sensors, LEDs, and doors that control the access to the water bottles. The system fits into a large Techniplast 2000P cage (20 × 55 × 38 cm) and allows housing of 16 mice per cage. Four triangular red houses are placed in the center to serve as sleeping quarters and posts to reach the food. The floor is covered with standard bedding. Mice are tagged with radiofrequency identification (RFID)-transponders, which are read with an RFID antenna integrated in the corner entrances. The corners give access to two holes with water bottles, which can be opened and closed by automated doors. Mice must make nosepokes (NP) i.e. peak through a light barrier to open the doors for water access. The IntelliCage is controlled by IntelliCage Plus software, which executes pre-programmed experimental tasks and schedules. The numbers and duration of corner visits, NP, and licks are recorded continuously without the need for handling of the mice during the recording times. Learning and memory can be supported by LEDs.

IntelliCage tasks address several different aspects of cognition as well as circadian rhythms and social interactions and were run sequentially. The tasks followed previously established protocols [47–49]. The IntelliCage experiments were done in old <u>female</u> mice to avoid fighting. Up to 16 mice were housed per cage. Mice were adapted to the cages for one week with free access to every corner, with all doors open, and water and food ad libitum. This free adaptation (FA) was followed by “nosepoke adaptation” (NP) for three weeks in which the doors were closed. The first NP of the visit opened the door for 5 s. To drink more, a mouse has to leave the corner and start a new visit. In the place preference learning (PPL1) task mice learned to prefer a specific corner, in which an NP opened the door to get water access. Each 4 mice were assigned to one corner. After conditioning to the corner for one week, the rewarding corner was switched to the opposite side (PPL reversal learning (PPL2) for another week, and then again to a different long-side distant corner (PPL3). In the final PPL4social task (PPL4s) all mice of one cage were assigned to one rewarding corner to observe social hierarchies and competition, albeit without additional time limitation (soft social competition). Outside of the module-active times, the doors remained closed. The mice in cage-1 received PLX3397 diet during the second and third week of the NP task.

### Assessment of thermal sensitivity (Thermal Gradient Ring (TGR))

A thermal gradient ring (TGR, Stoelting) was used to assess the temperature preferences and the exploration of the ring platform that consists of a circular ring platform that allows free choice of the comfort zone [50, 51]. The dimensions of inner and outer ring diameters are 45 cm and 57 cm. The inner walls consist of plexiglass and the outer walls of aluminum. Both are 12 cm high and build a 6 cm wide circular running arena. The aluminum surface provides a temperature gradient that is controlled with two Peltier elements and constantly measured with infrared cameras. The arena is divided into mirror-image semicircles of 12 temperature zones, so that duplicate readouts are provided for each zone. During measurements, the running track is illuminated, and the mouse behavior is videotaped with a regular CCD camera, mounted above the mid-point of the ring. The time spent in zones, temperature preferences, travel paths and distance, and body rotations are analyzed unbiased with the TGR ANY-Maze video tracking software (Stoelting). The time of grooming and rearing was manually recorded by pressing a defined key during the first quarter of the habituation as well as first and fourth quarter of the test.

### mRNA sequencing of mouse brain tissue

Total RNA was extracted from fresh frozen brain tissue, which included cortex and subcortical structures without olfactory bulb, cerebellum, and brain stem. Briefly, total RNA was purified using an RNAeasy plus micro kit (Qiagen), and the quantity and quality assessment was checked by Qubit Flex and RNA 6000 Nano chip on Agilent’s bioanalyzer, respectively. Next Generation paired-end mRNAseq from mouse brain tissue was performed at Novogene (Cambridge, UK) on an Illumina NovaSeq 6000 platform. Sample quality was evaluated using demultiplexed fastq.gz files. Sequenced reads were subjected to adapter trimming and processing via CLC Genomics workbench (Qiagen, v25) using standard setting for sequence alignment. Sequence reads were annotated according to the mouse genome mm10 assembly. Results were visualised using CLC expression browser, encompassing the number of mapped reads, target length, source length and position, strand, genes and gene IDs. Read counts were normalized using the EdgeR algorithm providing the trimmed mean of M-values, TMM (in CLC called CPM). TMM reads were Log2 transformed.

Differential gene expression was assessed using Principal component analyses, ANOVA-like methods, t-tests and fold change in CLC genomic workbench. Low expression genes were filtered out. The P value was set at 0.05 and adjusted according to the False Discovery Rate (FDR). Hierarchical clustering with Euclidean distance metrics was used to assess gene expression patterns. Results were displayed as heat maps with dendrograms. Genes were ranked according to P and q value, fold change and abundance. Ranked genes were submitted to gene ontology enrichment analysis using ShinyGO (https://bioinformatics.sdstate.edu/go/)[52] and the web tool Gorilla (https://cbl-gorilla.cs.technion.ac.il/) [53]. The RNAseq data have been deposited at the GEO database with the provisional accession number GSE287561.

### Primary microglia culture

Primary microglia were isolated using the Miltenyi Adult Brain Dissociation Kit in combination with CD11b MicroBeads according to the manufacturer’s protocol. Following CO₂ euthanasia and blood withdrawal by cardiac puncture, mouse brains were collected in HBSS and enzymatically dissociated with the gentleMACS Octo dissociator system using the manufacturer-provided enzyme mixes and DNAse I. The resulting suspension was filtered, subjected to debris removal and red blood cell lysis, and washed in HBSS to obtain a clean single-cell preparation.

For magnetic enrichment, cells were incubated with CD11b MicroBeads in protein-containing binding buffer and applied to MiniMACS MS columns placed on a magnetic stand. After washing, the flow-through (non-microglial fraction) was collected and stored at −80 °C. CD11b⁺ microglia were eluted from the columns with binding buffer, pelleted, and resuspended in prewarmed DMEM/F12-GlutaMax medium supplemented with 10% FCS and 1% Pen/Strep.

Cells were counted using a Neubauer chamber, and approximately 50,000 microglia were seeded per well of an 8-well culture slide for downstream experiments or processed directly for RNA extraction and qRT-PCR.

### Immunofluorescence analysis of primary microglia

Microglia were immune stained with anti-IBA1, and subsequent anti-mouse Alexa Fluor 488, and DAPI as nuclear counter stain. Microglia images were captured on a BioRevo BZ9000 Keyence microscope (RRID:SCR_015486). Microglia lipid load was assessed with BODIPY staining, and counterstained with anti-Lamp1 to identify lysosomes. JC-1 immunofluorescence analysis was used to assess the mitochondrial membrane potential in primary microglia cultures and in isolated synaptosomes as well as mitochondria from the mouse brain. The dye accumulates in mitochondria in a potential-dependent manner, forming red-fluorescent J-aggregates in polarized (high-potential) mitochondria, whereas depolarized mitochondria retain JC-1 in its monomeric, green-fluorescent form. The red/green fluorescence ratio was therefore used as an indicator of mitochondrial membrane potential.

For quantification of microglia morphology, images were converted into 8-bit binary images using the trainable weka segmentation tool in FIJI ImageJ and skeletonized. Subsequently, microglia morphometry was assessed by using the Fractal Generator (FracLac) plugin in ImageJ. The following parameters were obtained: (1) total foreground pixel within the ROI describing the size of the cell area (CA) including branches, (2) diameter of the surrounding circle, (3) area of a hull polygon connecting the tips of the outmost branches, (4) pixel density, defined as the ratio of the cell area (foreground pixel) to the area of hull polygon, (5) length-to-width ratio of the hull polygon, (6) counts of proximal branches, (7) counts of branch tips, (8) counts of junctions, (9) counts of triple junctions, (10) total lengths of branches, (11) ramification factor which is the ratio of proximal branches to soma size, (12) fractal dimension (FracD), which is an index describing the degree to which a complex structure fills out a graphic area and was calculated by a box-counting algorithm, (13) circularity, calculated according to the formula: circularity = 4π × cell area/CP^2^ where the cell perimeter (CP) is compared to that of a circle.

Multiple cells were analyzed per image that were captured from primary cultures of at least three mice per genotype and treatment. For multivariate analyses, measures for the parameters were log2-transformed because the data were mostly log-distributed, except for fractal dimension and pixel density which were used as linear data. Transformed data were subjected to supervised Canonical Discrimination and Random Forest analysis to assess if/how complex microglia morphology predicted the genotype or treatment. Data for individual parameters were submitted to Brown-Forsythe and Welch ANOVA and subsequent Dunnett’s T3 multiple comparisons test.

### Isolation of synaptosomes and mitochondria

Mice were euthanized by CO₂ and cardiac puncture. Brains were excised, the cerebellum removed, and tissue washed in freshly prepared sucrose–tris–EDTA (STE) buffer (pH 7.4). Each brain was homogenized in STE buffer using a Dounce homogenizer, followed by low-speed centrifugation (3 min, 1300 g, 4 °C). The pellet was re-homogenized and centrifuged again, and the pooled supernatants were cleared at 21 000 g for 10 min at 4 °C. Pellets were resuspended in 15% Percoll in STE buffer and layered onto a discontinuous 23%/40% Percoll gradient. After ultracentrifugation (30,700 *g*, 10 min, 4 °C), the 15-23% interface (synaptosomes) and 23-40% interface (mitochondria) were collected, diluted in STE buffer, and pelleted (10 min, 16,700 *g*, 4 °C). Final pellets were resuspended in TES buffer.

For JC-1 staining, 50 µl of the suspension was mixed with 250 µl JC-1 (1:2500 in TES buffer; stock 5 mg/ml) and incubated for 20 min at room temperature. Imaging was performed on a Stellaris 8 confocal microscope (excitation 488 nm; emission 515–540 nm for green and 580–620 nm for red). For each mouse, five images were captured with 40x objective lens, and three additional images at 40x with 2.5× digital zoom for mitochondria and synaptosomes.

### Seahorse analysis of respiration in primary microglia

Mitochondrial respiration was analyzed with the Mito Stress Test (Agilent Technologies) on an XFp Seahorse analyzer. It is a microplate-based live cell assay for monitoring the oxygen consumption rates (OCR) and extracellular acidification rates (ECAR) in living cells. Microglia were prepared as described above and seeded in XFp miniplates in complete medium at 60,000 cells per well. Twenty-four hours after seeding, the culture medium was switched to Seahorse XF medium supplemented with 1 mM pyruvate, 2 mM glutamine and 10 mM glucose, and microglia were allowed to equilibrate in a non CO_2_-inbubator. Cultures were then transferred to the XFp analyzer and subjected to the mitochondrial stress test protocol. Basal respiration was measured under full nutrient supply. Subsequently, oligomycin (1.5 µM) was added to assess ATP generation, proton leakage and extracellular acidification. Oligomycin blocks ATP synthase, manifesting in a drop of OCR and raise of ECAR. In the next step, FCCP (carbonyl cyanide-4-(trifluoromethoxy)-phenylhydrazone; 0.5 µM and 1.0 µM) was injected to assess maximum oxygen consumption. FCCP causes a leak of H^+^ across the mitochondrial membrane leading to a collapse of the membrane potential. Finally, residual non-mitochondrial respiration was determined after inhibition of complex I with rotenone (0.5 µM) and complex-III with antimycin A (0.5 µM). Finally, cultures were imaged for evaluation of equal density. Oxygen consumption rate (OCR) and extracellular acidification rate (ECAR) were analyzed using the Seahorse XF Wave® software.

### mRNA sequencing of primary mouse microglia and neurons/astrocytes

Total RNA was extracted from primary cells as described above. RNA-seq libraries were generated using the Lexogen QuantSeq 3′ mRNA-Seq Library Prep Kit, FWD (forward) according to the manufacturer’s instructions. In brief, first-strand cDNA synthesis from total RNA was initiated by oligo(dT) priming, which selectively targets polyadenylated transcripts and introduces the partial Illumina Read 1 adapter sequence. After reverse transcription, the RNA template was enzymatically removed, and second-strand synthesis was performed using random primers carrying the complementary Illumina adapter sequence, thereby generating double-stranded cDNA fragments close to the 3′ end of transcripts. The resulting libraries were purified using magnetic beads, followed by PCR amplification to complete adapter sequences and introduce i7 indices for multiplexing. Final libraries were purified, quantified, and assessed for size distribution before sequencing on an Illumina NexGen 2000 platform. Data processing was done as described above, and the RNAseq data have been deposited at the GEO database with the provisional accession number GSE324813.

### Gene expression analysis using quantitative rtPCR in primary microglia

For *Grn* gene expression analysis in primary neural and glial cells and brain tissue RNA was isolated from primary microglia and neurons/astrocytes obtained during the isolation protocol. The Qiagen All Prep DNA/RNA/Protein Isolation Kit and the Qiagen RNA Isolation Kit were used according to the manufacturer’s instructions. The amount of isolated RNA and purity were measured on a Nanodrop spectrophotometer. The Verso cDNA Synthesis Kit (Thermo Scientific) was used with 180-200 ng of RNA to generate cDNA via reverse transcription according to the manufacturer’s instructions. RT‒qPCR was conducted by using ORA^TM^ SEE qPCR Green ROX (highQu) in duplicate on a QuantStudio^TM^ 5 System (Thermo Fisher Scientific). Absolute values of target gene expression were normalized to the reference peptidylprolyl isomerase A (*Ppia*) and glyceraldehyde 3-phosphate dehydrogenase (*Gapdh*) values. Oligonucleotide sequences, amplicon sizes, annealing temperatures, and NCBI reference sequence numbers are provided in the Supplementary Material (Suppl. Table S4).

### Untargeted lipidomic and metabolomic analyses in mouse brain and primary cells

Mouse brain tissue samples were homogenized by adding ethanol:water with 10 µM indomethacin (1:3, v/v, tissue concentration 0.02 µg/ml) using a Precellys 24-Dual tissue homogenizer coupled with a Cryolys cooling module (both Bertin Technologies, Montigny-le-Bretonneux, France) with 10 zirconium dioxide grinding balls (3*20s at 6500 g with 60s breaks), operated at < 6 °C. Subsequently, 20 µl of the homogenate containing 1 mg of tissue were extracted using a liquid-liquid-extraction method. Primary cells (250,000 per sample) were directly extracted using the same liquid-liquid-extraction method.

Lipidomic and metabolomic analyses were conducted applying the same procedure as previously described [54]. Further protocol details for extraction and analyses are described in the supplementary methods (Excel file). Briefly, a methyl-tert-butyl-ether (MTBE) and methanol-based liquid-liquid extraction was used. For chromatographic separation of lipids, a Zorbax RRHD Eclipse Plus C8 1.8 µm 50 x 2.1 mm ID column (Agilent, Waldbronn, Germany) with a pre-column of the same type was used. The mobile phases were (A) 0.1 % formic acid and 10 mM ammonium formate and (B) 0.1 % formic acid in acetonitrile:isopropanol (2:3, v/v). Polar metabolites were separated on a SeQuant ZIC-HILIC,

3.5 µm, 100 mm × 2.1 mm I.D. column coupled to a guard column with the same chemistry (both Merck, Darmstadt, Germany) and a KrudKatcher inline filter (Phenomenex, Aschaffenburg, Germany). Using 0.1 % formic acid in water (solvent A) and 0.1 % formic acid in acetonitrile (solvent B), binary gradient elution was performed with a run time of 25 min.

Analysis was performed on an Orbitrap Exploris 480 with a Vanquish horizon UHPLC system (both Thermo Fisher Scientific, Dreieich, Germany). Data was acquired using Thermo Scientific XCalibur v4.4 (RRID:SCR_014593) and relative quantification was performed in Thermo Scientific TraceFinder 5.1 (RRID:SCR_023045). Full scan spectra were acquired from 180-1500 *m/z* for lipidomics, 70-700 *m/z* (metabolomics positive ion mode) and 59-590 *m/z* (metabolomics negative ion mode) at 120,000 mass resolution each for 0.6 sec, and data dependent MS/MS spectra at 15,000 mass resolution in between. Pooled quality controls were prepared from the first ten tissue homogenates and replicates measured along the run to verify system performance.

For all lipid analyses, the area under the curve (AUC) divided by the AUC of the Internal standard (AUC/IS) was used for quantification and statistical analyses. For cell type specific lipidomic and metabolomic studies, AUCs were normalized according to the probabilistic quotient normalization (PQN). It adjusts the median of the samples’ spectra to that of the pooled quality control. The normalization factor was specific for lipids/metabolites measured in positive or negative ion mode. For comparative lipidomic analyses of microglia and astrocytes, a sum normalization was used to adjust the total lipids which depend on cell mass and membrane size, requiring an adjustment of small microglia relative to neurons/astrocytes for inter-cell type comparative analyses. PQN-adjusted AUCs or AUC/IS were transformed to square root values to adjust a skewed distribution. For multivariate analyses sqrt AUC/IS were scaled to have a common average and standard deviation of 1 (autoscaling in MetaboAnalyst).

### Statistics

Group-level results are reported either as mean ± SD, mean ± SEM for behavioral datasets, or as median with interquartile range, as indicated in the individual figure legends. Statistical analyses were carried out using SPSS 29 (RRID:SCR_016479), GraphPad Prism versions 9 or 10 (RRID:SCR_002798), Origin Pro 2025 (RRID:SCR_014212), and MetaboAnalyst 5.0 (RRID:SCR_016723) [55]. The procedures used for processing and interpreting the various “omics” datasets (RNA-seq, lipidomics, metabolomics) as well as microglial morphology are described in the corresponding sections.

For lipidomic mass spectrometry, peak areas (AUC) were normalized to the internal standard (AUC/IS) and then square-root transformed to mitigate skewness. For multivariate analyses and heatmap visualization, variables were standardized to Z-scores (*x* − *x*ˉ)/*SD* to ensure comparable scaling. To evaluate group differences, datasets were subjected to two-way ANOVA with factors such as “feature” (e.g., lipid species, gene expression, behavioral parameter) and “group” (genotype and/or treatment). When ANOVA indicated significant effects, post hoc comparisons were performed using Šidák-corrected t-tests, false discovery rate (FDR) adjustment, or Dunnett’s test relative to the Grn-flfl control group. The interpretation of asterisks in figures is provided in the respective legends.

For lipidomic and transcriptomic datasets, Partial Least Squares–Discriminant Analysis (PLS-DA) and Random Forest classification were applied to evaluate group separation and feature importance. Volcano plots were generated to visualize fold changes against the negative log10 of the corresponding t-test P values. Hierarchical clustering using Euclidean distance and Ward’s linkage was used to generate heatmaps with dendrograms. Selection of the top 100 features was based on ANOVA P-values.

ANOVA-simultaneous component analysis (ASCA) [56] and Canonical Discriminant Analysis were employed to analyze multivariate behavioral outcomes across sequential tasks, as well as lipid or gene profiles across time points. ASCA integrates ANOVA with PCA and includes a feature-extraction step to model the main factors (“genotype” or “treatment”) and the repeated-measure dimension (“time/task”). Feature relevance was assessed using leverage (contribution to the ASCA model) and squared prediction error (SPE), which reflects how well the model fits each variable.

## Results

### Microglia depletion and repopulation efficiency

In the first set of experiments, we compared the efficacy of PLX3397-induced depletion of microglia and subsequent repopulation in progranulin-deficient mice and control mice. Immunofluorescence studies of four brain regions (Figure 1 and Supplementary Figure S1) confirmed the expected reduction in IBA1– and CD11b-positive microglia in the cortex, hippocampus, thalamus, and frontal cortex in all three genotypes (Grn-flfl, NesGrn KOBG, and PGRN KO) at the end of the PLX3397 diet (d0 repopulation), followed by a subsequent rise after eight days of repopulation. The residual microglial immunofluorescence at the lowest point is similar between genotypes, although the abundance of NesGrn KOBG and PGRN KO microglia drops from a higher initial value. In the knockout lines, microglial coverage also reached a higher final level than in the controls, indicating microgliosis at the start and faster repopulation. Interestingly, the level of CD68, a marker of activated microglia, decreased only in PGRN KO mice and remained low after PLX3397 treatment. In contrast, CD68 increased in Grn-flfl mice at d0 repopulation and in NesGrn KOBG mice at 8d repopulation in the cortex and thalamus, with no change observed at other sites. GFAP-positive astrocytes increased in NesGrn KOBG and PGRN KO mice in all regions, independently of PLX3397 treatment, demonstrating the astrocytosis characteristic of progranulin deficiency. Figure 1C shows distinct sites represented by different symbols. Suppl. Figure S1 shows each region separately.

**Figure 1.**
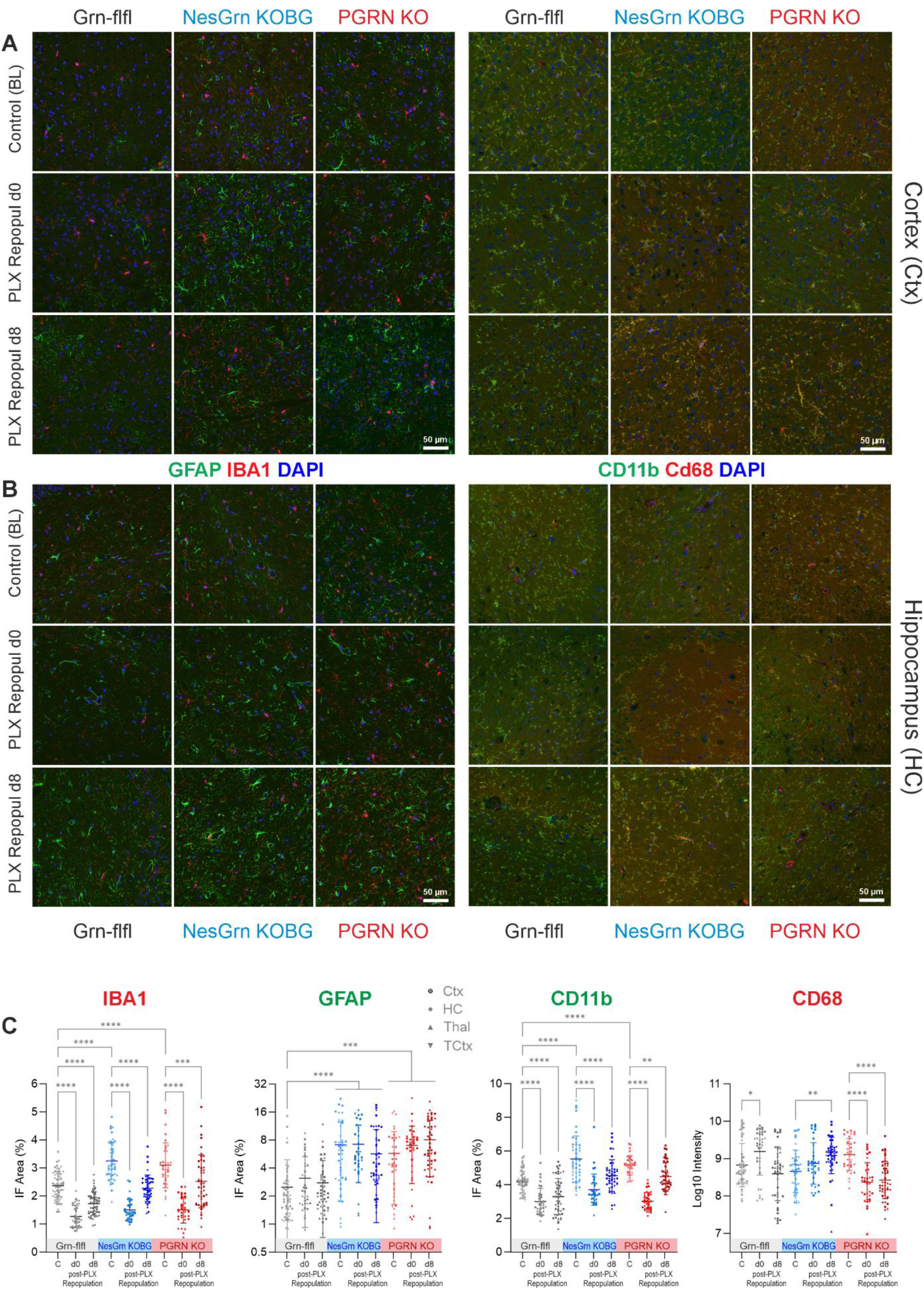
Immunofluorescence of brain microglia during PLX3397 induced depletion and repopulation. **A**: Exemplary immunofluorescent images of IBA1 immunoreactive microglia and of GFAP immunoreactive astrocytes (left panel) and of CD11b and CD68 microglia activity marker in the cortex of Grn-flfl control mice, NesGrn KOBG mice and full PGRN KO mice. DAPI is used as nuclear counterstain. **B:** In analogy to A, immunofluorescence images show the hippocampus. **C:** For quantitative analysis, images were converted to binary images using auto-threshold in FIJI ImageJ, and the relative area covered by specific immunofluorescence for IBA1, GFAP and CD11b was used for statistical comparison. For CD68, the log10-Intensity was used. The scatters show results per image of n = 3-5 mice per genotype. The line is the average, and whiskers show the standard deviation. The scatters for 4 different regions (Cortex, Hippocampus, Thalamus, Frontal cortex) are aligned but differentiated with distinct symbols. Site-specific results are shown in Suppl. Figure S1. Data were submitted to 2-way ANOVA for “brain region” by “group”, followed by posthoc analysis using an adjustment of Šidák. Asterisks reveal significant differences. P*<0.05, **<0.01, ***<0.001, ****<0.0001.

### Partial restoration of spines and synapses after PLX3397

Golgi-Cox staining of dendritic spines (Figure 2) and immunofluorescence analysis of synaptophysin-positive presynaptic sites and PSD95-positive postsynaptic sites (Figure 3) showed that PLX3397-induced depletion and repopulation of microglia was associated with an increase in synaptic structures in NesGrn KOBG and PGRN KO mice, which were seriously reduced in both progranulin-deficient lines at baseline at the onset of PLX3397 treatment. Quantification of spine numbers (Figure 2B) and synaptophysin– and PSD95-immunoreactive puncta (Figure 3C) shows almost full restoration of synapses, reaching the level of wild-type control Grn-flfl mice. In these control mice, the abundance of synaptic structures remained constant throughout PLX3397 treatment and washout.

**Figure 2.**
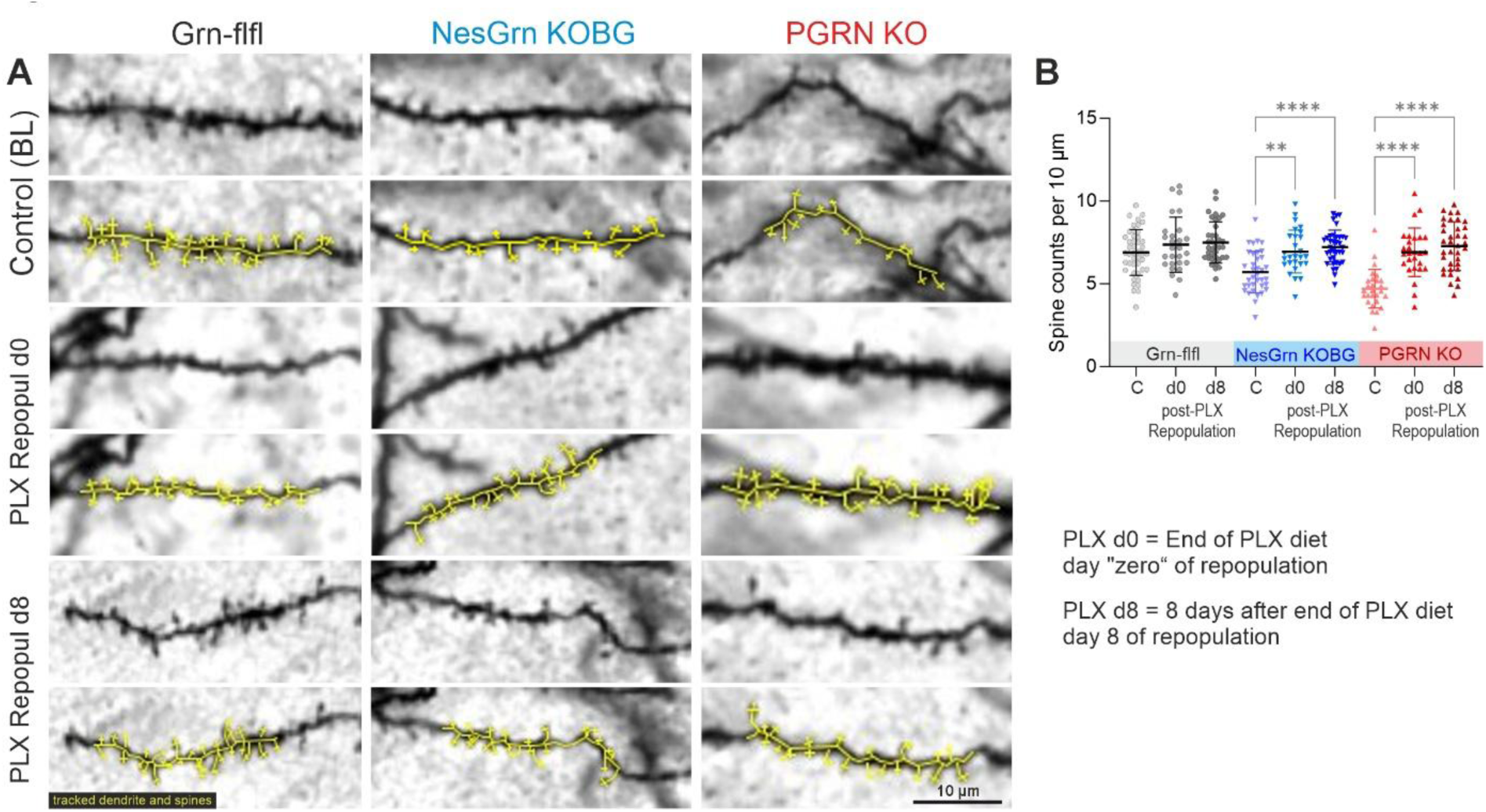
Dendritic spines of cortical neurons during PLX3397 induced microglia depletion and repopulation. **A**: Exemplary images of Golgi Cox analysis of dendritic spines in the frontal cortex of Grn-flfl, NesGrn KOBG and full PGRN KO mice before (baseline, BL) and after PLX3397 treatment for 14 days. The cortex was obtained directly at the end of the PLX3397 diet which is repopulation d0 and 8 days later (repopulation d8). The yellow markers in the respective bottom row highlight the spines, which were counted. **B:** Spine counts were normalized per 10 µm of the dendrite length representing the “spine density”. Each scatter shows the density of spines of one dendrite. At least 9 dendrites were quantified per mouse of n = 3-5 mice per genotype and treatment. Data were submitted to one-way ANOVA and subsequent posthoc comparison according to Dunnet versus the respective baseline condition for each genotype. P**<0.01, ****<0.0001.

**Figure 3.**
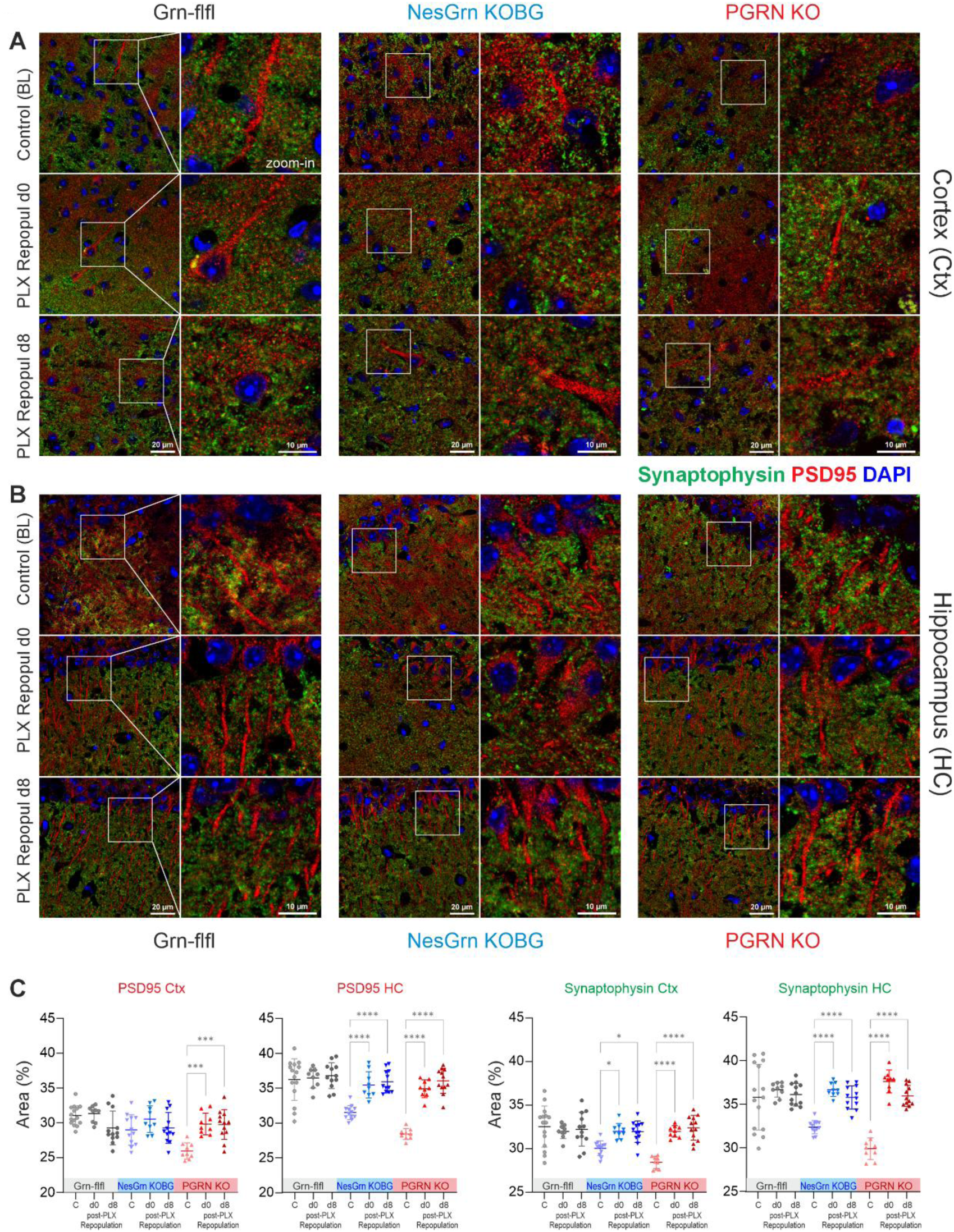
Pre– and postsynaptic densities during PLX3397 induced microglia depletion and repopulation. **A, B:** Exemplary immunofluorescent images of pre– and postsynaptic marker protein expression (Synaptophysin and PSD95) in the cortex and hippocampus of Grn-flfl control mice, NesGrn KOBG mice and PGRN KO mice before (Baseline BL) and during PLX3397 induced microglia depletion and repopulation. The brain was obtained directly at the end of 14d-PLX3397 diet period which is day-zero repopulation (repopulation d0) and 8 days later (repopulation d8). **C:** For quantitative analysis, images were converted to binary images using auto-threshold in FIJI ImageJ, and the relative area covered by specific immunofluorescence for Synaptophysin and PSD95 was used for statistical comparison. The scatters show results per image of n = 3-5 mice per genotype. The line is the average, and whiskers show the standard deviation. Data were submitted to one-way ANOVA and subsequent posthoc comparison according to Dunnet versus the respective baseline condition for each genotype. P*<0.05, ***<0.001, ****<0.0001.

### Restoration of homeostatic microglia morphology after PLX3397

The scarcity of synapses in progranulin deficiency is caused by excessive synaptic pruning [30, 57] by highly active, phagocytic microglia that express “disease-associated microglia” (DAM) marker genes, including *Lyz2*, *Gpnmb*, *Lgals3*, *Tyrobp*, *Mpeg*, cathepsins, and complement factors [31, 58–60]. Morphologically, such microglia are small and plump with short extensions (Figure 4A). Without PLX3397 treatment, we observed a biphasic distribution of microglia phenotypes in primary microglial cultures from all progranulin-deficient mouse lines, including NesGrn KOBG, PGRN KO, and PGRN heterozygous (het) mice (Figure 4B). The peaks represent “normal” homeostatic microglia, which prevail in Grn-flfl mice, and DAMs, respectively. This categorization is based on the distribution of Canonical Discrimination Factor 1, which uses multiple morphological features (see Supplementary Figure S2) to describe size, shape, branch number and arborization. Following PLX3397 treatment and two weeks repopulation, the second peak representing the DAMs was no longer evident, as revealed by a comparison of key raw features such as hull area, branch length and density (Figure 4C). In Grn-flfl mice, repopulated microglia after PLX3397 treatment show less segmentation, indicative of a more youthful phenotype. Repopulated microglia in progranulin-deficient mice exhibit a comparable youthful phenotype but lack the ’phagocyte-like’ characteristics of neuron-aggressive DAMs.

**Figure 4.**
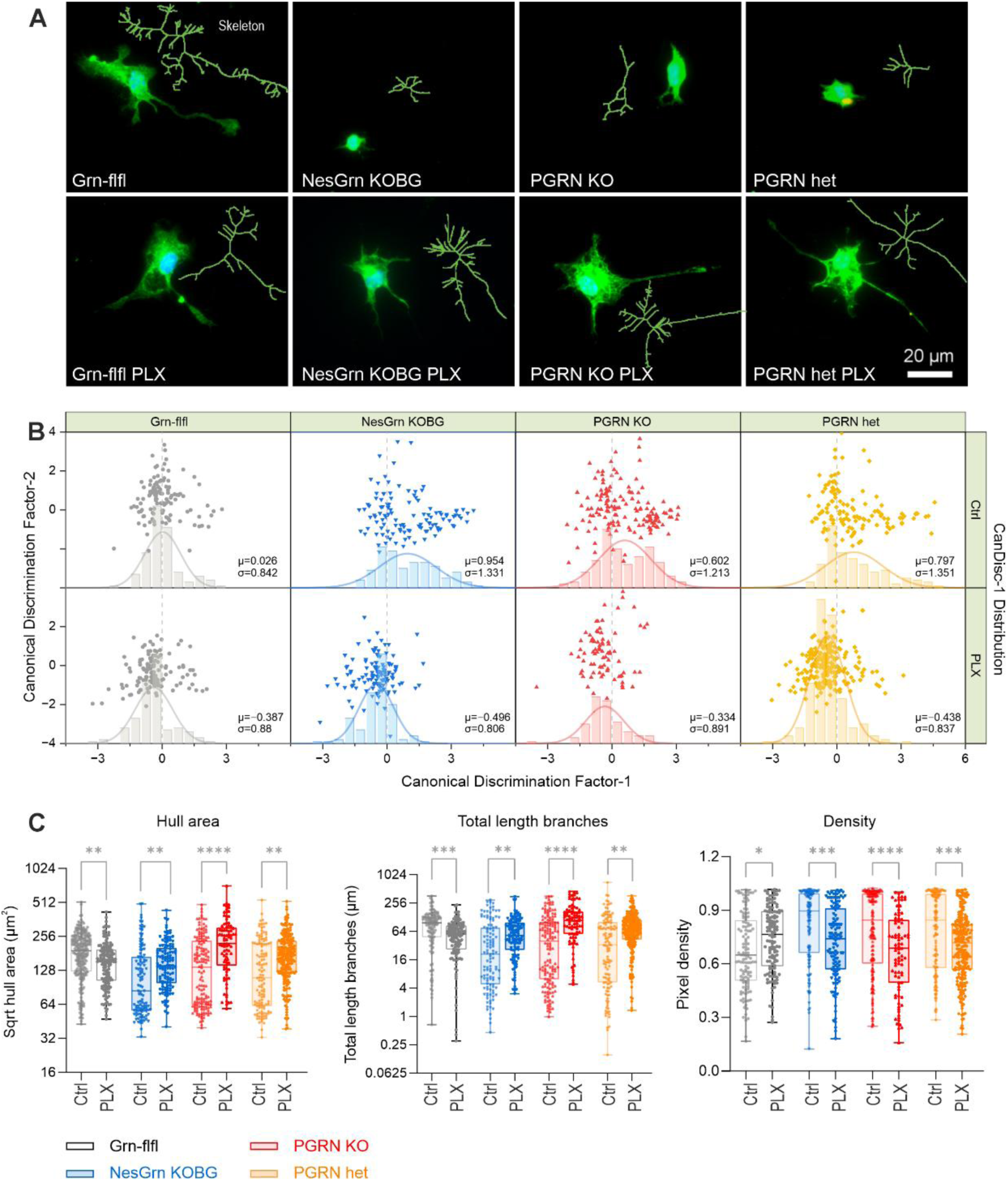
Microglia morphology and subpopulations. **A**: Microglia morphology of primary microglia from old mouse brains (14-22 months), stained with anti-IBA1 and DAPI. Inserts in green show the skeleton, which was used for analysis. Further details of the morphometry are shown in Suppl. Figure S2. **B:** Canonical discrimination analysis of complex microglia morphology features (Supp. Fig. S2) was used to assess subpopulations in control and PLX-treated mice, at day-14 of the repopulation (PLX). CanDisc scores for factor-1 and factor-2 are presented as XY scatter plots. The distribution of CanDisc-1 scores shows a biphasic distribution revealing “healthy” and “activated” microglia in untreated progranulin deficient mice. **C:** Quantitative analysis of Hull area, total branch lengths and density which describes the occupation of the polygon hull with immunofluorescent cell structures. A total of 85-232 cells of 3-8 animals per genotype were analysed. The box is the interquartile range, the line is the median, and whiskers show minimum to maximum. The data were submitted to Brown-Forsythe and Welch ANOVA and posthoc analysis using a correction of alpha according to Dunnett versus the respective control condition; P*<0.05, **<0.01, ***<0.001, ****<0.0001.

### Microglia lipid accumulation and respiratory functions not improved with PLX3397

Previous studies have shown that progranulin-deficient microglia accumulate large amounts of neutral lipids [21, 61, 62] and cause substantial oxidative damage, suggesting defects in lipophagy and respiratory functions that do not necessarily align with morphological features. Therefore, we studied lipid deposits and respiratory functions in primary microglia isolated from PLX-treated versus control mice (see Figure 5). BODIPY immunofluorescence of lipid droplets shows large lipid deposits in progranulin-deficient mice (Figures 5A and 5B), in both microglia obtained at baseline and from mice treated with PLX3397, suggesting that lipid-laden microglia are beyond the reach of PLX3397 therapy. Furthermore, microglia from PGRN KO mice exhibited enlarged lysosomes (Figure 5A, B), which were unaffected in NesGrn KO mice, suggesting that an extracellular supply of progranulin from neurons was sufficient to enhance the autophagolysosomal flux.

**Figure 5.**
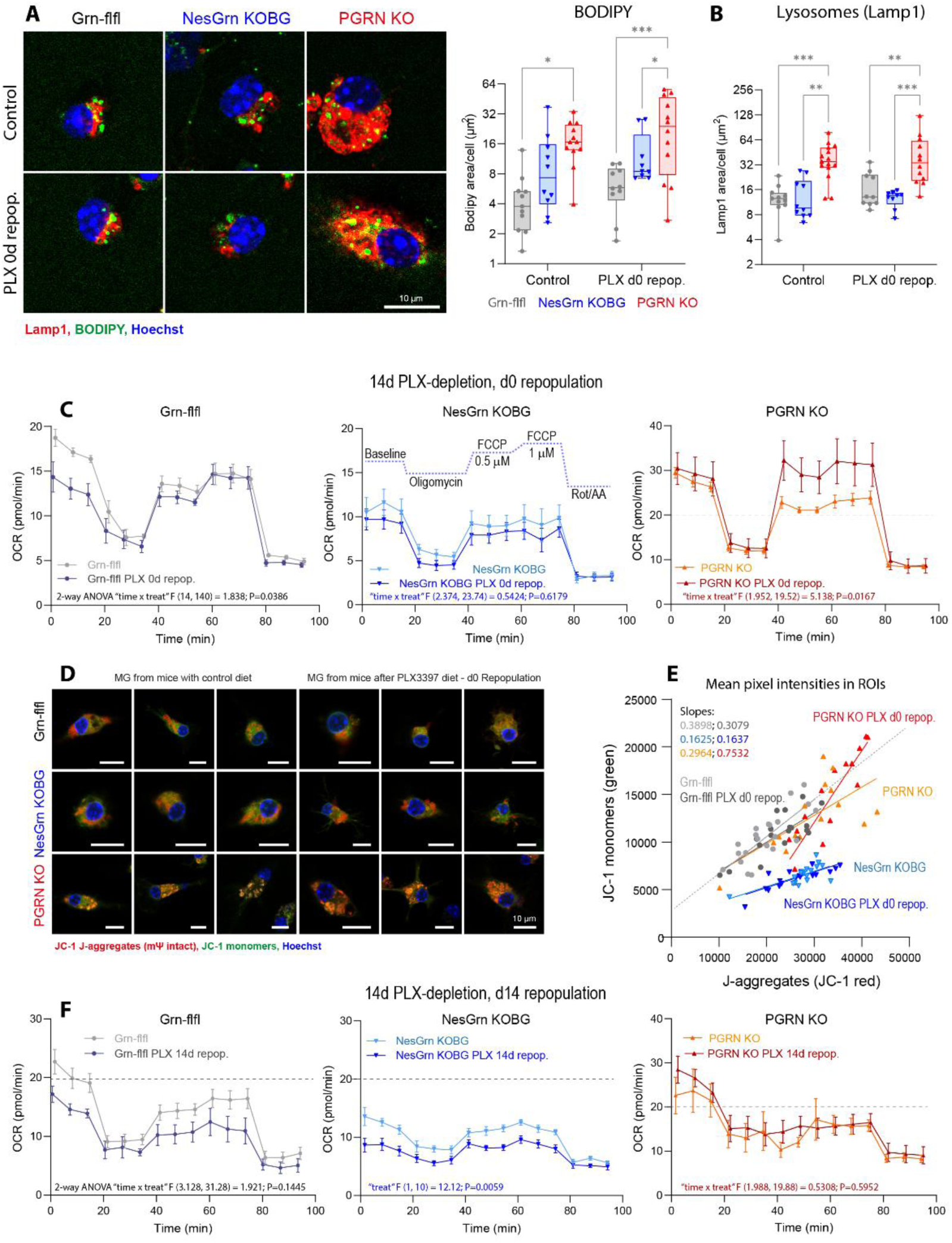
Microglia lipid load and mitochondrial respiratory functions. **A**: Exemplary immunofluorescent images of primary microglia of Grn-flfl control, NesGrn KOBG and PGRN KO mice at control conditions and after 14d of PLX3397 diet, which were prepared directly at the end of the depletion diet which is repopulation d0. Neutral lipids are stained with BODIPY and lysosomes with anti-Lamp1. **B:** The area of BODIPY and Lamp1 was used for quantification and submitted to one-way ANOVA and subsequent Dunnet posthoc analysis versus the control condition. The box is the interquartile range, the line is the median, scatters show the average total IF size per cell. Asterisks show significant difference versus control, *<0.05, **<0.01, ***<0.001. **C:** Seahorse analysis of mitochondrial oxygen consumption rate (OCR) in primary microglia cultures of n = 6 mice per genotype and treatment using a Mito-stress test (Agilent). Microglia cultures were prepared from Grn-flfl, NesGrn KOBG and PGRN KO mice without and after PLX3397 diet for 14 days, directly at the end of the diet at repopulation d0. After 3 cycles of baseline respiration, oligomycin 1.5 µM was injected to block ATP generation. Subsequently, FCCP was injected at 0.5 µM and then 1 µM to induce maximum oxygen consumption. Finally, Complex I and Complex III were inhibited by rotenone and antimycin A (Rot/AA) both at 0.5 µM to assess non mitochondrial residual respiration. Data were compared with 2-way ANOVA and subsequent Šidác posthoc analysis per time point. ANOVA results for “time x treatment” are shown in the graphs. Genotype comparisons show that OCR was high in PGRN KO microglia (Y-axis scale 0-40, reference line at 20; P<0.01) and further increased in PLX-resistant residual microglia (P<0.001), and low in NesGrn KOBG (P<0.05) and further decreased in in PLX-resistant residual microglia (P<0.01). **D:** JC-1 immunofluorescence images show each three exemplary microglia cells per genotype and treatment (time point repopulation d0 directly at the end of PLX-diet). JC-1 accumulates in mitochondria in a potential-dependent manner, forming red-fluorescent J-aggregates in polarized (high-potential) mitochondria, whereas depolarized mitochondria retain JC-1 in its monomeric, green-fluorescent form. DAPI was used as nuclear counterstain. **E:** Intensities of red J-aggregates were plotted against green JC-1 monomers. The slope of the regression line is an indicator of the mitochondrial membrane potential. Low slope means high potential. The potential is reduced in mitochondria of PGRN KO microglia after PLX treatment, whereas mitochondria of NesGrn KOBG microglia are over-polarized. **F:** Seahorse analysis of mitochondrial oxygen consumption rate (OCR) from microglia at 14 days of repopulation versus control conditions. Experiments and analysis as in C. OCR was significantly lower in microglia of NesGrn KOBG mice (P=0.09 control, 0.0008 after PLX) than Grn-flfl mice. No significant differences between Grn-flfl and PGRN KO microglia were observed.

Seahorse analysis of oxygen consumption rate (OCR) and quantification of JC-1 fluorescence area per cell revealed relatively low OCR in the microglia of NesGrn KOBG mice, as well as a reduction in both the red (J-aggregates) and green (monomeric) JC-1 signals, compared to all other conditions, both with and without PLX3397 treatment (see Figures 5C-F). This pattern suggests a global decrease in the amount of JC-1-accumulating mitochondrial membranes per cell, with the remaining mitochondria exhibiting a relatively preserved or increased membrane potential (high JC-1 red/green ratio). Consistent with this, the reduced OCR but normal relative response to stimuli indicates a reduction in mitochondrial mass with normal function. However, as JC-1 is potential-dependent and its green signal is not mitochondria-specific, these data do not permit a definitive conclusion regarding mitochondrial mass. In contrast to NesGrn KOBG microglia, OCR was increased at baseline and further increased upon FCCP stimulation in PGRN KO microglia, particularly in those obtained at the end of the PLX3397 diet (d0 repopulation) (Figure 5C). These PLX3397-resistant microglia exhibited a significant increase in the monomeric JC-1 signal (Figures 5D and 5E), indicating a disruption in membrane potential and mitochondrial functions, and an increased production of reactive oxygen species. Importantly, after 14 days of repopulation (Figure 5F), baseline OCR remained increased, but the extreme response to FCCP had resolved, indicating that microglia from PLX3397-treated PGRN KO mice had regained pre-PLX status.

Together, the BODIPY and mitochondrial data show that the positive resetting of microglia morphology observed following PLX3397 treatment did not readily translate into normalization of organelle function.

### PLX3397 treatment leads to escalation of excessive grooming and overactivity

Motivated by the successful renewal of microglia and restoration of synapses, we set out to investigate whether the rejuvenation of microglia was associated with an attenuation of pathological behavior in vivo. PGRN heterozygous mice were used in this experiment based on the assumption that some progranulin was essentially required to reset the microglia phenotype, and because haploinsufficiency resembles the genetics of human progranulin-associated FTD (heterozygous dominant negative). Multiple behavioral features were continuously monitored before, during, and after the PLX3397 or placebo diet in IntelliCages in old female NesGrn KOBG and PGRN het mice (Figures 6–7). Eight mice per genotype were randomly assigned to cages 1 (PLX) and 2 (placebo). As previously demonstrated, progranulin-deficient mice exhibited increased activity and compulsive behavior compared to historical age-matched controls (not shown). Overactivity manifested as a high number of ’useless’ visits without licks (NPVisits with nosepokes and SVisits without licks or nosepokes), whereas compulsiveness manifested as a high number of licks per visit (see Supplementary Figure S3). Interestingly, PGRN het mice were more overactive than NesGrn KOBG mice, despite showing higher overall brain progranulin expression (approximately 50 % versus 25 % of normal levels [31]. The overactivity in PGRN het mice was associated with a reduced circadian amplitude due to increased daytime activity.

**Figure 6.**
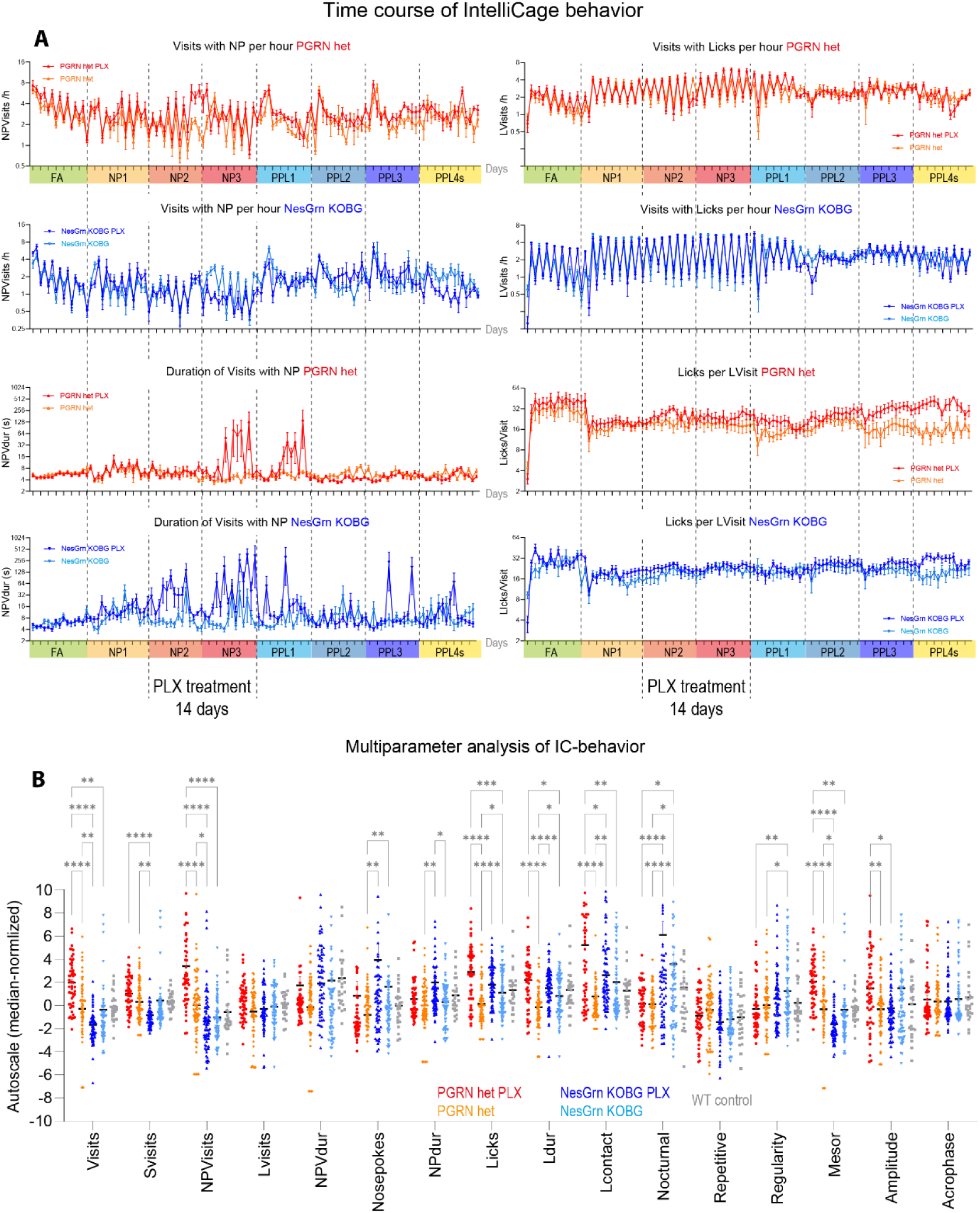
Overactivity and compulsive licking and disruption of NP behaviour during and after PLX. **A**: Time course of key behavioural parameters during different tasks in IntelliCages before, during and after treatment with PLX3397 diet versus control diet of NesGrn KOB and PGRN het mice. The tasks and the PLX-treatment period are described at the bottom of the graph and the periods shaded in different colours. The abbreviations of the tasks are FA, free adaptation; NP, nosepoke adaptation; PPL, place preference learning; PPLs (“s” for social challenge) where all mice of a cage are assigned to one rewarding corner. Further details about the tasks and IntelliCage abbreviations are shown in Suppl. Tables 2, 3. The data show means ± SEM of 8 female mice per group. The fluctuations of the behaviour show nighttime and daytime differences (12h Bins) and show the circadian rhythm. **B:** For statistical comparisons of multiple parameters behavioural features were averaged over each of the 8 tasks. Therefore, each mouse is represented by eight scatters showing the behaviour of the specific mouse in each task. The line is the average. Data were compared with 2-way ANOVA for the factors “IC-parameter” X “group” and posthoc comparison for “genotype” with adjustment of alpha according to Šidák. P*<0.05, **<0.01, ***<0.001, ****<0.0001.

**Figure 7.**
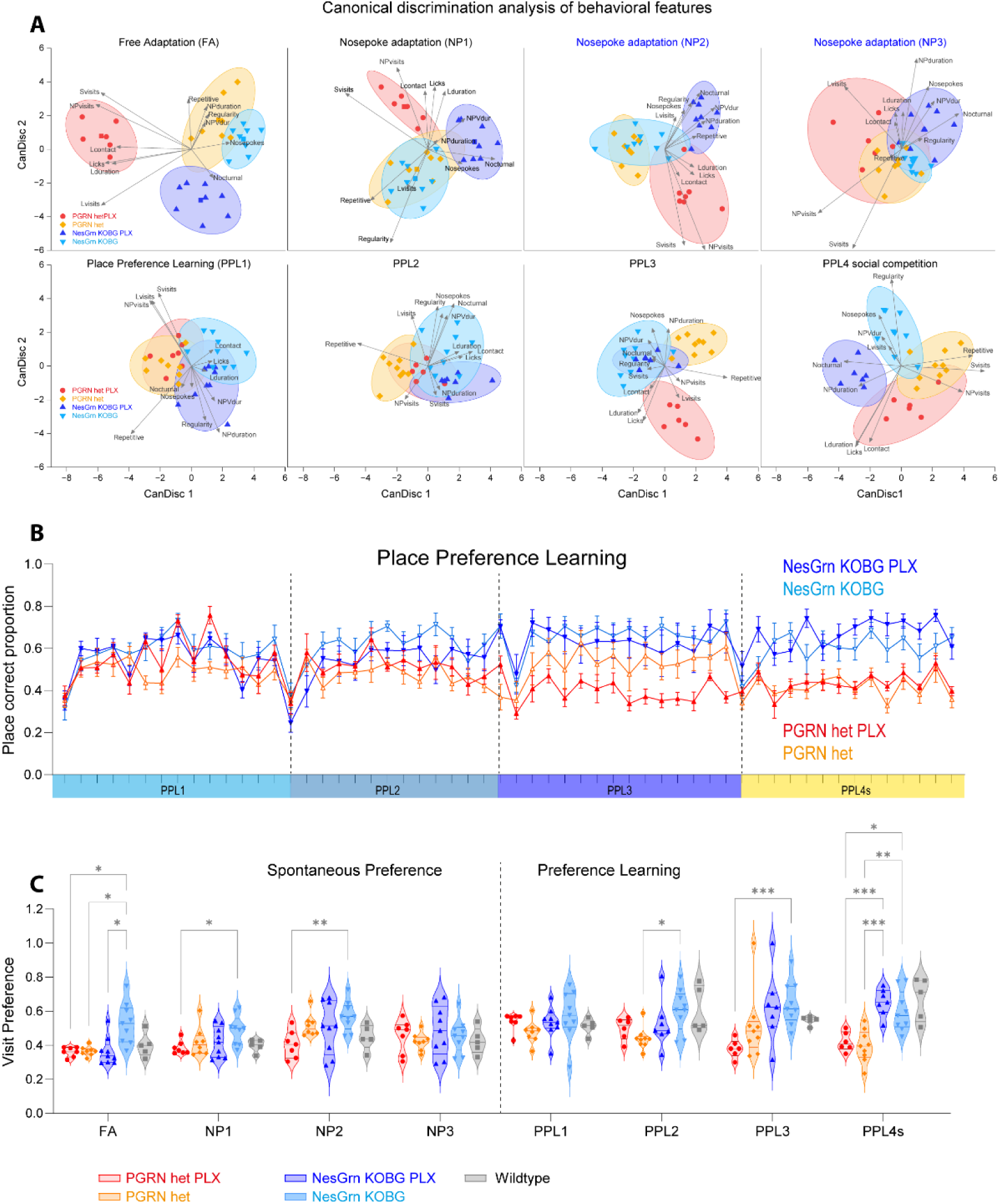
Impact of PLX treatment on genotype-dependent behaviour and learning ability. **A**: Canonical discrimination biplots using multiple behavioural parameters described in Fig. 6B as input. The ellipsoids show the 80% confidence interval for the group membership prediction. The treatment periods, NP2 and NP3, are highlighted in blue. Depending on tasks, differences between groups temporarily disappear but are reestablished in the final tasks. **B:** Time course of the accuracy of corner visits during place preference (PPL) tasks. While the proportion of correct corner visits is similar between genotypes initially, the curves diverge in later tasks because NesGrn KOBG with/without PLX steadily increase the accuracy whereas PGRN het mice do not learn to prefer a specific corner and the accuracy does not exceed the random level in the final PPL3 and PPL4s tasks. **C**: Violin/scatter plots and statistical comparison of corner visit accuracy (visit preference) as indicator of learning and memory. Each scatter is one mouse. Data were compared with 2-way ANOVA for the factors “task” X “group” and subsequent posthoc analysis using an adjustment of alpha according to Šidák. P*<0.05, **<0.01, ***<0.001.

Due to randomization, behavior in both cages was similar at baseline during free adaptation (FA) and the first week of nosepoke adaptation (NP1). The first week of the PLX3397 diet (NP2) was fairly uneventful, except for an increase in the duration of visits with nosepokes in NesGrn KOBG mice (Figure 6A and Supplementary Figure S3), accompanied by an increase in the number of NPVisits in the second PLX week. Similar behavioral changes occurred in PGRN het mice, albeit starting only in the second PLX week. In both genotypes, abnormal nosepoke behavior mostly resolved within seven to ten days after the end of the PLX diet, but compulsiveness (licks per visit) steadily increased in PLX-treated mice of both genotypes, whereas this parameter remained largely constant in placebo-fed mice. Compulsive licking was associated with compulsive grooming, resulting in serious skin lesions. Three mice (two NesGrn KOBG and one PGRN het) had to be euthanized during the first week of the PLX washout.

To statistically compare multiple behavioral features, the behavior over 24– or 12-hour intervals was averaged across the duration of the respective task. The averages were then pooled and presented as scatter plots in Figure 6B. Statistical comparisons confirmed an escalation in compulsiveness (Licks, Licking duration and Licking contact time) and overactive nosepoke behavior (nosepokes and NP duration) in PLX3397-treated mice compared to the control group. Summary behavior also showed stronger hyperactivity in PGRN het mice compared with NesGrn KOBG mice (Visits, Mesor).

To assess whether and how group membership could be predicted by behavior, the behavioral parameters were submitted to a Canonical Discrimination Analysis for each task (Figure 7A). The analysis showed that the PLX-treated groups of NesGrn KOBG and PGRN het mice were more divergent than their respective untreated groups, particularly in tasks where there was no need to memorize the rewarding corner. During the PPL tasks, the differences between the groups diminished, but reappeared in the social competition PPL4social task. During the final two PPL tasks (PPL3 and PPL4social), differences between genotypes became apparent in the accuracy of corner visits (Figures 7B and 7C). While the proportion of correct corner visits increased slightly over time in NesGrn KOBG mice, it remained at a random level in PGRN het mice, particularly in those that had received the PLX3397 diet. No such deterioration in cognitive behavior was observed in PLX-treated NesGrn KOBG mice.

### Thermal gradient ring excessive running after PLX3397

After completing the IntelliCage experiments, the same mice were observed in the TGR maze environment to obtain an unbiased view of their locomotion, activity, exploration, and grooming. Compared to historical control mice of the same age, all mice exhibited relatively flat temperature preference curves, but with a normal preference temperature of 28-30 °C (Figure 8A, B). The flatness of the curve is caused by overactivity, which prevents the mice from recognizing and settling into their preferred temperature zone. This overactivity manifests as long travel distances and high numbers of rotations (Figure 8C). The normal total distance traveled is 50–100 m with 50–100 changes in direction (rotations). While all animals were overactive, the PGRN het mice treated with PLX3397 showed excessive running, covering up to 1500 m in the 60-minute test. Consequently, the preference zone is nearly random, and the apparent preference temperature is low at 26 °C (Figures 8B and 8D). Figure 8D shows the interdependence of distance and preference temperature, for which data from each quarter were pooled. Some mice also exhibited excessive grooming, particularly in Q4 of the test, though there was no difference between PLX3397-treated and untreated mice (Figure 8E).

**Figure 8.**
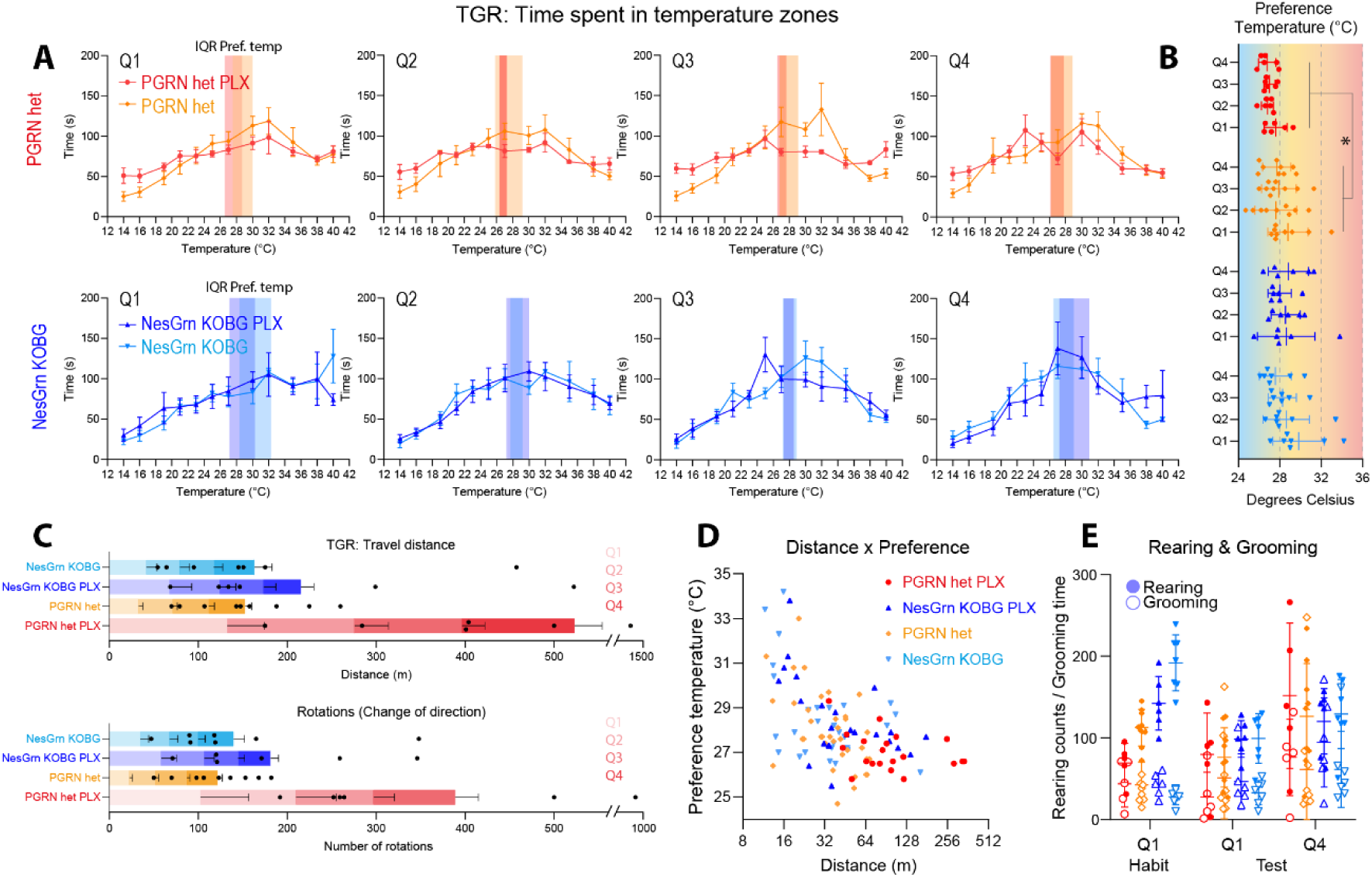
Motor and sensory behaviour in a Thermal Gradient Ring (TGR) maze. **A**: The same mice observed in IntelliCages in Figure 6-7 were subsequently subjected to further behavioural analyses in a TGR to assess locomotion, exploratory behaviour, grooming and rearing, and thermal preference. The test is split in 4 quarters for time dependent analysis (Q1-Q4). The graphs show the time spent in the 12 TGR temperature zones. The average preference temperature is highlighted as colour-shaded areas. The steepness of the preference curve increased from Q1-Q4 in NesGrn KOBG but not in PGRN KO mice, which were strongly overactive (C) and therefore did not develop a zone preference. **B:** Scatter plot showing the preference temperature over the 60-min test period. Each scatter is one mouse, the line is the mean, and whiskers show the SD. Data were analysed with one-way ANOVA and subsequent pairwise comparison of PLX treated versus non-treated groups. Asterisks show significant differences. *P<0.05. **C:** Stacked bar charts show the travel distances per quarter and body rotations per quarter. Body rotations lead to a change of clockwise or anti clockwise moving direction. **D:** XY-scatter plots show the inverse relationship between travel distance and preference temperature. Data pairs for each quarter were pooled. Hence, each mouse is represented with 4 scatters. **E:** Scatter plots with mean and SD (line and whiskers) of rearing and grooming behaviour in Q1 and Q4 of the habituation and TGR test. Rearing behaviour (filled triangles) predominates over grooming (open triangles) in Q1, particularly in NesGrn KOBG mice and shows normal initial exploration of the environment. In Q4 rearing and grooming are more balanced, but highly variable in PGRN het mice.

### Long lasting pro-inflammatory changes of brain transcriptome after PLX3397

To evaluate the long-term effects of PLX3397-induced microglia renewal on transcriptional and metabolic levels, the final brain tissues of IntelliCage mice and additional Grn-flfl mice, which were housed and treated simultaneously, underwent bulk mRNA sequencing and lipidome and metabolome analyses. Tissue was obtained eight weeks after the end of the PLX3397 diet to reflect the long-term outcome of microglia depletion and full repopulation. As we previously demonstrated genotype-dependent differential gene expression in our mice [31], we focused here on the impact of PLX3397-mediated depletion and repopulation.

Score plots from an unsupervised principal component analysis (PCA) of transcript expression (Figure 9A) show that genotypes are separable, but treatment groups (PLX3397 versus control diet) are not. Further supervised partial least squares-discrimination (PLS-DA) analyses, including ANOVA-significant genes, revealed clear differences between genotypes, yet there was no separation between PLX-treated and control-diet mice (Figure 9B). According to PLS-DA, NesGrn KOBG mice are unexpectedly more closely related to Grn-flfl mice than PGRN het mice, even though PGRN het mice have higher overall progranulin expression than NesGrn KOBG mice [31].

**Figure 9.**
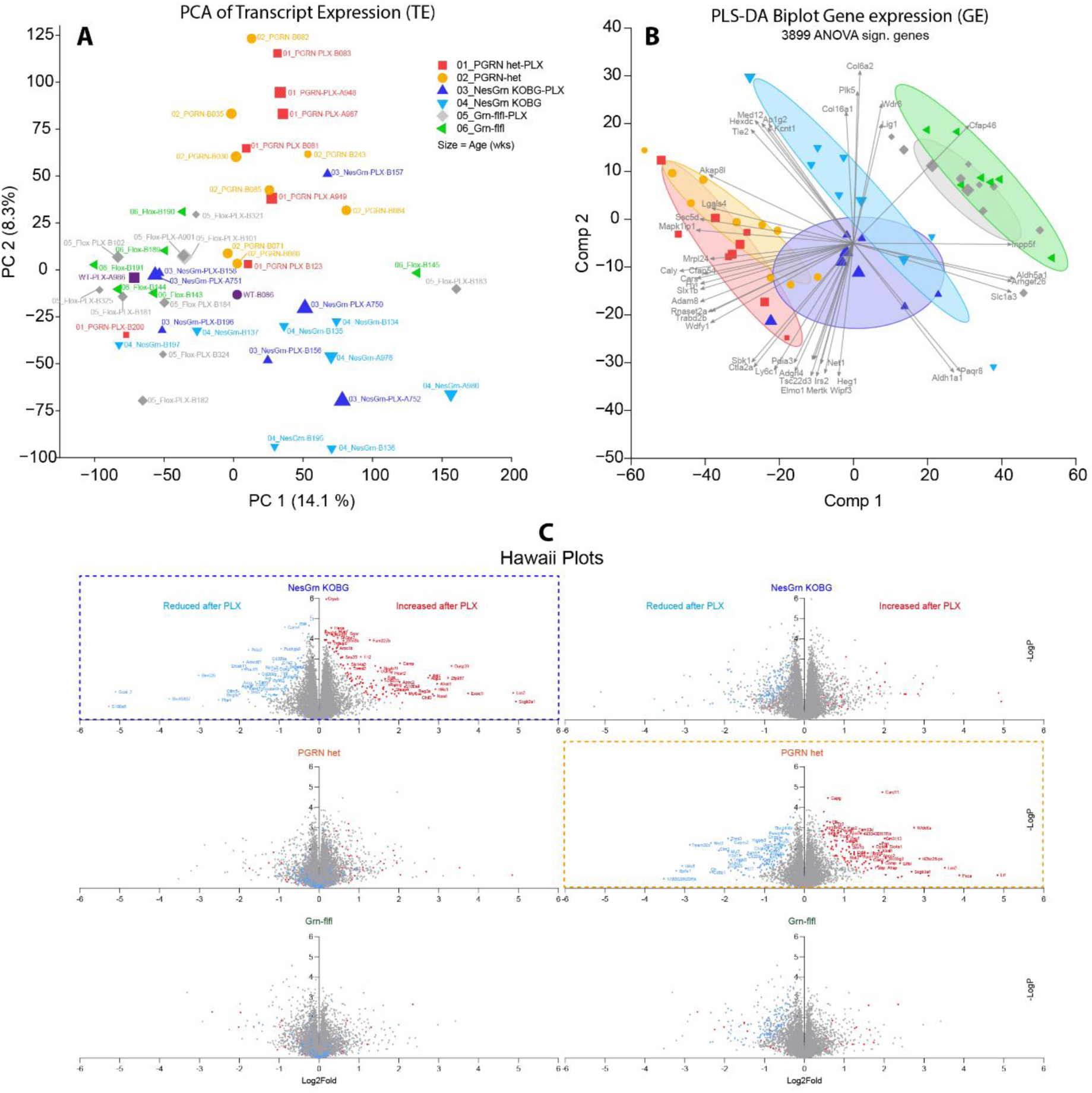
Brain transcriptome 8 weeks after PLX3397 treatment versus control. **A:** Principal component analysis (PCA) score plots of component –1 versus –2 of transcript expression in the brain of Grn-flfl, NesGrn KOBG and PGRN het mice including few PGRN KO mice. Mice were treated with a 2-week cycle of PLX3397 diet 8 weeks prior to tissue extraction and compared with mice who received normal control diet. Gene and transcript expression were analysed by 3’ mRNA sequencing. **B:** Group membership prediction and variable importance were further analysed by supervised partial least square discrimination (PLS-DA) analysis including all ANOVA significant genes (not adjusted P<0.05). The ellipsoids show the 90% confidence intervals for the prediction. The superimposed vector plot shows leading variables for PLS component-1 and –2. **C**: Hawaii plots of gene expression for each genotype for PLX-treated versus not-treated groups. The x-axis shows the Log2(Fold change) of normalized counts (CPM). The y-axis shows the negative logarithm of the t-test P-value. The data are obtained from n = 8 mice per group. In the left panel genes upregulated in NesGrn KOBG after PLX treatment are in red, downregulated genes in blue. The same genes are also colour coded in the respective PLX-vs-control diet plots of PGRN het and Grn-flfl mice. In the right panel, PGRN het mice are used as the reference genotype i.e. up-genes in red and down-genes in blue. The comparison of the Volcano plots shows that there is some agreement in PLX-dependent gene downregulation but little for upregulated genes.

We and others have shown that the genes that are upregulated in progranulin-deficient mice are mostly associated with microglia and the immune system [20, 31, 44, 60, 63, 64]. These genes include *Lyz2, Mpeg1, Gpnmb, Tyrobp, Cd68*, some complement factors (*C1q*’s), some cathepsins (*Cts*), *Trem2, Hexa, Hexb, Lgals3*, integrins, and immunoglobulin receptors. These genes are considered microglia marker genes in mice [65]. The RNA-seq results of the present study confirm genotype differences (key candidates are shown in the heatmap in Figure 9). In this study, we examined the impact of PLX3397 treatment by comparing PLX3397-treated and non-treated mice for each genotype.

The Hawaii plots (Figure 9C) show that the upregulated and downregulated genes are genotype-dependent and that the regulations were moderate, considering the product of log₂ fold change × log P-value. There were almost no lasting gene regulations in Grn-flfl mice. The genes that were up– or down-regulated in NesGrn KOB (left panel, framed plot) or in PGRN het mice (right panel, framed plot) after PLX3397 treatment do not align, indicating that they were regulated by PLX3397 treatment either in NesGrn KOB or in PGRN het, but rarely in both. Based on our histology results obtained early after the end of PLX3397 treatment, we expected a decrease in pro-inflammatory genes and an increase in neuronal structure and synaptic genes. However, this hypothesis was not true at the late time point of RNA sequencing. Instead, the genes that were consistently upregulated in NesGrn KOB and PGRN het after PLX3397 treatment were immune genes (Figure 10A and 10B). Some of these genes are key pro-inflammatory candidates that are characteristically associated with progranulin-deficient microglia (progranulin-DAMs), such as *Gpnmb, Lgals3, Tlr2*, and *S100* genes. Further analyses also showed an increase in cathepsins and complement factors, which are critical to synaptic pruning (Figures 10C and 10D). Although these genes were not among the top 100 regulated genes (Figure 10A Heatmap), they suggest that the intended attenuation of neuroinflammation was not achieved, but rather escalated. This is also confirmed by GO analyses of the commonly upregulated and downregulated genes (Figure 10E). Up-regulated genes after the PLX diet were involved in immune processes, while downregulated genes were involved in cell projection organization and development.

**Figure 10.**
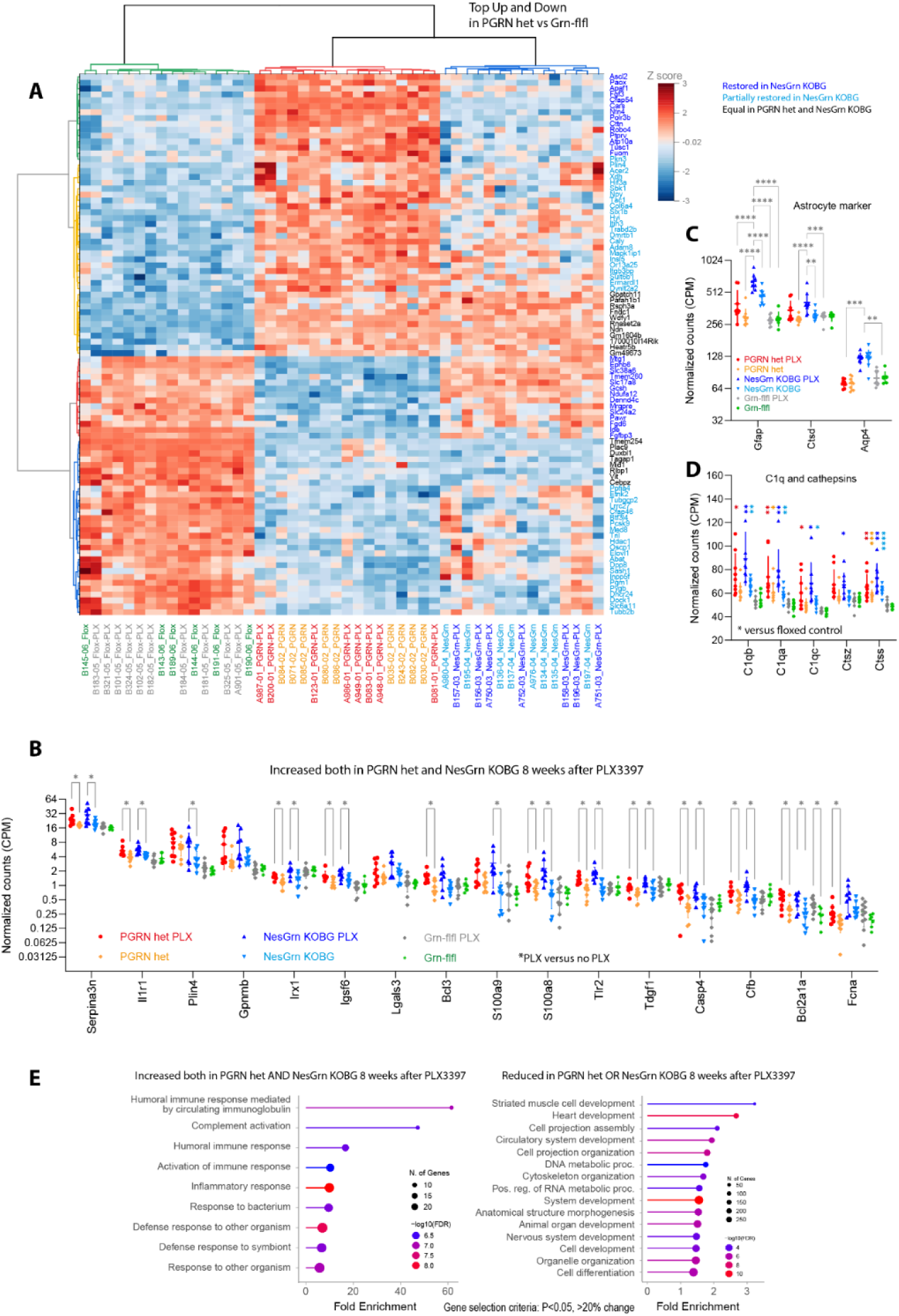
Long-lasting impact of PLX3397 treatment on brain gene expression. **A:** Heatmap of RNAseq top 100 regulated genes, selected by ANOVA P values, and hierarchical clustering of genes and mice. The tree shows Euclidean distance metrics, Ward method. Gene names in blue indicate difference between PGRN het and NesGrn KOBG mice. Major clusters differentiate Grn-flfl from progranulin deficient mice, the latter further clustered in PGRN het and NesGrn KOBG mice. PLX-treated and not treated mice were similar regarding the expression of the leading genotype-defining genes. **B, C, D:** Expression of candidate genes (mean ± SD of normalized counts) that were upregulated in PLX treatment groups of PGRN het and NesGrn KOBG mice. Data were submitted to 2-way ANOVA for “gene” x “group” and subsequent FDR adjusted posthoc analysis between groups. Significant results are marked with asterisks. In C, D Šidák posthoc analysis was used. P*<0.05, **<0.01, ***<0.001, ****<0.0001. **E:** Gene ontology enrichment analysis of genes significantly regulated in PLX-treated groups of NesGrn KOBG and PGRN het mice (criteria P<0.05, 20% change). The lollipop plots show the top significantly enriched GO BP terms. Upregulated genes are involved in inflammatory processed (left), downregulated genes in morphogenesis and development (right).

### No lasting impact of PLX3397 treatment on bulk brain lipidome

Progranulin-deficient microglia accumulate large amounts of neutral lipids, likely due to pathological scavenging and defective reutilization through lipophagy [21, 61, 62]. Plin4, a marker of lipid droplets, was strongly upregulated (Figure 10B). We confirmed the genotype-dependent accumulation of lipids in primary microglia of our mice (Figure 5). To determine whether PLX-mediated microglia renewal had a lasting impact on progranulin-associated lipid pathology, we subjected final brain tissue of the IntelliCage cohort mice and Grn-flfl controls to a lipidomic and metabolomic mass spectrometry analysis.

The results revealed genotype-dependent differences in four lipid classes (Figure 11A and 11B), partially confirming the results of previous studies [21, 31]. Specifically, there was a reduction in phosphatidylglycerols (PG), which partly represent the lysosome-specific group of lysobisphosphatidic acids (LBPA), also known as bis(monoacylglycero)phosphates (BMP) (e.g., PG 36:2 and PG 40:7), as well as an opposing increase in lysophosphatidylglycerols (LPG) (Figures 11A and 11B; Supplementary Figure S4). The PGs were the only lipid class that differentiated the PGRN het mice from the few PGRN knockout mice included in the analysis (indicated with arrows in Supplementary Figure S4B). There was also a decrease in some hexosylceramides (HexCer) and a strong decrease in long-chain acylcarnitines (Figure 11A, B; Supplementary Figure S5), confirming our previous studies [31]. Additionally, there was a strong reduction in cardiolipins (Figure 11A, B; Supplementary Figure S5), which are lipids of the inner mitochondrial membrane. Acylcarnitines transfer long-chain fatty acids through the inner mitochondrial membrane to supply beta oxidation. The reduction in both cardiolipins and acylcarnitines suggests a reduction in mitochondrial mass and function.

**Figure 11.**
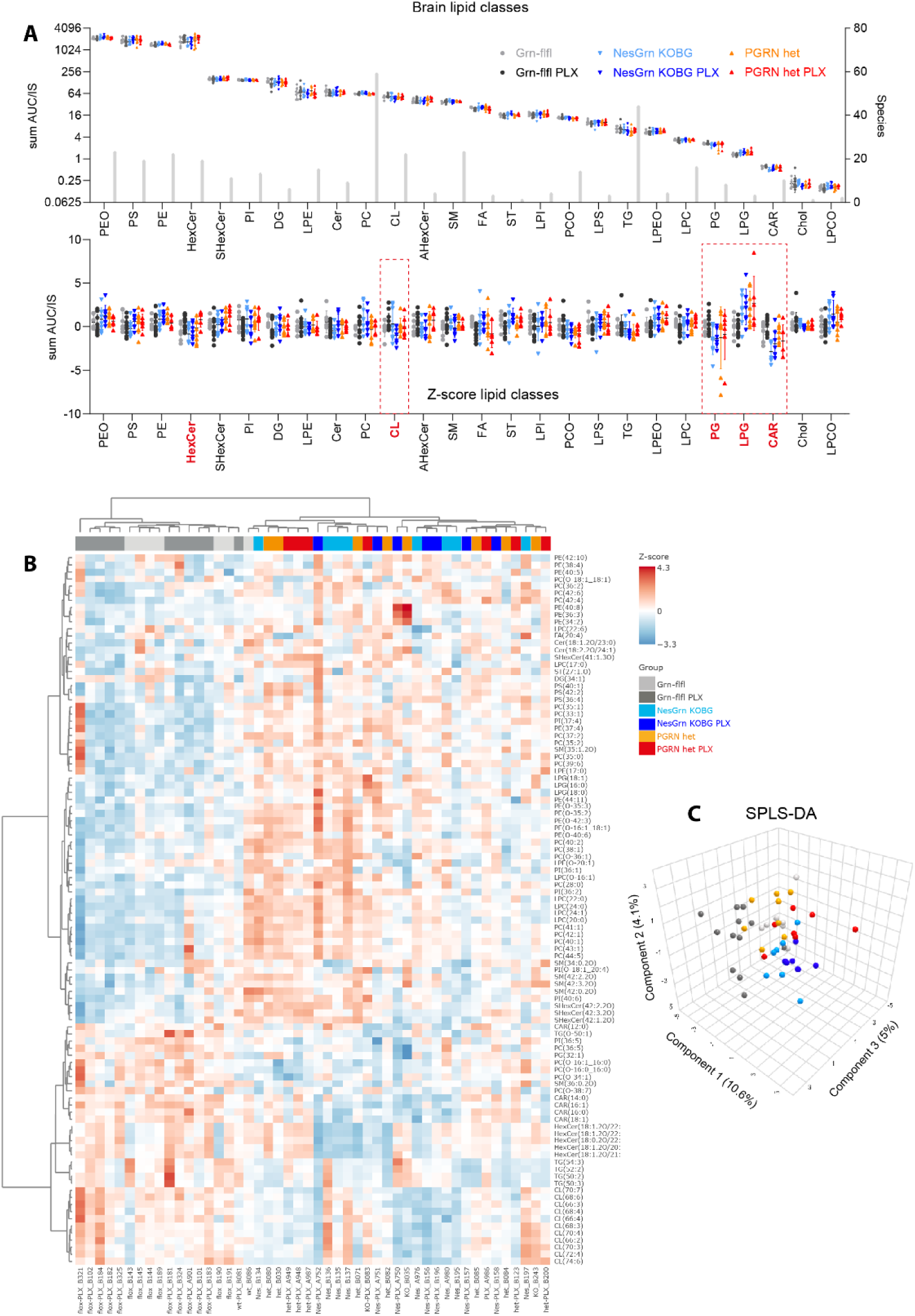
Brain lipidome 8 weeks after PLX3397 treatment versus control. **A**: Regulation of brain lipid classes sorted according to the abundance (high to low abundant), presented as summed AUCs (top) and Z-scores (bottom). The number of species per class is shown as grey bars and scaled on the right Y-axis. The Z-scores reveal a genotype dependent ANOVA-significant regulation of red framed lipid classes (PG, LPG, CAR, CL) and PLX-dependent regulation of HexCer and FA in Grn-flfl control mice. Volcano plots and individual lipid species of regulated classes are shown in Suppl. Figure S4 and S5. **B:** Heatmap of top 100 regulated lipids, selected by ANOVA P values, and hierarchical clustering of lipid species and mice. The tree shows Euclidean distance metrics, Ward method. Major clusters differentiate Grn-flfl from progranulin deficient mice. PLX-treated and not treated mice were similar. **C:** 3D scatter score plot of supervised sparse PLS-DA analysis. PLX treated Grn-flfl can be differentiated from the other groups owing to the differences in HexCer. Abbreviations: AHexCer, acetylhexosylceramides; CAR, <u>acylcarnitines</u>; Cer, ceramides; CE, cholesterol ester; CL, cardiolipins; DG, diglycerides; FA, fatty acids; HexCer, hexosylceramides; LPC, lysophosphatidylcholines; LPE, lysophosphatidylethanolamines; LPG, lysophosphatidylglycerols; LPI, lysophosphatidylinositols; LPS, lysophosphatidylserines; PC, phosphatidylcholines; PE, phosphatidylethanolamines; PG, phosphatidylglycerols; PI, phosphatidylinositols; PS, phosphatidylserines; SHexCer, sulfohexosylceramides; SM, sphingomyelins; ST, sterols; TG, triglycerides; –O ether bound lipids.

Indeed, JC-1 immunofluorescence studies of brain synaptosomes and isolated mitochondria show a reduction in JC-1 aggregates relative to monomers (JC-1 red/green ratio) in the absence of progranulin, indicating a loss of mitochondrial membrane potential. The synaptosomes of NesGrn KOBG mice were similar to the controls, which is consistent with the restoration of neuronal progranulin. However, the JC-1 ratio in isolated mitochondria was reduced (see Supplementary Figures S5A and S5B). PLX3397 treatment did not have a lasting impact on any genotype-associated lipid alterations (Figure 11 and Supplementary Figures S4 and S5), except for an increase in HexCer that occurred only in Grn-flfl controls (Supplementary Figures S4A and S4B). Since HexCer mainly originate from the neuronal/astrocytic fraction, the results suggest that astrocytes of Grn-flfl mice persist longer in ex-microglia niches than the astrocytes of progranulin-deficient mice because repopulation of microglia is slower in Grn-flfl mice. This finding is significant in light of human studies showing that loss of hexosylceramides (HexCer) in the brain and plasma correlates with the extent of axonal loss and disease duration in behavioral variant frontotemporal dementia (bvFTD) [66] and that high levels of HexCer in human plasma are associated with extreme longevity in humans [67]. Due in part to differences in HexCer levels, a sparse PLS-DA analysis distinguishes the Grn-flfl-PLX group from the others (Figure 10C).

### Microglial transcriptome

The bulk brain transcriptome and lipidome had revealed few lasting effects of the PLX3397 diet. However, since microglia only account for about 6% of the cells in the brain, subtle effects may be lost. Therefore, we isolated primary microglia and neurons/astrocytes from NesGrn KOBG mice at three time points after the PLX3397 diet: 0-day, 8-day, and 8-week repopulation versus the control diet (six mice per group) for RNA sequencing and lipidomic/metabolomic studies. We used NesGrn KOBG mice because their microglia are progranulin deficient, while their neurons have partially restored progranulin. This creates a condition similar to the currently favored clinical treatment approach of AAV9-mediated neuronal progranulin replacement.

Unsupervised principal component analysis (PCA) (Figure 12A and 12B) shows that the persisting microglia at the end of the PLX diet (day 0 repopulation) are distinct from the other groups, whose scatter clouds overlap. Neurons and astrocytes are mostly unaffected by the PLX diet, and all groups overlap. The late eight-week repopulation group is most distinct from the control group, suggesting that microglia renewal has a lasting impact on non-microglial cells. Microglia cluster analysis, including genes that were significantly regulated at day 0 repopulation, shows four major patterns: (i) an increase with overshooting normalization, (ii) an increase with slow normalization, (iii) a decrease with slow normalization, and (iv) a decrease with overshooting normalization (Figure 12C). Gene ontology analysis of the functions of the upregulated (a1, a2) and downregulated (b1, b2) genes shows that genes highly expressed in persistent microglia at d0 repopulation are involved in microtubule organization, cell projection assembly, and movement (Figure 12D), whereas genes low in persistent microglia are involved in the immune response and include key microglial marker genes, such as *CD74, CD86, ATP6V0, Mrc1, Ctss,* and *Fcgr3*. The top 200 regulated genes, according to ANOVA statistics, are shown in Supplementary Figure S6. An alluvial plot detailing the time courses in microglia and neurons/astrocytes is shown in Suppl. Figure S7. The data suggest that the PLX-resistant microglia, from which repopulation originates, are less proinflammatory than the populations lost during PLX treatment. Nonetheless, the repopulating microglia easily adopt the features of the eliminated population, with no obvious lasting benefit. Neuron/astrocyte genes that remain upregulated at eight weeks of repopulation (Figure 12E) are involved in nervous system development and protein metabolism (Figure 12F), suggesting ongoing reorganization to fill the microglial niche, which requires high protein turnover and cell migration.

**Figure 12.**
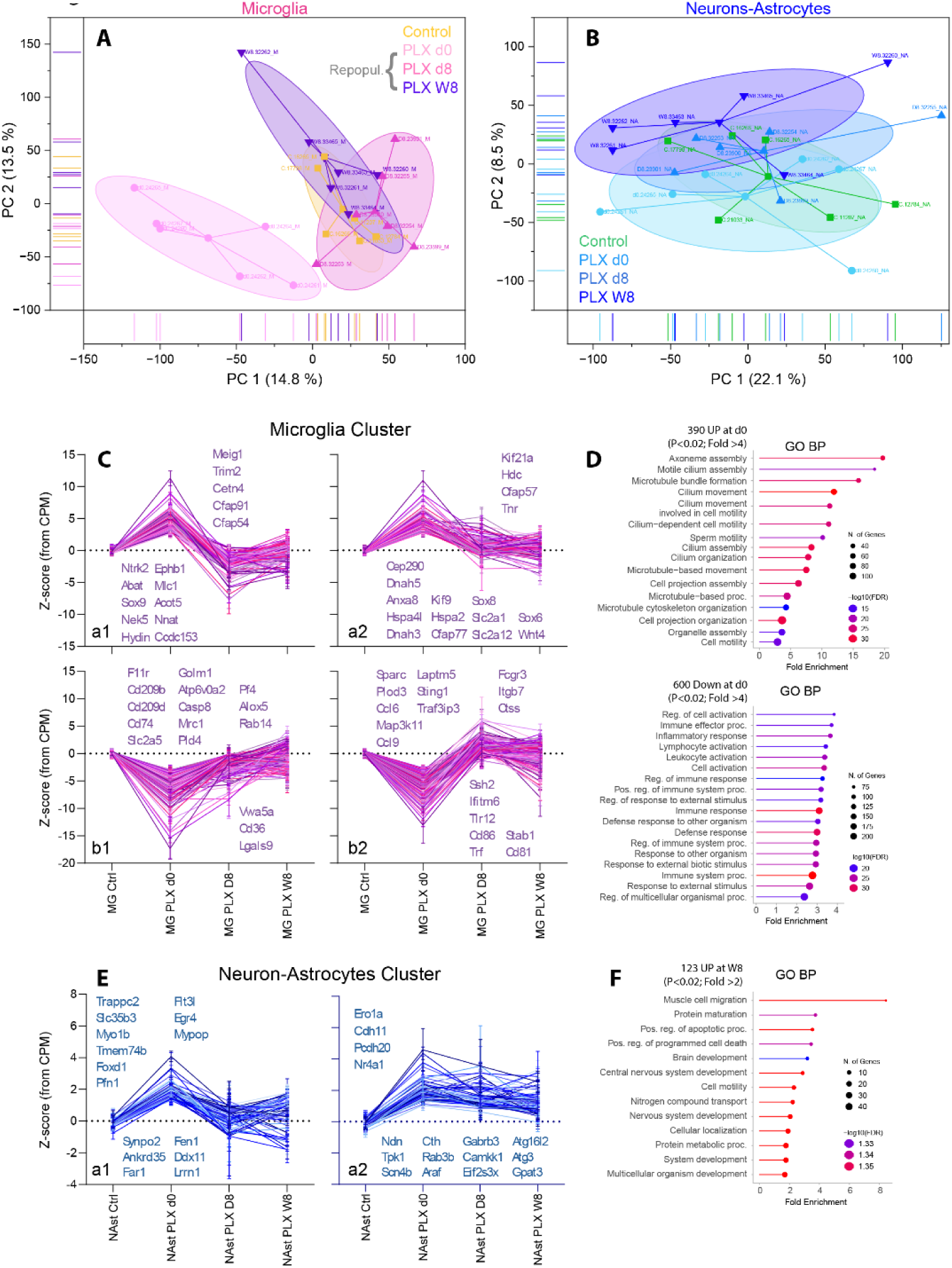
Microglial and astrocytic transcriptome during and after PLX3397 treatment of mice. **A, B**: Principal component analysis (PCA) score plots of RNAseq based gene expression in microglia and non-microglial cells (mostly neurons and astrocytes) before (control) and after treatment of mice with PLX3397-diet for 14 days. Tissue and cells were obtained from control mice and at the end of the depletion (day-0 repopulation) and 8 days and 8 weeks later (d8 repopulation, W8 repopulation). The ellipsoids show the 90% confidence interval for the prediction of group membership. The spots are labelled with the mouse IDs. For microglia, PLX d0 repopulation was distinct from the other groups. **C:** Time course of top up– and downregulated genes in microglia obtained at d0 repopulation (end of PLX). Genes follow 4 major patterns (a1, a2, b1, b2) (further details in Suppl. Figure S6, S7). Leading candidates are named. **D:** Gene ontology of biological processes (GO BP) shows that upregulated genes (cluster a1, a2) are involved in cell projection organization and movement, whereas downregulated genes (cluster b1, b2) are involved in immune responses. **E, F:** Fewer genes were regulated in neurons/astrocytes and showed two major up-regulation clusters (a1, a2). These genes are involved in protein metabolism and brain development.

### PLX3397 induced switch of fat load from microglia towards astroglia

Due to differences in cell and membrane size between neurons/astrocytes and the much smaller microglia, equal cell counts yielded different amounts of total lipids. Therefore, the lipids of the microglia were sum-normalized to harmonize with the total lipids of neurons/astrocytes and allow for a comparison of the lipid patterns between cell types. Analysis of lipid classes (Figure 13A) shows that phosphatidylglycerol (PG) and lysophosphatidylglycerol (LPG), which were affected by progranulin deficiency in the mouse brain (Figure 11A), predominantly originate from microglia. Hexosylceramides (HexCer), the only lipid class showing lasting effects of PLX3397, albeit only in Grn-flfl control mice, was mainly produced by neurons and astrocytes and temporarily increased after PLX3397 treatment.

**Figure 13.**
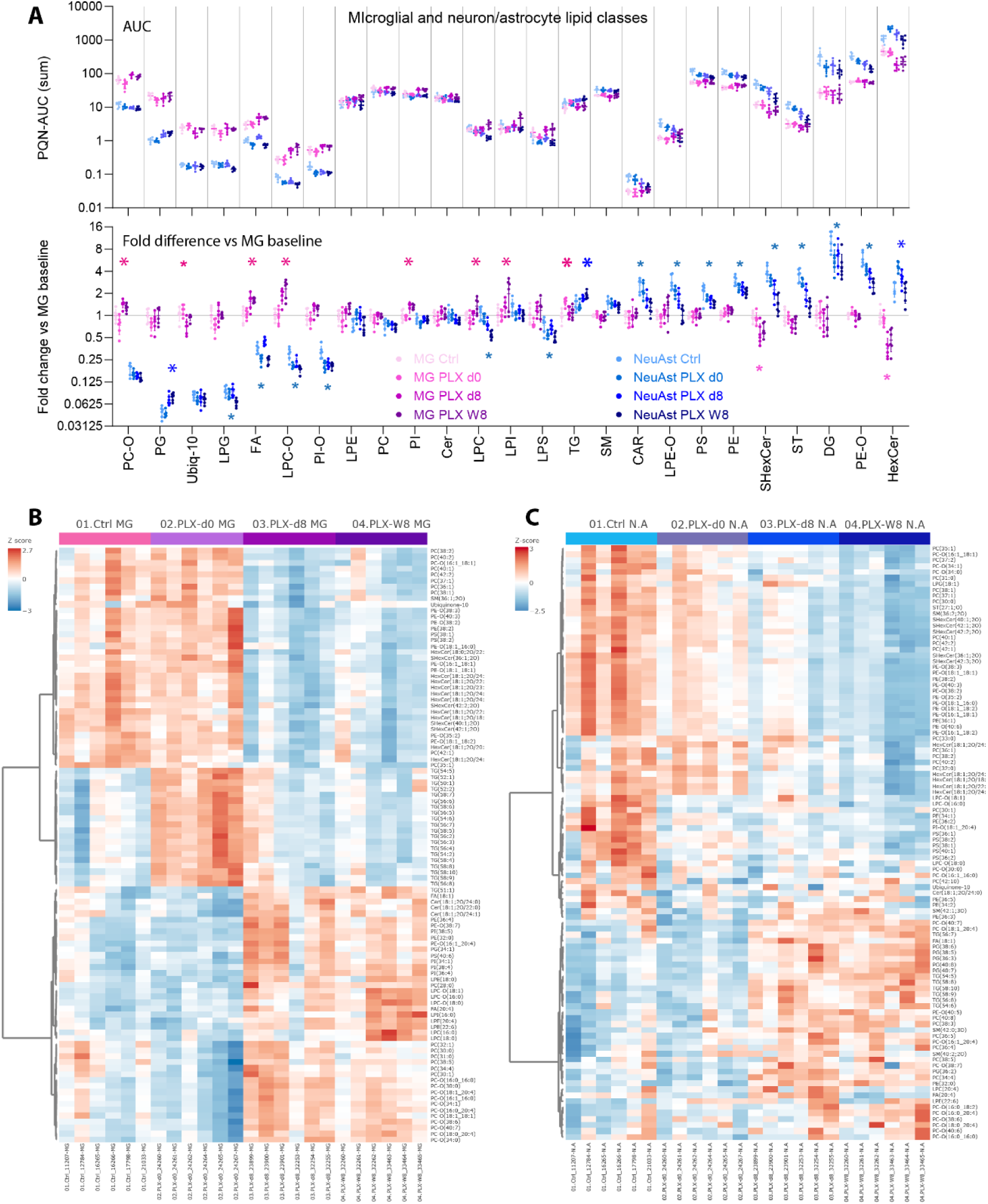
Microglia and astrocytic cell type specific untargeted lipidome during and after PLX3397. **A**: Regulation of lipid classes in primary microglia (MG) and primary neurons/astrocytes (NeuAst) sorted from left to right from high-to-low in MG to NeuAst ratios, presented as summed AUCs of lipid species (top) and fold change versus control microglia (bottom). Mice were treated in vivo with PLX3397 diet for 2 weeks and cells were isolated from NesGrn KOBG mouse brains at the end of PLX-diet (day-0 repopulation) and at day-8 (d8) and week-8 (W8) of the repopulation. Normalized data (fold change versus control MG) were submitted to 2-way ANOVA and subsequent FDR adjusted posthoc analysis, separately for MG and NeuAst. Large asterisks – pink for MG and blue for NeuAst – show lipid classes for which at least one time point after PLX was significantly increased, small asterisks show classes where at least one time point was decreased. Individual species of prominent regulated lipid classes are shown in Suppl. Figure S8. **B, C:** Heatmap and hierarchical clustering of top 100 regulated lipids in microglia (B) and in neurons/astrocytes (C), selected by ANOVA P values. The trees show Euclidean distance metrics, using the Ward method. For microglia, PLX d8 and W8 repopulation were similar but profoundly different from control MG. While repopulating MG got rid of their TG burden, NeuAst pick up the TG in the repopulation period (d8 and W8). Abbreviations: CAR, acylcarnitines; Cer, ceramides; CE, cholesterol ester; CL, cardiolipins; DG, diglycerides; FA, fatty acids; HexCer, hexosylceramides; LPC, lysophosphatidylcholines; LPE, lysophosphatidylethanolamines; LPG, lysophosphatidylglycerols; LPI, lysophosphatidylinositols; LPS, lysophosphatidylserines; PC, phosphatidylcholines; PE, phosphatidylethanolamines; PG, phosphatidylglycerols; PI, phosphatidylinositols; PS, phosphatidylserines; SHexCer, sulfohexosylceramides; SM, sphingomyelins; ST, sterols; TG, triglycerides; –O ether bound lipids.

In microglia, the PLX3397 diet caused an increase in ether-bound lipids (PC-O, LPC-O, and PI-O) and fatty acids (FA) during repopulation (days 8 and 8 weeks), whereas triacylglycerols (TGs), which cause the BODIPY immunofluorescence signal, were increased at the end of the PLX-diet (d0 repopulation), which agrees with the observed high BODIPY lipid load of the PLX-resistant population. Individual species in Suppl. Figure S8 show strong TG overload in microglia at d0-repopulation, particularly for long C-chain TGs. While TGs in microglia return to baseline levels during repopulation (Figure 13B and Supplementary Figure S8), TGs steadily increase in neurons and astrocytes (Figure 13C), suggesting that astrocytes scavenge some of the lipids released from microglia. While astrocytes get involved in handling the TG load, most other lipids, which are originally high in astrocytes, tend to decrease during repopulation, including CAR, PE, LPE, LPE-O, PS, SHexCer, and cholesterol (Figures 13A and 13C; Supplementary Figure S8). The data show that a single course of the PLX diet in vivo causes substantial changes in cell-type-dependent lipid profiles, which are not restricted to microglia. The metabolic change in neurons and astrocytes appears to be more profound than the transcriptional adaptation.

### Microglial rtPCR candidate genes

Upregulation of a panel of selected microglia relevant genes in isolated primary microglia was confirmed by rtPCR analysis also including PGRN het mice (not shown). Homeostatic and proinflammatory microglia marker genes were downregulated in microglia from PLX-treated mice in all genotypes directly at the end of the PLX-diet (d0 repopulation) but mostly returned to the baseline level during repopulation, some with an overshooting counter-regulation. The time to return depended on sex and genotype but the overall pattern was similar albeit with high interindividual variability not caused by age.

## Discussion

We have previously shown that restoration of progranulin only in neurons was insufficient to cure the FTD-like phenotype, resulting in behavioral deficits of NesGrn KOBG mice that were very similar to those of PGRN KO mice. Here we tested whether additional pharmacologic microglia renewal would improve the outcome. The rationale was that replacing old, neuron-aggressive microglia with newborn microglia would unmask positive effects of neuronal restoration because newborn microglia are believed to be less neuron-aggressive [32, 38, 68–70] and potentially more progranulin-responsive. To achieve this renewal, mice were fed a PLX3397 (pexidartinib) diet to deplete microglia, followed by spontaneous repopulation during washout. Although PLX3397 dieting temporarily restored synaptic spines, synapse density, and microglial morphology, it did not improve the behavioral FTD-like manifestations of hyperactivity and compulsiveness. Instead, it caused a temporary escalation of excessive grooming, accompanied by a lasting upregulation of several pro-inflammatory genes (e.g., complement, cathepsins). Progranulin-associated lipid deregulations in the brain remained unchanged after PLX treatment, but a loss of mitochondrial cardiolipins emerged. This loss was associated with abnormal mitochondrial respiration in primary microglia—excessive oxygen consumption in PGRN KO microglia and hyperpolarization in NesGrn KOBG microglia—both of which were intensified after PLX treatment.

Ex vivo cell type specific transcriptomic analyses further revealed that although pro-inflammatory genes were decreased directly at the end of the PLX diet in microglia, the effect was not maintained and was followed in part by an overshooting normalization. Neurons and astrocytes upregulated genes involved in brain development, which may reflect a transient regenerative response. However, lipid homeostasis in neurons and astrocytes was more strongly affected by PLX treatment than in microglia. While microglia mainly upregulated ether-bound lipids that influence membrane fluidity, these lipids were reduced in neurons and astrocytes [71]. Further classes decreased in neurons/astrocytes after PLX whereas phosphatidylglycerols (PG), triglycerides (TG), and HexCer (temporarily) increased. The latter is notable in light of a recent clinical study linking low peripheral and brain HexCer to the extent of neurodegeneration in FTD [66], while high plasma levels in another study were associated with extreme longevity in humans [67]. The increase of HexCer therefore points to a positive therapeutic effect. However, the observed increase in TG suggests that neurons and astrocytes may get involved in the accumulation of lipids when old lipid-laden microglia are replaced. The overload with neutral lipids is a process previously suggested to contribute to neurodegeneration [72, 73]. The profound impact on neurons and astrocytes indicates that microglia depletion—and particularly repopulation—has a strong indirect effect on the metabolism of neighboring cells.

Contrary to therapeutic expectations, compulsive grooming and restlessness escalated during and after PLX treatment, causing serious skin lesions in some animals that required euthanasia. PLX diet did not elicit skin scratching or lesions in Grn-flfl mice treated in parallel and did not affect all progranulin-deficient mice equally. Since progranulin is required for skin wound healing [74, 75] and CSF1R-signaling is crucial for the functioning of skin dendritic cells [76–78], it is plausible that small skin defects in aged mice fail to heal in the absence of progranulin and under concomitant depletion of skin dendritic cells, leading to itching, scratching, and progressive lesion formation. Once initiated, excessive grooming and local inflammation likely perpetuated each other. Importantly, deterioration of behavior and health occurred in both NesGrn KOBG and PGRN het mice, which retain about 30 % and 50 % of normal progranulin, respectively [31]. In PGRN het mice, this reduced amount is normally sufficient to prevent most brain pathology such as gliosis and lipofuscinosis [79], which characterize progranulin knockout mice [45], and should also support wound healing in the skin. PGRN het mice have so far been considered healthy except for subtle social deficits [16]. Likewise, although neuron-only restoration of progranulin did not prevent the FTD-like phenotype of NesGrn KOBG mice in our previous study [31], the mice were not seriously sick; compulsiveness was moderate and did not reduce lifespan. Behavioral studies and subsequent tissue analyses were performed in female mice only, due to the social constraints of IntelliCage housing. Progranulin deficiency appears to affect male and female mice similarly, but sex-dependent outcomes of PLX3397 treatment have been reported [80], with males mostly showing stronger benefits [80–82] which might be contributed by sex-differences in pharmacokinetic drug handling. In our mixed-sex molecular experiments, we observed some sex differences in PLX-induced gene expression but not in clinical or histologic effects. Nonetheless, it remains conceivable that male progranulin-deficient mice might experience stronger therapeutic benefits but also potentially stronger adverse effects. Taken together, PLX3397 treatment appeared to cause a serious deterioration even in mice with only partial progranulin deficiency, suggesting that therapeutic microglia depletion–repopulation might pose substantial risks in clinical conditions with progranulin-deficiency.

Because we could not observe all four genotype colonies with and without PLX3397 in parallel, we compared the IntelliCage results of the present cohorts with historical PGRN KO and Grn-flfl cohorts. Hyperactivity and compulsive over-licking behavior were reproducible across all progranulin-deficient cohorts [31, 47, 83]. PGRN het mice in the present study also showed this phenotype. However, paradoxically and unlike PGRN KO mice, PGRN het mice did not develop place preference behavior. Their corner visit accuracy remained at random levels, particularly in the PLX-treated group. In contrast, NesGrn KOBG mice did not show impaired preference learning, although their nosepoke behavior during and shortly after PLX diet was more disrupted than in PGRN het mice. Unexpectedly, exploratory and temperature preference behavior of NesGrn KOBG mice with or without PLX treatment was closer to normal in the Thermal Gradient Ring, whereas PLX-treated PGRN het mice were extremely overactive. Thus, although PGRN het mice produce more progranulin than NesGrn KOBG mice [31], they did not have a better outcome. This suggests that behavioral vulnerability depends on the neuronal subpopulations and regions in which progranulin is preserved or restored. Spatial transcriptomics may clarify this in future studies. In our previous snRNAseq studies, progranulin was restored in about two-thirds of normally positive neurons in NesGrn KOBG mice, distributed across excitatory and inhibitory neurons [31]. It is unknown whether progranulin expression in PGRN het mice is equally distributed or restricted to specific subtypes or regions, which might explain their unexpected vulnerability to PLX3397 treatment.

For functional and morphological studies of primary microglia, we mostly used PGRN KO microglia as comparators because NesGrn KOBG microglia are also progranulin-deficient and should theoretically resemble PGRN KO microglia if morphology and function depended solely on intrinsic progranulin expression. Differences between the two would therefore reflect environmental influences in vivo. A limitation is that PGRN het microglia were not included in all experiments. Morphologically, NesGrn KOBG, PGRN KO, and PGRN het microglia were similar and responded equally to PLX3397-mediated depletion and early repopulation, which manifested in a loss of a phagocyte-like subpopulation, previously described as *Gpnmb*, *Mertk*, *Lgals3*, *Apobec*, and/or *ATP6v0d2* positive [31, 60]. However, despite similar morphology, PGRN KO microglia had a higher lipid load and very high baseline, stimulated and residual oxygen consumption, whereas OCR was low in NesGrn KOBG microglia. This pattern persisted and was accentuated after PLX diet and was reproducible across timely distant mouse cohorts, clearly showing that PGRN KO microglia were metabolically hyperactive. A similar microglial OXPHOS hyperactivity was previously described in induced human microglia from patients carrying TSC (tuberous sclerosis complex) mutations in association with upregulations of genes involved in lipid metabolism, innate immunity and phagocytosis [84]. Although genetically distinct, the pathology of progranulin and TSC deficiency is linked via defective lysosomal functions [85]. Mitochondria in microglia form non-fusogenic physical contact site with lysosomes, promoting signal transduction and modulation of inflammatory phenotypes [86]. JC-1 analysis revealed a loss of mitochondrial membrane potential in PGRN KO microglia directly after PLX diet in line with previous studies [87], whereas NesGrn KOBG microglia were hyperpolarized with or without PLX, explaining their low OCR. These results suggest that progranulin in the surrounding brain environment shapes microglial oxidative capacity, an effect maintained ex vivo. Since oxidative burst capacity influences both pathogen defense and neuronal attack, PGRN KO microglia may be simultaneously more protective and more destructive than NesGrn KOBG microglia.

Our findings have potential therapeutic implications and add to the understanding of cell type–specific functions of PGRN. While neuronal progranulin restoration alone was insufficient to cure the phenotype, the present results show that microglia depletion and repopulation may carry substantial risks and cause temporary deterioration. It remains to be seen whether the adverse effects observed in mice predict outcomes in humans once more patients with dementia receive pexidartinib or similar drugs. Current trials are restricted to Alzheimer’s disease, which may be less vulnerable to CSF1R inhibitor effects [88, 89]. For progranulin-FTD, combining microglia depletion with replacement by genome-edited myeloid cells or allogeneic stem cell transplantation may be necessary, an approach that showed promising results in a progranulin-knockout mouse model [44, 90, 91] and in rare cases of inherited human leukodystrophy [92, 93]. Combining microglia renewal with brain-penetrant progranulin analogues or uptake-enhancing fusion constructs [20, 21] may offer another strategy to simultaneously renew and reset microglia and rescue neurons.

## Supporting information

Supplemental Figures with legends

Supplemental Analytical Procedures

## Acknowledgements

The authors would like to thank Rahmat Mojaradfar for technical support in performing the LC-HRMS and LC-MS/MS experiments.

## Data availability statement

All data that were analysed for the study are presented within the manuscript or supplementary files. The RNAseq data have been uploaded to the GEO repository and are available under the accession number GSE287561 (mouse brain RNAseq) and GSE324813 (RNAseq microglia and neuron/astrocytes). The submissions are still private until publication of this manuscript.

Lipidomic studies of mouse brain are available at BioStudies under the accession number S-BSST2143. Lipidomic studies of microglia and neurons/astrocytes are are available at BioStudies under the accession number S-BSST2921.

The data is still private and will be released upon acceptance of the manuscript.

## Author contributions

MPW performed behavioural experiments in mice, collected tissue and blood samples, performed RNA, primary microglia and histology studies, analysed data, created tables and histology figures and drafted parts of the manuscript; LH established, performed and supervised untargeted lipidomic analyses; LF contributed to preparation of primary cultures; CA and YS performed lipidomic analyses; MKES discussed the study design and analysis, contributed to immunofluorescence studies, and edited the manuscript; IT conceived and designed the study, acquired funding and ethical approval, analysed IntelliCage and RNAseq and Seahorse data, compiled and analysed lipidomic and metabolomic data, created figures and wrote the paper (draft and editing). All authors contributed to writing or editing parts of the manuscript and agree to the latest version of the manuscript.

## Ethical Approval

Mouse behavioural studies were approved by the local Ethics Committee for animal research (Darmstadt, Germany) under FK1103 and FU2080. For primary cultures and tissue analyses of naive mice, tissue was obtained after euthanization according to $4 of the German Tierschutzgesetz in compliance with the ARRIVE principles [94], and with European guidelines and GV-SOLAS recommendations for animal welfare in science.

## Abbreviations

See table of abbreviations

## References

1. Petkau TL, Neal SJ, Orban PC, MacDonald JL, Hill AM, Lu G, Feldman HH, Mackenzie IR, Leavitt BR: Progranulin expression in the developing and adult murine brain. J Comp Neurol 2010, 518:3931–3947.

2. Neill T, Buraschi S, Goyal A, Sharpe C, Natkanski E, Schaefer L, Morrione A, Iozzo RV: EphA2 is a functional receptor for the growth factor progranulin. J Cell Biol 2016, 215:687–703.

3. Altmann C, Vasic V, Hardt S, Heidler J, Haussler A, Wittig I, Schmidt MH, Tegeder I: Progranulin promotes peripheral nerve regeneration and reinnervation: role of notch signaling. Mol Neurodegener 2016, 11:69.

4. Tang W, Lu Y, Tian QY, Zhang Y, Guo FJ, Liu GY, Syed NM, Lai Y, Lin EA, Kong L, et al: The growth factor progranulin binds to TNF receptors and is therapeutic against inflammatory arthritis in mice. Science 2011, 332:478–484.

5. Hu F, Padukkavidana T, Vaegter CB, Brady OA, Zheng Y, Mackenzie IR, Feldman HH, Nykjaer A, Strittmatter SM: Sortilin-mediated endocytosis determines levels of the frontotemporal dementia protein, progranulin. Neuron 2010, 68:654–667.

6. Davis SE, Roth JR, Aljabi Q, Hakim AR, Savell KE, Day JJ, Arrant AE: Delivering progranulin to neuronal lysosomes protects against excitotoxicity. J Biol Chem 2021, 297:100993.

7. Altmann C, Hardt S, Fischer C, Heidler J, Lim HY, Haussler A, Albuquerque B, Zimmer B, Moser C, Behrends C, et al: Progranulin overexpression in sensory neurons attenuates neuropathic pain in mice: Role of autophagy. Neurobiol Dis 2016, 96:294–311.

8. Du H, Zhou X, Feng T, Hu F: Regulation of lysosomal trafficking of progranulin by sortilin and prosaposin. Brain Commun 2022, 4:fcab310.

9. Baker M, Mackenzie IR, Pickering-Brown SM, Gass J, Rademakers R, Lindholm C, Snowden J, Adamson J, Sadovnick AD, Rollinson S, et al: Mutations in progranulin cause tau-negative frontotemporal dementia linked to chromosome 17. Nature 2006, 442:916–919.

10. Gass J, Cannon A, Mackenzie IR, Boeve B, Baker M, Adamson J, Crook R, Melquist S, Kuntz K, Petersen R, et al: Mutations in progranulin are a major cause of ubiquitin-positive frontotemporal lobar degeneration. Hum Mol Genet 2006, 15:2988–3001.

11. Cruts M, Gijselinck I, van der Zee J, Engelborghs S, Wils H, Pirici D, Rademakers R, Vandenberghe R, Dermaut B, Martin JJ, et al: Null mutations in progranulin cause ubiquitin-positive frontotemporal dementia linked to chromosome 17q21. Nature 2006, 442:920–924.

12. van Swieten JC, Heutink P: Mutations in progranulin (GRN) within the spectrum of clinical and pathological phenotypes of frontotemporal dementia. Lancet Neurol 2008, 7:965–974.

13. Feng T, Minevich G, Liu P, Qin HX, Wozniak G, Pham J, Pham K, Korgaonkar A, Kurnellas M, Defranoux NA, et al: AAV-GRN partially corrects motor deficits and ALS/FTLD-related pathology in Tmem106b(-/-)Grn(-/-) mice. iScience 2023, 26:107247.

14. Kurnellas M, Mitra A, Schwabe T, Paul R, Arrant AE, Roberson ED, Ward M, Yeh F, Long H, Rosenthal A: Latozinemab, a novel progranulin-elevating therapy for frontotemporal dementia. J Transl Med 2023, 21:387.

15. Kashyap SN, Fox SN, Wilson KI, Murchison CF, Ambaw YA, Walther TC, Farese RV, Arrant AE, Roberson ED: Carboxy-terminal blockade of sortilin binding enhances progranulin gene therapy, a potential treatment for frontotemporal dementia. bioRxiv 2024.

16. Arrant AE, Filiano AJ, Unger DE, Young AH, Roberson ED: Restoring neuronal progranulin reverses deficits in a mouse model of frontotemporal dementia. Brain 2017, 140:1447–1465.

17. Arrant AE, Onyilo VC, Unger DE, Roberson ED: Progranulin Gene Therapy Improves Lysosomal Dysfunction and Microglial Pathology Associated with Frontotemporal Dementia and Neuronal Ceroid Lipofuscinosis. J Neurosci 2018, 38:2341–2358.

18. Doyle JJ, Maios C, Vrancx C, Duhaime S, Chitramuthu B, Bennett HPJ, Bateman A, Parker JA: Chemical and genetic rescue of in vivo progranulin-deficient lysosomal and autophagic defects. Proc Natl Acad Sci U S A 2021, 118.

19. Hinderer C, Miller R, Dyer C, Johansson J, Bell P, Buza E, Wilson JM: Adeno-associated virus serotype 1-based gene therapy for FTD caused by GRN mutations. Ann Clin Transl Neurol 2020, 7:1843–1853.

20. Reich M, Simon MJ, Polke B, Paris I, Werner G, Schrader C, Spieth L, Davis SS, Robinson S, de Melo GL, et al: Peripheral expression of brain-penetrant progranulin rescues pathologies in mouse models of frontotemporal lobar degeneration. Sci Transl Med 2024, 16:eadj7308.

21. Logan T, Simon MJ, Rana A, Cherf GM, Srivastava A, Davis SS, Low RLY, Chiu CL, Fang M, Huang F, et al: Rescue of a lysosomal storage disorder caused by Grn loss of function with a brain penetrant progranulin biologic. Cell 2021, 184:4651–4668.e4625.

22. Kashyap SN, Boyle NR, Roberson ED: Preclinical Interventions in Mouse Models of Frontotemporal Dementia Due to Progranulin Mutations. Neurotherapeutics 2023, 20:140–153.

23. Ward M, Carter LP, Huang JY, Maslyar D, Budda B, Paul R, Rosenthal A: Phase 1 study of latozinemab in progranulin-associated frontotemporal dementia. Alzheimers Dement (N Y) 2024, 10:e12452.

24. Sevigny J, Uspenskaya O, Heckman LD, Wong LC, Hatch DA, Tewari A, Vandenberghe R, Irwin DJ, Saracino D, Le Ber I, et al: Progranulin AAV gene therapy for frontotemporal dementia: translational studies and phase 1/2 trial interim results. Nat Med 2024, 30:1406–1415.

25. Amado DA, Rieders JM, Diatta F, Hernandez-Con P, Singer A, Mak JT, Zhang J, Lancaster E, Davidson BL, Chen-Plotkin AS: AAV-Mediated Progranulin Delivery to a Mouse Model of Progranulin Deficiency Causes T Cell-Mediated Toxicity. Mol Ther 2019, 27:465–478.

26. Wang S, Weyer MP, Hummel R, Wilken-Schmitz A, Tegeder I, Schäfer MKE: Selective neuronal expression of progranulin is sufficient to provide neuroprotective and anti-inflammatory effects after traumatic brain injury. J Neuroinflammation 2024, 21:257.

27. Vasek MJ, Garber C, Dorsey D, Durrant DM, Bollman B, Soung A, Yu J, Perez-Torres C, Frouin A, Wilton DK, et al: A complement-microglial axis drives synapse loss during virus-induced memory impairment. Nature 2016, 534:538–543.

28. Sellgren CM, Gracias J, Watmuff B, Biag JD, Thanos JM, Whittredge PB, Fu T, Worringer K, Brown HE, Wang J, et al: Increased synapse elimination by microglia in schizophrenia patient-derived models of synaptic pruning. Nat Neurosci 2019, 22:374–385.

29. Kurematsu C, Sawada M, Ohmuraya M, Tanaka M, Kuboyama K, Ogino T, Matsumoto M, Oishi H, Inada H, Ishido Y, et al: Synaptic pruning of murine adult-born neurons by microglia depends on phosphatidylserine. J Exp Med 2022, 219.

30. Lui H, Zhang J, Makinson SR, Cahill MK, Kelley KW, Huang HY, Shang Y, Oldham MC, Martens LH, Gao F, et al: Progranulin Deficiency Promotes Circuit-Specific Synaptic Pruning by Microglia via Complement Activation. Cell 2016, 165:921–935.

31. Weyer MP, Hahnefeld L, Franck L, Angioni C, Klein M, Geisslinger G, Schäfer MKE, Tegeder I: Selective neuronal restoration of progranulin does not prevent the frontotemporal dementia like-phenotype of progranulin knockout mice. J Neuroinflammation 2026, 23:34.

32. Huang Y, Xu Z, Xiong S, Sun F, Qin G, Hu G, Wang J, Zhao L, Liang YX, Wu T, et al: Repopulated microglia are solely derived from the proliferation of residual microglia after acute depletion. Nat Neurosci 2018, 21:530–540.

33. Zhan L, Krabbe G, Du F, Jones I, Reichert MC, Telpoukhovskaia M, Kodama L, Wang C, Cho SH, Sayed F, et al: Proximal recolonization by self-renewing microglia re-establishes microglial homeostasis in the adult mouse brain. PLoS Biol 2019, 17:e3000134.

34. Willis EF, MacDonald KPA, Nguyen QH, Garrido AL, Gillespie ER, Harley SBR, Bartlett PF, Schroder WA, Yates AG, Anthony DC, et al: Repopulating Microglia Promote Brain Repair in an IL-6-Dependent Manner. Cell 2020, 180:833–846.e816.

35. Sosna J, Philipp S, Albay R, 3rd, Reyes-Ruiz JM, Baglietto-Vargas D, LaFerla FM, Glabe CG: Early long-term administration of the CSF1R inhibitor PLX3397 ablates microglia and reduces accumulation of intraneuronal amyloid, neuritic plaque deposition and pre-fibrillar oligomers in 5XFAD mouse model of Alzheimer’s disease. Mol Neurodegener 2018, 13:11.

36. Crapser JD, Ochaba J, Soni N, Reidling JC, Thompson LM, Green KN: Microglial depletion prevents extracellular matrix changes and striatal volume reduction in a model of Huntington’s disease. Brain 2020, 143:266–288.

37. Neal ML, Fleming SM, Budge KM, Boyle AM, Kim C, Alam G, Beier EE, Wu LJ, Richardson JR: Pharmacological inhibition of CSF1R by GW2580 reduces microglial proliferation and is protective against neuroinflammation and dopaminergic neurodegeneration. FASEB J 2020, 34:1679–1694.

38. Spangenberg E, Severson PL, Hohsfield LA, Crapser J, Zhang J, Burton EA, Zhang Y, Spevak W, Lin J, Phan NY, et al: Sustained microglial depletion with CSF1R inhibitor impairs parenchymal plaque development in an Alzheimer’s disease model. Nat Commun 2019, 10:3758.

39. Wang S, Wang Y, Strehle J, Wernersbach I, Papakonstantinou E, Somnuke P, Ritter K, Klein M, Tegeder I, Schäfer MKE: CSF1R and IL1R1 inhibitors synergistically attenuate the early pathogenesis of traumatic brain injury in mice. Neurotherapeutics 2025:e00787.

40. Heinz R, Brandenburg S, Nieminen-Kelhä M, Kremenetskaia I, Boehm-Sturm P, Vajkoczy P, Schneider UC: Microglia as target for anti-inflammatory approaches to prevent secondary brain injury after subarachnoid hemorrhage (SAH). J Neuroinflammation 2021, 18:36.

41. Elmore MR, Lee RJ, West BL, Green KN: Characterizing newly repopulated microglia in the adult mouse: impacts on animal behavior, cell morphology, and neuroinflammation. PLoS One 2015, 10:e0122912.

42. Lund H, Pieber M, Parsa R, Han J, Grommisch D, Ewing E, Kular L, Needhamsen M, Espinosa A, Nilsson E, et al: Competitive repopulation of an empty microglial niche yields functionally distinct subsets of microglia-like cells. Nat Commun 2018, 9:4845.

43. Shemer A, Grozovski J, Tay TL, Tao J, Volaski A, Süß P, Ardura-Fabregat A, Gross-Vered M, Kim JS, David E, et al: Engrafted parenchymal brain macrophages differ from microglia in transcriptome, chromatin landscape and response to challenge. Nat Commun 2018, 9:5206.

44. Colella P, Sayana R, Suarez-Nieto MV, Sarno J, Nyame K, Xiong J, Pimentel Vera LN, Arozqueta Basurto J, Corbo M, Limaye A, et al: CNS-wide repopulation by hematopoietic-derived microglia-like cells corrects progranulin deficiency in mice. Nat Commun 2024, 15:5654.

45. Yin F, Banerjee R, Thomas B, Zhou P, Qian L, Jia T, Ma X, Ma Y, Iadecola C, Beal MF, et al: Exaggerated inflammation, impaired host defense, and neuropathology in progranulin-deficient mice. J Exp Med 2010, 207:117–128.

46. Elmore MR, Najafi AR, Koike MA, Dagher NN, Spangenberg EE, Rice RA, Kitazawa M, Matusow B, Nguyen H, West BL, Green KN: Colony-stimulating factor 1 receptor signaling is necessary for microglia viability, unmasking a microglia progenitor cell in the adult brain. Neuron 2014, 82:380–397.

47. Albuquerque B, Haussler A, Vannoni E, Wolfer DP, Tegeder I: Learning and memory with neuropathic pain: impact of old age and progranulin deficiency. Front Behav Neurosci 2013, 7:174.

48. Fischer C, Endle H, Schumann L, Wilken-Schmitz A, Kaiser J, Gerber S, Vogelaar CF, Schmidt MHH, Nitsch R, Snodgrass I, et al: Prevention of age-associated neuronal hyperexcitability with improved learning and attention upon knockout or antagonism of LPAR2. Cell Mol Life Sci 2021, 78:1029–1050.

49. Vogel A, Ueberbach T, Wilken-Schmitz A, Hahnefeld L, Franck L, Weyer MP, Jungenitz T, Schmid T, Buchmann G, Freudenberg F, et al: Repetitive and compulsive behavior after Early-Life-Pain associated with reduced long-chain sphingolipid species. Cell Biosci 2023, 13:155.

50. Valek L, Tran B, Wilken-Schmitz A, Trautmann S, Heidler J, Schmid T, Brüne B, Thomas D, Deller T, Geisslinger G, et al: Prodromal sensory neuropathy in Pink1(-/-) SNCA(A53T) double mutant Parkinson mice. Neuropathol Appl Neurobiol 2021, 47:1060–1079.

51. Valek L, Tran BN, Tegeder I: Cold avoidance and heat pain hypersensitivity in neuronal nucleoredoxin knockout mice. Free Radic Biol Med 2022, 192:84–97.

52. Ge SX, Jung D, Yao R: ShinyGO: a graphical gene-set enrichment tool for animals and plants. Bioinformatics 2020, 36:2628–2629.

53. Eden E, Navon R, Steinfeld I, Lipson D, Yakhini Z: GOrilla: a tool for discovery and visualization of enriched GO terms in ranked gene lists. BMC Bioinformatics 2009, 10:48.

54. Sens A, Rischke S, Hahnefeld L, Dorochow E, Schäfer SMG, Thomas D, Köhm M, Geisslinger G, Behrens F, Gurke R: Pre-analytical sample handling standardization for reliable measurement of metabolites and lipids in LC-MS-based clinical research. J Mass Spectrom Adv Clin Lab 2023, 28:35–46.

55. Pang Z, Zhou G, Ewald J, Chang L, Hacariz O, Basu N, Xia J: Using MetaboAnalyst 5.0 for LC-HRMS spectra processing, multi-omics integration and covariate adjustment of global metabolomics data. Nat Protoc 2022, 17:1735–1761.

56. Smilde AK, Jansen JJ, Hoefsloot HC, Lamers RJ, van der Greef J, Timmerman ME: ANOVA-simultaneous component analysis (ASCA): a new tool for analyzing designed metabolomics data. Bioinformatics 2005, 21:3043–3048.

57. Lu J, Di Florio DN, Boya P, Maday S, Springer W, Chu CT: Autophagy and mitophagy at the synapse and beyond: implications for learning, memory and neurological disorders. Autophagy 2026, 22:10–52.

58. Huang M, Modeste E, Dammer E, Merino P, Taylor G, Duong DM, Deng Q, Holler CJ, Gearing M, Dickson D, et al: Network analysis of the progranulin-deficient mouse brain proteome reveals pathogenic mechanisms shared in human frontotemporal dementia caused by GRN mutations. Acta Neuropathol Commun 2020, 8:163.

59. Wallings RL, Gillett DA, Staley HA, Mahn S, Mark J, Neighbarger N, Kordasiewicz H, Hirst WD, Tansey MG: ASO-mediated knock-down of GPNMB in mutant-GRN and in Grn-deficient peripheral myeloid cells disrupts lysosomal function and immune responses. Mol Neurodegener 2025, 20:41.

60. Clelland CD, Fan L, Saloner R, Etchegaray JI, Altobelli CR, Salomonsson S, Maltos AM, Sachdev A, Zhu J, Lee SI, et al: Opposing role of phagocytic receptors MERTK and AXL in Progranulin deficient FTD. Commun Biol 2025, 8:971.

61. Zhang T, Feng T, Wu K, Guo J, Nana AL, Yang G, Seeley WW, Hu F: Progranulin deficiency results in sex-dependent alterations in microglia in response to demyelination. Acta Neuropathol 2023, 146:97–119.

62. Evers BM, Rodriguez-Navas C, Tesla RJ, Prange-Kiel J, Wasser CR, Yoo KS, McDonald J, Cenik B, Ravenscroft TA, Plattner F, et al: Lipidomic and Transcriptomic Basis of Lysosomal Dysfunction in Progranulin Deficiency. Cell Rep 2017, 20:2565–2574.

63. Marschallinger J, Iram T, Zardeneta M, Lee SE, Lehallier B, Haney MS, Pluvinage JV, Mathur V, Hahn O, Morgens DW, et al: Lipid-droplet-accumulating microglia represent a dysfunctional and proinflammatory state in the aging brain. Nat Neurosci 2020, 23:194–208.

64. Zhang J, Velmeshev D, Hashimoto K, Huang YH, Hofmann JW, Shi X, Chen J, Leidal AM, Dishart JG, Cahill MK, et al: Neurotoxic microglia promote TDP-43 proteinopathy in progranulin deficiency. Nature 2020, 588:459–465.

65. Hu C, Li T, Xu Y, Zhang X, Li F, Bai J, Chen J, Jiang W, Yang K, Ou Q, et al: CellMarker 2.0: an updated database of manually curated cell markers in human/mouse and web tools based on scRNA-seq data. Nucleic Acids Res 2023, 51:D870–d876.

66. Marian OC, Matis S, Dobson-Stone C, Kim WS, Kwok JB, Piguet O, Halliday GM, Landin-Romero R, Don AS: Reduced plasma hexosylceramides in frontotemporal dementia are a biomarker of white matter integrity. Alzheimers Dement (Amst) 2025, 17:e70131.

67. Fernàndez-Bernal A, Sol J, Galo-Licona JD, Mota-Martorell N, Mas-Bargues C, Belenguer-Varea Á, Obis È, Viña J, Borrás C, Jové M, Pamplona R: Phenotypic upregulation of hexocylceramides and ether-linked phosphocholines as markers of human extreme longevity. Aging Cell 2025, 24:e14429.

68. Chen D, Wang C, Chen X, Li J, Chen S, Li Y, Ma F, Li T, Zou M, Li X, et al: Brain-wide microglia replacement using a nonconditioning strategy ameliorates pathology in mouse models of neurological disorders. Sci Transl Med 2025, 17:eads6111.

69. Varvel NH, Grathwohl SA, Baumann F, Liebig C, Bosch A, Brawek B, Thal DR, Charo IF, Heppner FL, Aguzzi A, et al: Microglial repopulation model reveals a robust homeostatic process for replacing CNS myeloid cells. Proc Natl Acad Sci U S A 2012, 109:18150–18155.

70. Church KA, Rodriguez D, Vanegas D, Gutierrez IL, Cardona SM, Madrigal JLM, Kaur T, Cardona AE: Models of microglia depletion and replenishment elicit protective effects to alleviate vascular and neuronal damage in the diabetic murine retina. J Neuroinflammation 2022, 19:300.

71. Lefèvre-Arbogast S, Hejblum BP, Helmer C, Klose C, Manach C, Low DY, Urpi-Sarda M, Andres-Lacueva C, González-Domínguez R, Aigner L, et al: Early signature in the blood lipidome associated with subsequent cognitive decline in the elderly: A case-control analysis nested within the Three-City cohort study. EBioMedicine 2021, 64:103216.

72. Mi Y, Qi G, Vitali F, Shang Y, Raikes AC, Wang T, Jin Y, Brinton RD, Gu H, Yin F: Loss of fatty acid degradation by astrocytic mitochondria triggers neuroinflammation and neurodegeneration. Nat Metab 2023, 5:445–465.

73. Kwon YH, Kim J, Kim CS, Tu TH, Kim MS, Suk K, Kim DH, Lee BJ, Choi HS, Park T, et al: Hypothalamic lipid-laden astrocytes induce microglia migration and activation. FEBS Lett 2017, 591:1742–1751.

74. He Z, Ong CH, Halper J, Bateman A: Progranulin is a mediator of the wound response. Nat Med 2003, 9:225–229.

75. Li SS, Zhang MX, Wang Y, Wang W, Zhao CM, Sun XM, Dong GK, Li ZR, Yin WJ, Zhu B, Cai HX: Reduction of PGRN increased fibrosis during skin wound healing in mice. Histol Histopathol 2018:18076.

76. Rojo R, Raper A, Ozdemir DD, Lefevre L, Grabert K, Wollscheid-Lengeling E, Bradford B, Caruso M, Gazova I, Sánchez A, et al: Deletion of a Csf1r enhancer selectively impacts CSF1R expression and development of tissue macrophage populations. Nat Commun 2019, 10:3215.

77. Lonardi S, Scutera S, Licini S, Lorenzi L, Cesinaro AM, Gatta LB, Castagnoli C, Bollero D, Sparti R, Tomaselli M, et al: CSF1R Is Required for Differentiation and Migration of Langerhans Cells and Langerhans Cell Histiocytosis. Cancer Immunol Res 2020, 8:829–841.

78. Wang Y, Szretter KJ, Vermi W, Gilfillan S, Rossini C, Cella M, Barrow AD, Diamond MS, Colonna M: IL-34 is a tissue-restricted ligand of CSF1R required for the development of Langerhans cells and microglia. Nat Immunol 2012, 13:753–760.

79. Filiano AJ, Martens LH, Young AH, Warmus BA, Zhou P, Diaz-Ramirez G, Jiao J, Zhang Z, Huang EJ, Gao FB, et al: Dissociation of frontotemporal dementia-related deficits and neuroinflammation in progranulin haploinsufficient mice. J Neurosci 2013, 33:5352–5361.

80. Wang Y, Wernersbach I, Strehle J, Li S, Appel D, Klein M, Ritter K, Hummel R, Tegeder I, Schäfer MKE: Early posttraumatic CSF1R inhibition via PLX3397 leads to time– and sex-dependent effects on inflammation and neuronal maintenance after traumatic brain injury in mice. Brain Behav Immun 2022, 106:49–66.

81. Torres-Rodriguez O, Ortiz-Nazario E, Rivera-Escobales Y, Velazquez B, Colón M, Porter JT: Sex-dependent effects of microglial reduction on impaired fear extinction induced by single prolonged stress. Front Behav Neurosci 2022, 16:1014767.

82. Berve K, West BL, Martini R, Groh J: Sex– and region-biased depletion of microglia/macrophages attenuates CLN1 disease in mice. J Neuroinflammation 2020, 17:323.

83. Hardt S, Heidler J, Albuquerque B, Valek L, Altmann C, Wilken-Schmitz A, Schafer MKE, Wittig I, Tegeder I: Loss of synaptic zinc transport in progranulin deficient mice may contribute to progranulin-associated psychopathology and chronic pain. Biochim Biophys Acta 2017, 1863:2727–2745.

84. Kalf RS, Luinenburg MJ, Dematteis G, Scheper M, Anink JJ, Cavallo G, Mattarei A, Van Hecke W, Mühlebner A, Tapella L, et al: TSC-associated microglial hyperactivity: enhanced calcium signaling, metabolism, and phagocytosis. Acta Neuropathol 2026, 151:16.

85. Lane-Donovan C, Paredes M, Kao AW: The lysosome and proteostatic stress at the intersection of pediatric neurological disorders and adult neurodegenerative diseases. Prog Neurobiol 2025, 255:102854.

86. Zou X, Shi M, Xiao X, Lv X, Yang M, Tian M, Xie B, Wang L, Wang J, Qin D: Focusing on microglial mitochondria-lysosome crosstalk and neuroinflammation underlying depression: from molecular pathways to potential therapeutic interventions. Front Immunol 2026, 17:1775841.

87. Rodríguez-Periñán G, de la Encarnación A, Moreno F, López de Munain A, Martínez A, Martín-Requero Á, Alquézar C, Bartolomé F: Progranulin Deficiency Induces Mitochondrial Dysfunction in Frontotemporal Lobar Degeneration with TDP-43 Inclusions. Antioxidants (Basel) 2023, 12.

88. Le LHD, O’Banion MK, Majewska AK: Partial microglial depletion and repopulation exert subtle but differential effects on amyloid pathology at different disease stages. Sci Rep 2024, 14:30912.

89. Piccioni G, Maisto N, d’Ettorre A, Strimpakos G, Nisticò R, Triaca V, Mango D: Switch to phagocytic microglia by CSFR1 inhibition drives amyloid-beta clearance from glutamatergic terminals rescuing LTP in acute hippocampal slices. Transl Psychiatry 2024, 14:338.

90. Yang Y, Aloi MS, Cudaback E, Josephsen SR, Rice SJ, Jorstad NL, Keene CD, Montine TJ: Wild-type bone marrow transplant partially reverses neuroinflammation in progranulin-deficient mice. Lab Invest 2014, 94:1224–1236.

91. Davtyan H, Naguib S, Voskobiynyk Y, Chadarevian JP, Capocchi JK, Giacchino JL, Eskandari-Sedighi G, DeNittis V, Ford JB, Agababian A, et al: Transplantation of Human IPSC-derived Microglia Ameliorates Neuropathology and Circuit Dysfunction in Progranulin-Deficient Mice. Res Sq 2026.

92. Gelfand JM, Greenfield AL, Barkovich M, Mendelsohn BA, Van Haren K, Hess CP, Mannis GN: Allogeneic HSCT for adult-onset leukoencephalopathy with spheroids and pigmented glia. Brain 2020, 143:503–511.

93. Dulski J, Heckman MG, White LJ, Żur-Wyrozumska K, Lund TC, Wszolek ZK: Hematopoietic Stem Cell Transplantation in CSF1R-Related Leukoencephalopathy: Retrospective Study on Predictors of Outcomes. Pharmaceutics 2022, 14.

94. Percie du Sert N, Hurst V, Ahluwalia A, Alam S, Avey MT, Baker M, Browne WJ, Clark A, Cuthill IC, Dirnagl U, et al: The ARRIVE guidelines 2.0: updated guidelines for reporting animal research. BMJ Open Sci 2020, 4:e100115.

