## Supplemental Figures with legends for "Temporary deterioration of health and behavior during pexidartinib-mediated microglia depletion and repopulation in progranulin-deficient mice"

Supplementary figures and legends

Suppl. Figure S1A

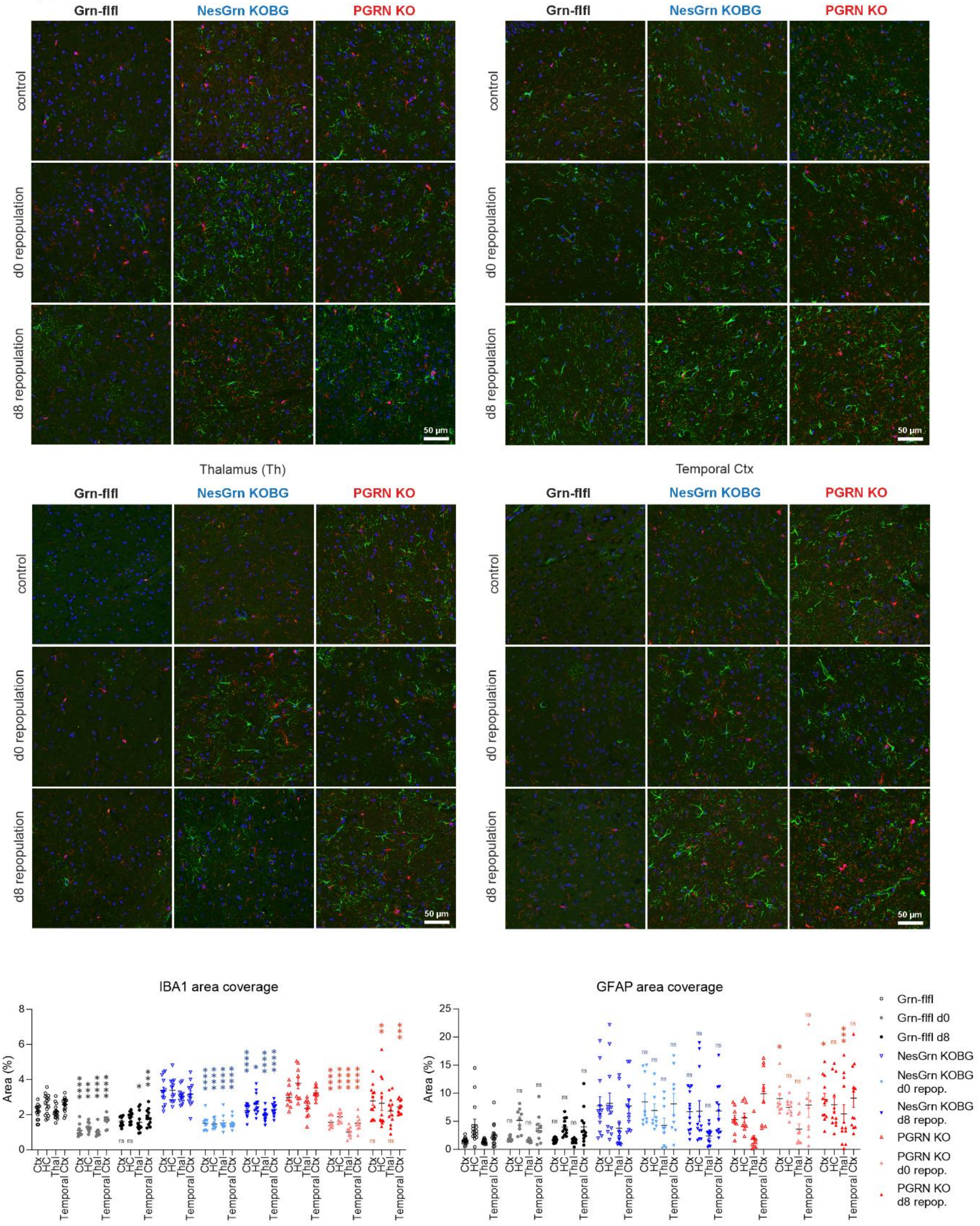

Suppl. Figure S1B

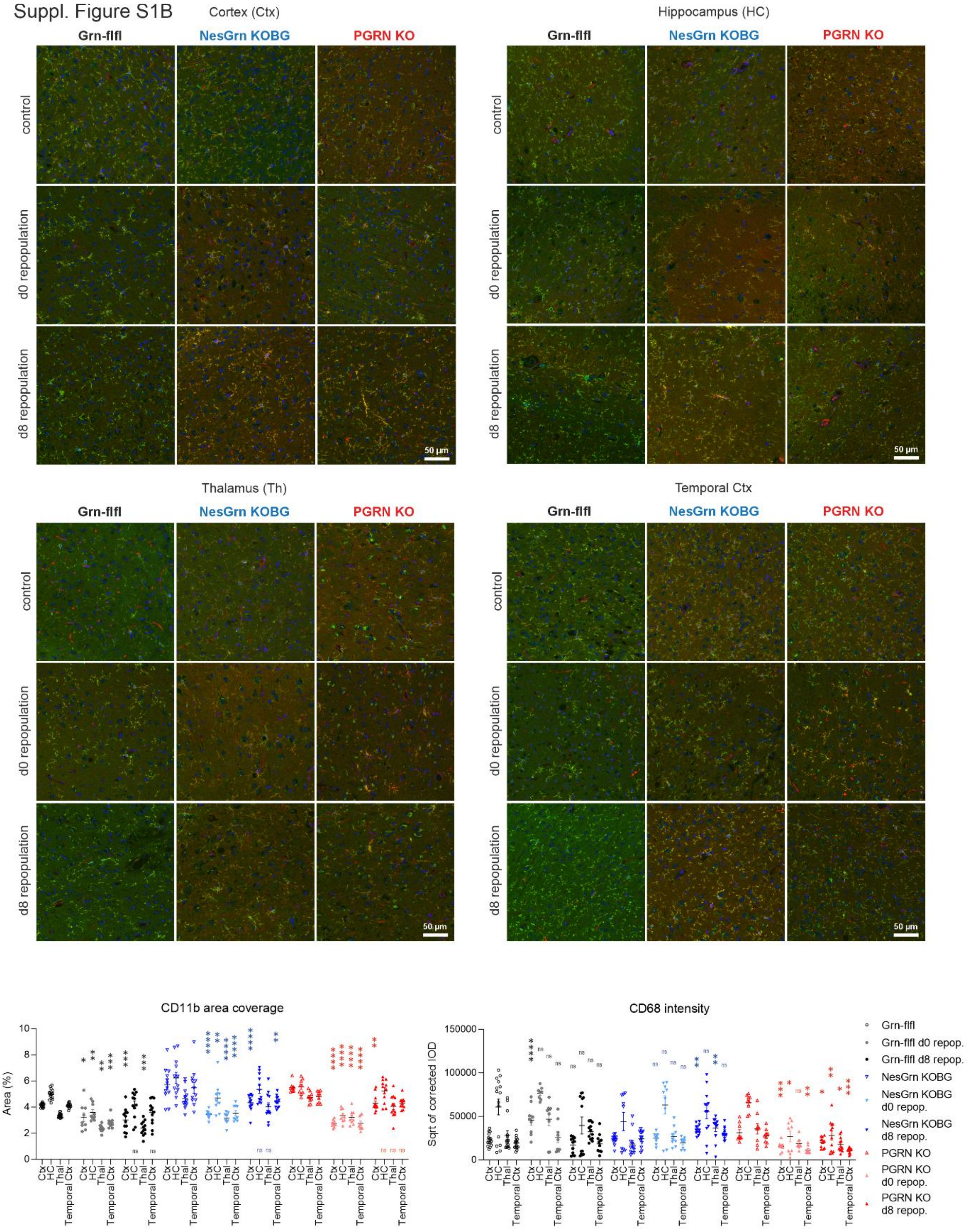

Suppl. Figure S1C

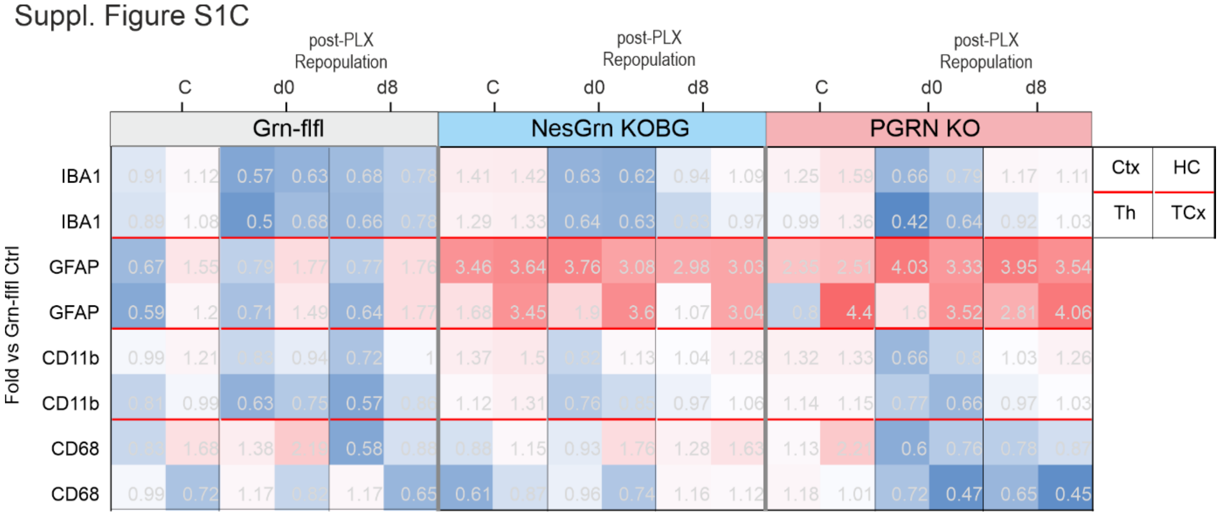


### Suppl. Figure S1

**Immunofluorescence of brain microglia depletion and repopulation**

**A:** Exemplary immunofluorescent images of IBA1 immunoreactive microglia and of GFAP immunoreactive astrocytes in the cortex, hippocampus, thalamus and temporal cortex of Grn-flfl control mice, of NesGrn KOBG mice and of full PGRN KO mice. DAPI is used as nuclear counterstain. For quantitative analysis, images were converted to binary images using auto-threshold in FIJI ImageJ, and the relative area covered by specific immunofluorescence for either IBA1 or GFAP was used for statistical comparison. The areas after PLX3397 were compared with the area coverage at baseline (no PLX). Each scatter shows the quantitative result of one non-adjacent image of 3-5 mice per genotype and treatment.

**B:** In analogy to A, the images show immunofluorescent images of the microglia activity markers CD11b and CD68, again DAPI as nuclear counterstain. For CD68 The squared root transformed corrected integrated optical density was used for quantitative analysis.

Data were submitted to 2-way ANOVA for “brain region” by “PLX3397 treatment”. Each region was then compared to the respective region of baseline group using an adjustment of alpha according to Dunnett. Asterisks reveal significant differences versus no PLX3397 treatment (baseline) P*<0.05, **<0.01, ***<0.001, ****<0.0001.

**C:** The heatmap summarizes the quantitative results where the immunofluorescent signal or area was normalized to the respective signal/area of the Grn-flfl control mice as fold-change and is depicted as blue (reduced) to red (increased) colour. Each square is one region.

Suppl. Figure S2

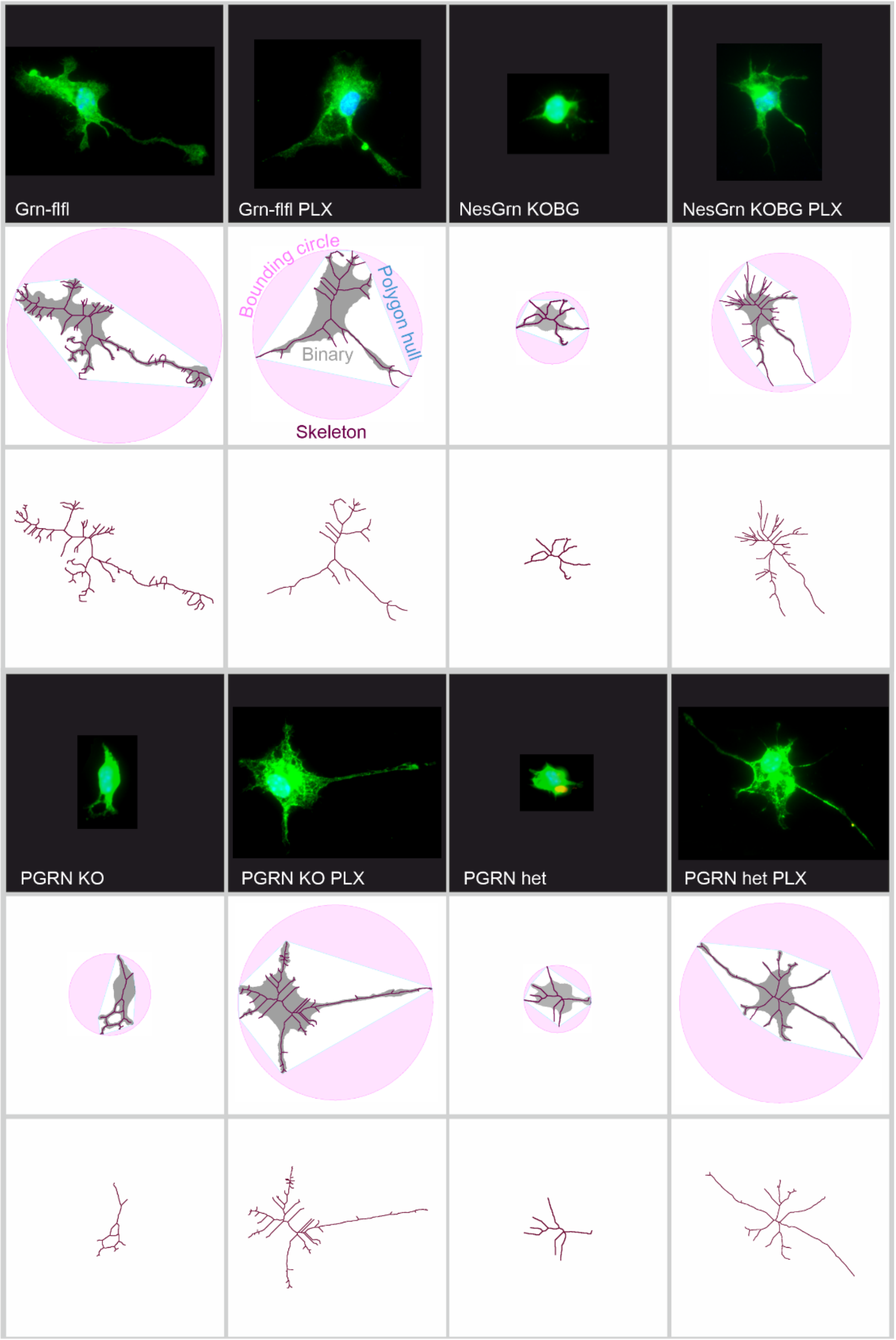


### Suppl. Figure S2

**Morphometric microglia analysis**

Microglia morphometry was obtained from primary microglia cultures from old mouse brains, stained with anti-IBA1 and DAPI. The analysis was based on the binary images, skeleton, polygon hull and bounding circle, from which the area, area coverage, fractal dimensions, number and length of branches and arborization were extracted. Quantitative features were normalized and submitted to multivariate analyses (main body).

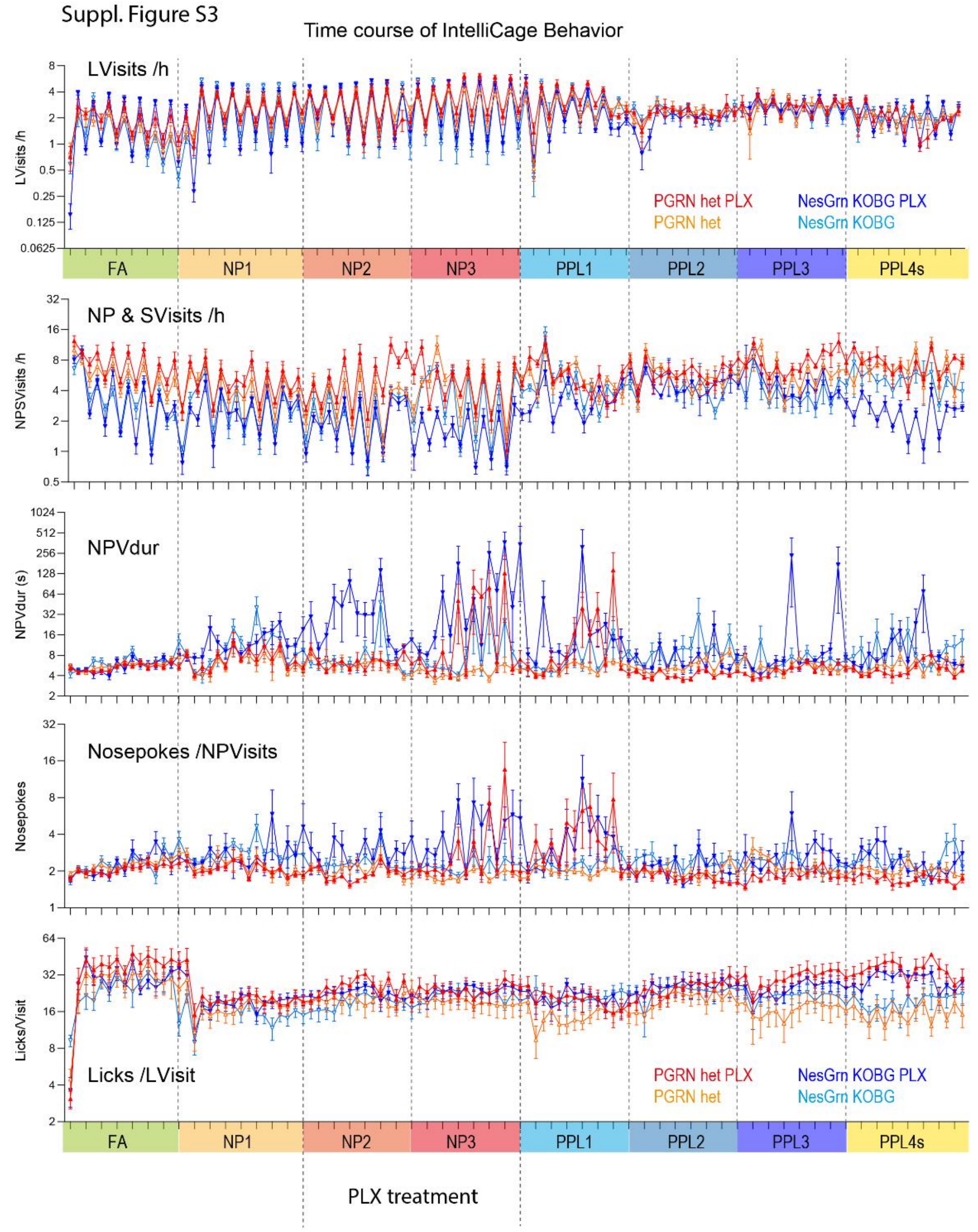


### Suppl. Figure S3

**Time course of IntelliCage nighttime and daytime behaviour in 12h Bins**

The panels show from top to bottoms:

Corner visits with licks per hour (LVisits /h), the sum of visits with nosepokes but without licks (NPVisits) plus visits without nosepokes or licks (SVisits) per hour (NPSVisits/h), the median duration of visits with nosepokes (NPVdur), the number of nosepokes per visit with nosepokes (NPVisits), and the number of licks per visit (Licks/Visit). The tasks are described at the bottom of the graph and the periods shaded in different colours. Further details about the tasks and IntelliCage abbreviations are shown in Suppl. Tables 2, 3. The abbreviations of the tasks are FA, Free adaptation; NP, nosepoke adaptation; PPL, place preference learning; PPLs ("s" for social challenge) where all mice of a cage are assigned to one rewarding corner. The data show means ± SEM of 7-8 female mice per group. The fluctuations show nighttime and daytime differences (12h Bins) and reveal the circadian rhythm. During and after the PLX3397 treatment period, there is a disruption of the nosepoke behaviour in both PLX groups.

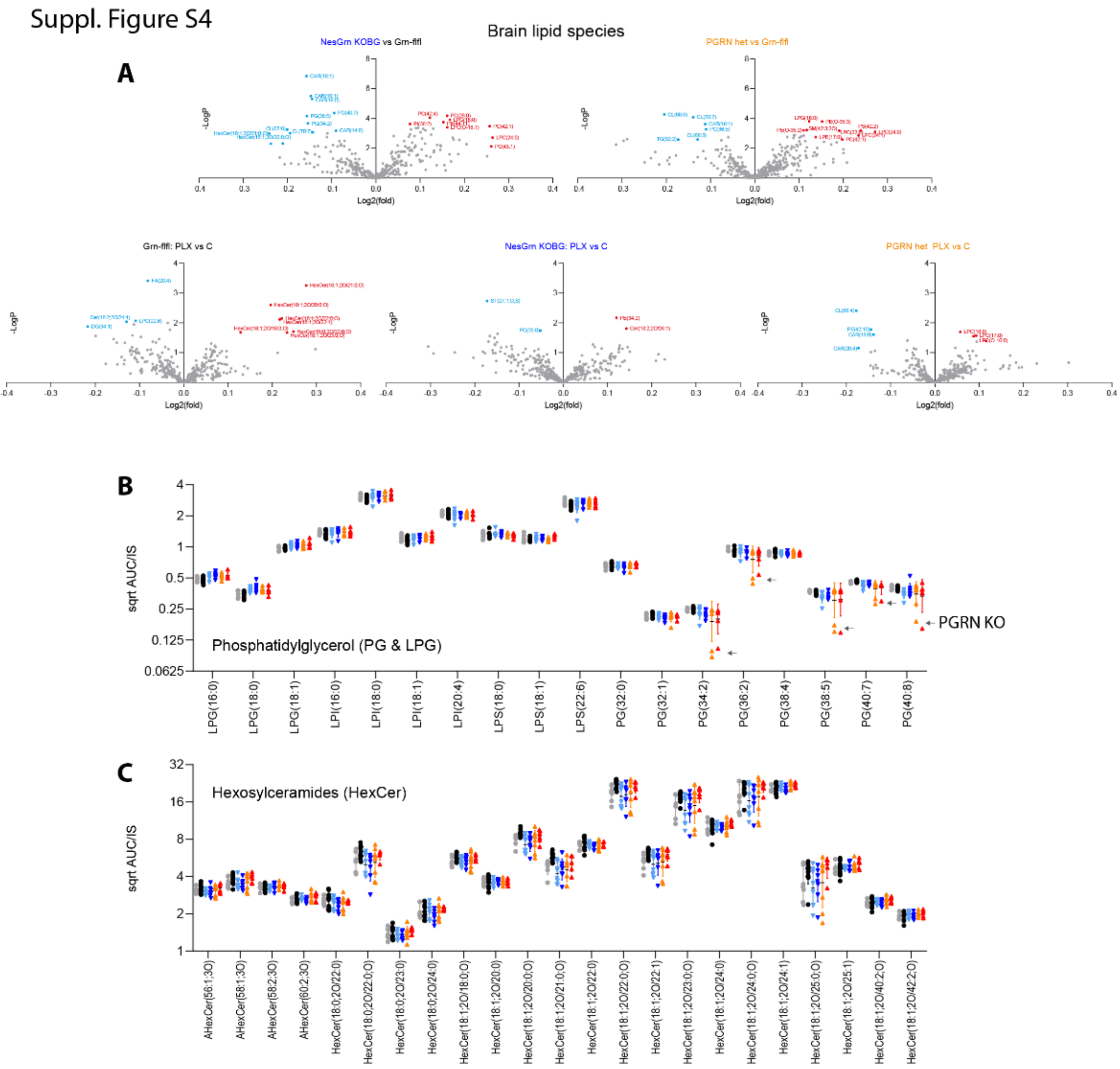


### Suppl. Figure S4

**Untargeted lipidomic analysis in the brain 8 weeks after PLX3397 treatment**

**A**: Volcano plots showing lipid species in the brain of NesGrn KOBG versus Grn-flfl control mice and of PGRN het mice versus Grn-flfl control mice in the upper row and PLX3397-treated versus control diet for each genotype in the bottom row. Lipids were analysed by untargeted UHPLC-MS/MS lipidomic screening. The X-axis shows the log2(Fold change), the Y-axis minus Log10 of the t-test P-value. Upregulated lipids are shown in red, downregulated in blue.

**B/C**: Scatter plots of square root transformed AUC/IS values of phosphatidylglycerol (PG) and related lipid species (LPG, LPI, LPS) and of hexosylceramides (HexCer). PG species and HexCer species were particularly low in few PGRN KO mice which were included and are depicted together with the PGRN het group.

Abbreviations: AHexCer, acetylhexosylceramides; CAR, acylcarnitines; Cer, ceramides; CE, cholesterol ester; CL, cardiolipins; DG, diglycerides; FA, fatty acids; HexCer, hexosylceramides; LPC, lysophosphatidylcholines; LPE, lysophosphatidylethanolamines; LPG, lysophosphatidylglycerols; LPI, lysophosphatidylinositols; PC, phosphatidylcholines; PE, phosphatidylethanolamines; PG, phosphatidylglycerols; PI, phosphatidylinositols; SM, sphingomyelins; ST, sterols; TG, triglycerides; –O ether bound lipids.

Suppl. Figure S5

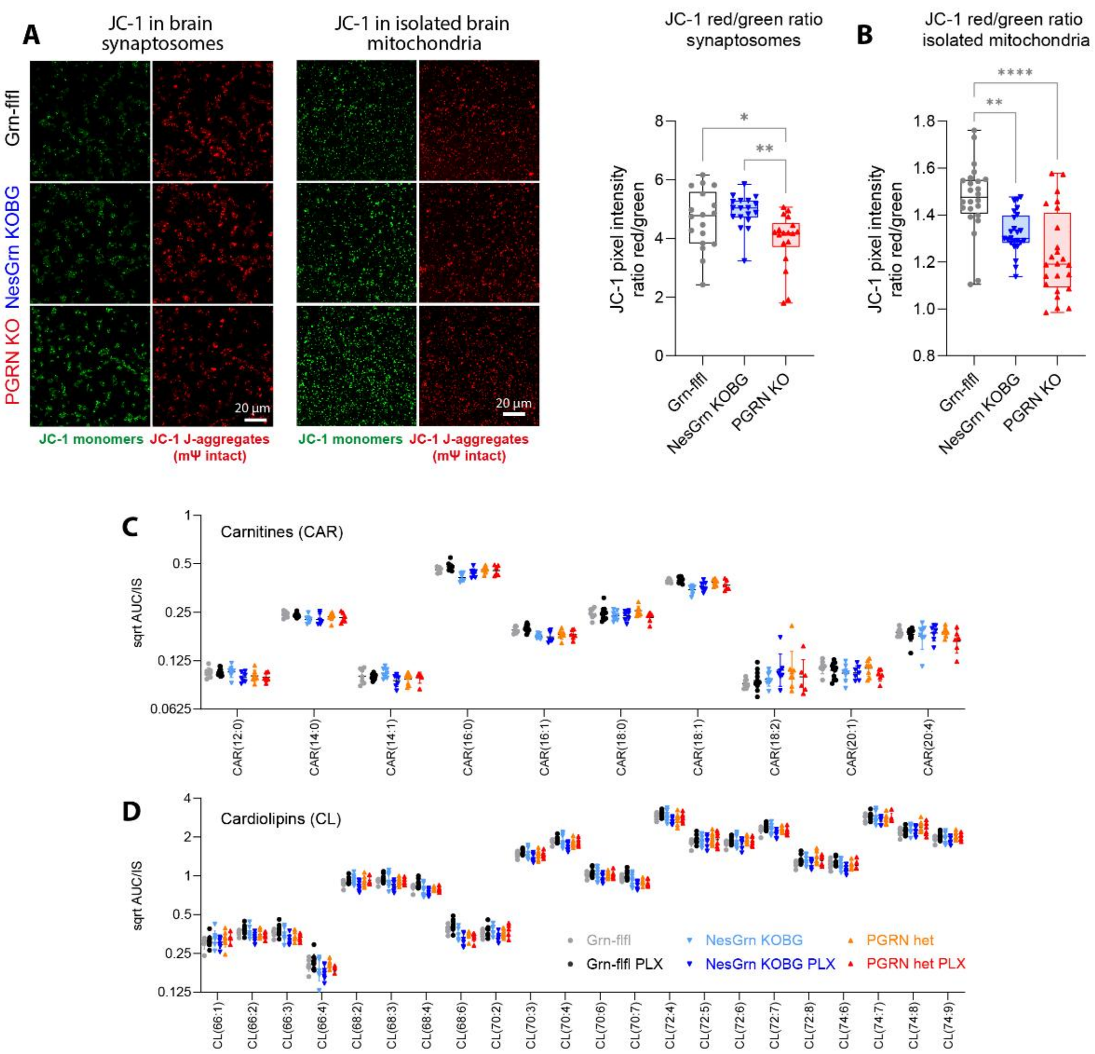


### Suppl. Figure S5

**JC-1 based mitochondrial membrane potential and mitochondrial lipids**

**A**: JC-1 immunofluorescent analysis of mitochondrial membrane potentials in brain synaptosomes and in isolated brain mitochondria. JC-1 dye accumulates in mitochondria in a potential dependent manner, forming red fluorescent J aggregates in polarized (high potential) mitochondria, whereas depolarized mitochondria retain JC 1 in its monomeric, green fluorescent form. The red/green fluorescence ratio was used as an indicator of mitochondrial membrane potential.

**B**: Box/scatter plots show the quantification of JC-1 red/green ratios per image obtained from n = 3 mice per group. The box is the interquartile range, whiskers show minimum to maximum. Data were submitted to univariate ANOVA and subsequent Dunnett posthoc comparison versus Grn-flfl controls. Asterisks show significant differences, *P<0.05, **P<0.01; ****P<0.0001.

**C, D**: Lipidomic analyses in brain tissue 8 weeks after PLX3397 treatment revealed low acylcarnitines and cardiolipins in progranulin deficient mice, both in NesGrn KOBG and PGRN het mice. Scatter plots show square root transformed AUC/IS values of long chain acyl-carnitines (CAR) and of cardiolipins (CL). Acylcarnitines are required for transfer of fatty acids into mitochondria to fuel beta-oxidation. Cardiolipins are mitochondria specific lipids of the inner mitochondrial membrane. The CL deficit in NesGrn KOBG brains was stronger after PLX treatment than without.

Suppl. Figure S6

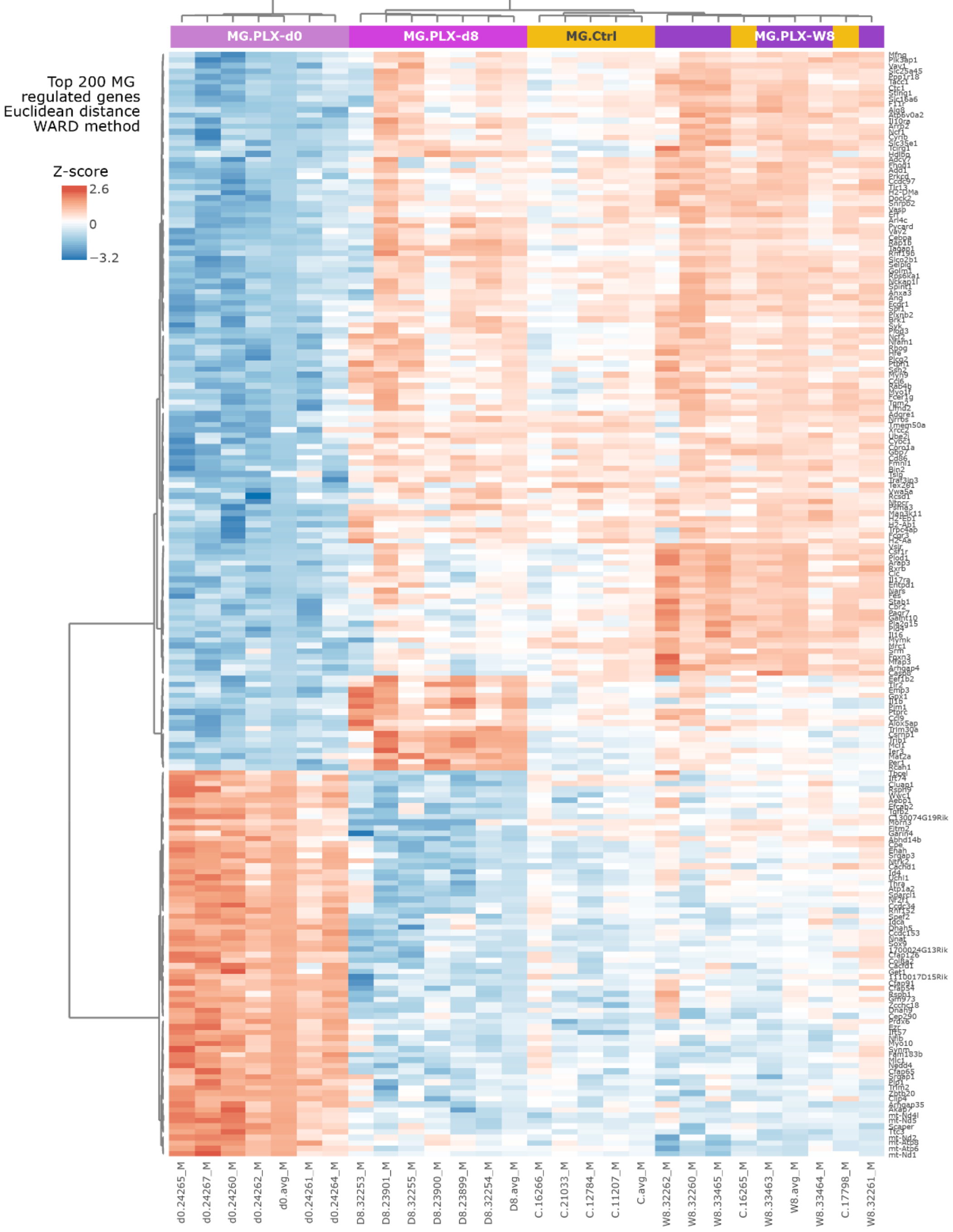


### Suppl. Figure S6

**Heatmap with dendrograms of top 200 regulated genes in microglia before/after PLX3397**

NesGrn KOBG mice were treated with PLX3397 diet for 14 days. Brain microglia were isolated at the end of the PLX3397 diet (d0-repopulation) and 8d and 8 weeks after the end (8d-repopulation, W8-repopulation), and from control mice who received normal diet. Each group comprised 6 mice. Total RNA of microglia and non-microglial cells was subjected to 3' mRNA sequencing. Top 200 genes were selected on the basis of ANOVA statistics that compared the transcriptome after PLX with control conditions. Gene and mice are clustered according to Euclidean distance metrics using the Ward method. For depiction as heatmap normalized counts were auto-scaled to have a common average and variance of 1.

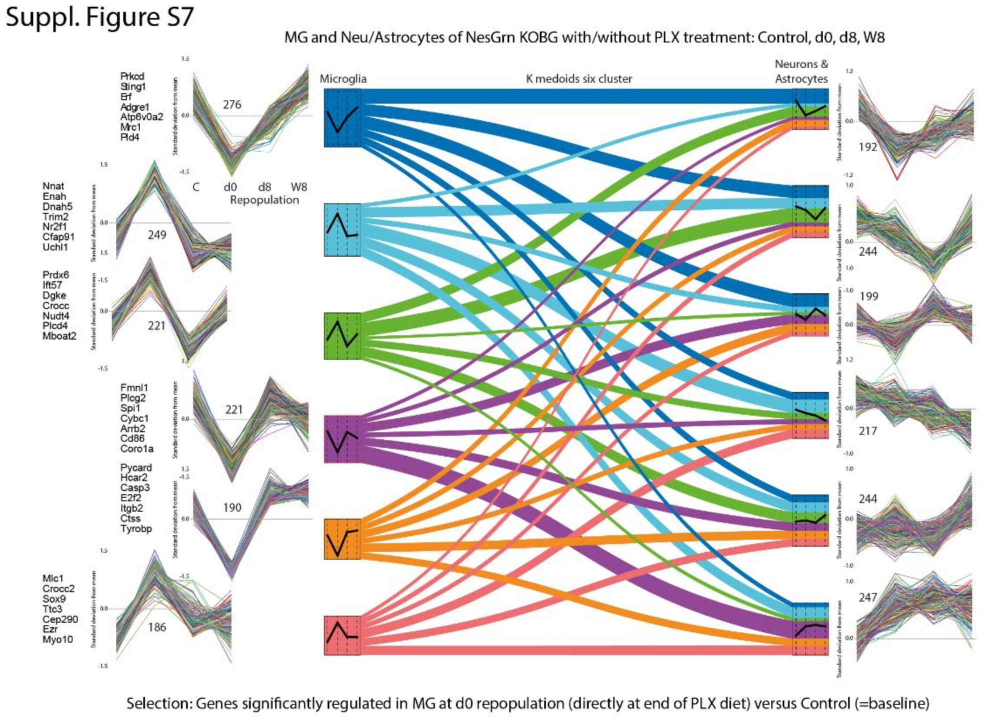


### Suppl. Figure S7

**K-medoids and alluvial plot of MG candidates and representation in non-microglial cells**

Transcriptomic data were obtained from NesGrn KOBG microglia and non-microglial cells as described in S6. Non-microglial cells were a mixture of astrocytes and neurons based on marker gene expression. For K-means clustering, genes were selected which differed significantly in microglia at the end of the PLX diet, i.e. at time "0" of the repopulation versus control conditions (1343 genes). The cluster show 2 major patterns (i) drop and return or (ii) raise and return but with differences in the early and late repopulation with either slow return or overshooting return. Most of the genes are also regulated in neurons/astrocytes but regulations are softer and the genes contribute to different clusters with in part opposing time-dependent changes.

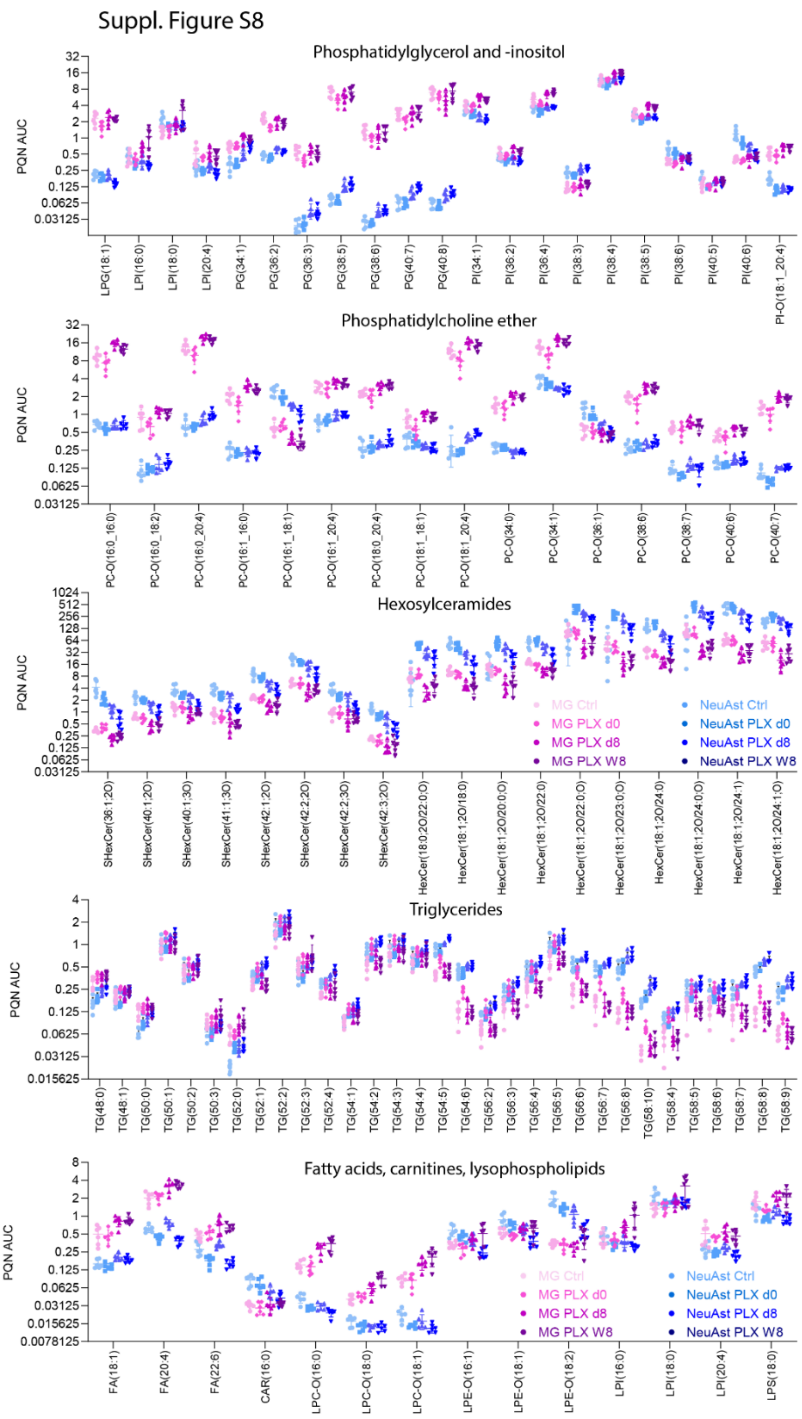


### Suppl. Figure S8

**Lipidomic species in primary brain microglia and non-microglia after PLX3397 treatment**

Microglia and non-microglia were obtained from NesGrn KOBG mouse brains as described in S6. Lipids were extracted from 250,000 cells and subjected to UHPLC/MS-MS lipidomic screening.

The scatter plots show the PQN normalized AUC values of lipid species of lipid classes which were regulated during PLX3397 induced microglia depletion and repopulation. The values of microglia were further sum-normalized to the total lipids of neurons/astrocytes to allow for a comparison of lipid distributions between cell types, which differ in cell size and therefore lipid content. The scatters show the data of each one mouse. The statistics are shown for lipid classes in the main body (Figure 13).
