## Supplemental Tables for "Temporary deterioration of health and behavior during pexidartinib-mediated microglia depletion and repopulation in progranulin-deficient mice"

### Supplementary Tables

#### Suppl. Table 1

Overview of mouse groups, ages and sex in behavioural and biological experiments to assess effects of PLX3397 mediated microglia depletion and repopulation in progranulin deficient (PGRN KO, PGRN het, NesGrn KOBG) and control mice (Grn-flfl)

| Experiment | Readout | Interpretation | Grn-flfl | NesGrn KOBG | PGRN KO/het |
|---|---|---|---|---|---|
| **IntelliCage** | Learning and memory | Activity, exploratory behaviour, Learning/memory, social interaction | Female, n = 3, age 14-15 months at start | Female, n = 16, age 8-16 months at start | Female, n = 15, age 9-14 months at start |
| **TGR** | Activity, locomotion, temperature preference | Activity, exploratory behaviour, sensory functions | | Female, n = 13, age 11-19 months at start | Female, n = 13, age 12-17 months at start |
| **snRNAseq** | Single nucleus mRNA | Differential gene expression | Male, n = 2, age 23 months | Male, n = 2, age 19 or 23 months | M/F, n = 3, age 17 months |
| **RNAseq** | mRNA brain | Differential gene expression | Female, n = 16, End of IC, age 9-18 months | Female, n = 16, End of IC, age 11-19 months | Female, n = 15, End of IC, age 12-17 months |
| **Lipidomic** | Brain lipids | Structural and metabolic brain alterations, lipid-laden microglia | Female, n = 16, End of IC, age 9-18 months | Female, n = 16, End of IC, age 11-19 months | Female, n = 15, End of IC, age 12-17 months |
| **Microglia morphology** | Morphology | Differential cell morphology | Male, n = 6, age 15-20 months<br>Female, n = 3, age 7-9 months | Male, n = 4, age 18 months<br>Female, n = 3, age 9-10 months | Male, n = 4 (PGRN het), age 16 months, n = 4 (PGRN KO), age 16 months<br>Female, n = 8 (PGRN het), age 7-18 months, n = 4 (PGRN KO), age 10-16 months |
| **Microglia BODIPY** | Lipid droplets | Lipid laden microglia | Male, n = 6, age 8-26 months | Female, n = 2, age 12-13 months,<br>Male, n = 4, age 11-12 months | Male, n = 6, age 9-24 months |
| **Microglia Seahorse** | Mitochondrial function | Respiratory functions of primary microglia | Male, n = 9, age 8-18 months,<br>Female, n = 3, age 13-22 months | Male, n = 8, age 11-20 months<br>Female, n = 4, age 11-20 months | Male, n = 9, age 10-20 months<br>Female, n = 3, age 10-16 months |
| **Microglia – NeuAst Lipidomic and RNAseq** | Cell type specific lipidomic profiles | Cell specific lipidomic and transcriptomic profiles w/wo PLX | | Male, n = 15, age 11-24 months,<br>Female, n = 9, age 13-18 months | |

| | | | | | |
|---|---|---|---|---|---|
| **Gliosis histology** | Gliosis | Differential cell coverage in different brain regions | Female, n = 12, age 5-18 months | Female, n = 11, age 15-22 months | Female, n = 10, age 10-15 months |
| **Synapse histology** | Synaptic density brain | Differential PSD95 and synaptophysin coverage in cortex and HC | Female, n = 12, age 5-18 months | Female, n = 11, age 15-22 months | Female, n = 10, age 10-15 months |
| **Spine density** | Spine density | Differential spine density | Female, n = 12, age 5-18 months | Female, n = 11, age 15-22 months | Female, n = 10, age 10-15 months |
| **rtPCR candidate genes** | Gene expression as log2(Fold mRNA) | Differential gene expression of isolated microglia cells | Male, n = 13, age 6-19 months Female, n = 16, age 7-19 months | Male, n = 12, age 6-18 months Female, n = 9, age 10-18 | Male, n = 5 (PGRN het), age 10-18 months, n = 15 (PGRN KO), age 10-17 months Female, n = 7 (PGRN het), age 6-13 months, n = 9 (PGRN KO), age 10-17 |
| **Mitochondrial membrane potential** | Mitochondrial function | Membrane potential | M/F, n = 3, age 11-12 months | M/F, n = 3, age 12-17 months | Female, n = 3, age 15 months |

#### Suppl. Table 2

IntelliCage Tasks

| Experiment | Duration | Task description | Readouts, interpretation |
|---|---|---|---|
| **Free adaptation (FA)** | 8 days | General habituation to the cage with open access to every corner, all doors open, and water/food ad libitum | Exploratory behaviour, activity, circadian rhythms, social interaction |
| **Nosepoke adaptation (NP)** | 8 days | The first nosepoke of a visit opened the door for 5 s. To drink more, the mouse has to start a new visit. | Exploratory behaviour, activity, circadian rhythms, social interaction |
| **Nosepoke adaptation (NP2)** | 7 days | Same protocol as in “nosepoke adaptation”. Treatment of cage-1 animals with pexidartinib chow. | Exploratory behaviour, activity, circadian rhythms, social interaction |
| **Nosepoke adaptation (NP3)** | 7 days | Same protocol as in “nosepoke adaptation”. Treatment of cage-1 animals with pexidartinib chow. | Exploratory behaviour, activity, circadian rhythms, social interaction |
| **Place preference learning (PPL1)** | 7 days | Mice were allowed to drink in one of the 4 corners. The first correct nosepoke of a visit opened the door for 5 s. To drink more, the mouse has to start a new visit. 4 mice were assigned to one corner. Learning was supported by LED | Spatial preference learning by reward, exploratory behaviour, activity, social interaction |
| **Place preference reversal 2 (PPL2)** | 7 days | Equal protocol as in “place preference learning” but with the opposite corner as the correct corner. Learning was supported by LED | Cognitive flexibility of Reversal Learning requires the dorsal and ventral hippocampus and their functional interactions with the prefrontal cortex (Vila-Ballo, Mas-Herrero et al. 2017, Avigan, Cammack et al. 2020) |

| | | | |
|---|---|---|---|
| **Place preference reversal 3 (PPL3)** | 7 days | Equal protocol as in “place preference learning” but with a new corner as the correct corner. Learning was supported by LED | Cognitive flexibility of Reversal Learning requires the dorsal and ventral hippocampus and their functional interactions with the prefrontal cortex (Vila-Ballo, Mas-Herrero et al. 2017, Avigan, Cammack et al. 2020) |
| **Place preference learning social (PPL4s)** | 3 days | All mice are assigned the same corner as correct between 12-10 am. Between 10-12 am all corners are assigned correct. | Social structure |

#### Suppl. Table 3

Abbreviations of behavioural parameters of IntelliCage experiments

| Parameter | Description |
|---|---|
| **Visits** | Visits / h |
| **NPvisits** | Visits with Nosepoke without Licks / h |
| **Lvisits** | Visits with Licks / h |
| **SVisits** | Visits without Licks and without Nosepokes / h |
| **NPVdur** | Median duration of Visits with NP w/out Lick (s) |
| **Nosepokes (NP)** | Mean number of Nosepokes during Visits with NP w/out Licks |
| **NPduration** | Median duration of such Nosepokes during a Visit (s) |
| **Licks** | Median number of Licks per Visit |
| **Lduration** | Median duration of Licking during a Visit (s) |
| **Lcontact** | Median bottle cap contact time during a Visit (s) |
| **Nocturnal** | Log(Visit frequency during dark phase / Visit frequency during light phase) |
| **Repetitive** | Repetitiveness, log(sum of observed returns to same corner / sum of expected such switches) |
| **IVI** | Intervisit intervals (s) i.e. time from end of visit to start of next corner visit |
| **IVIrepdens** | Intervisit intervals (s) for repeated use of the same corner |
| **Unevenness** | Describes the relative use of corners, ranges from 0-1 (0=equal use of 4 corners, 1=exclusive use of 1 corner) |
| **Sidedness** | Ratio of visits with first left versus first right NP of visits with NPs |
| **Mesor** | **M**idline **es**timating **s**tatistic **o**f **r**hythm. The mesor is a circadian rhythm-adjusted mean based on the parameters of a cosine function fitted to the raw data of the visits. |
| **Amplitude** | Difference between Mesor and Peak activity |
| **Acrophase** | Time to maximum activity after Light Off (Light off set to 0) |
| **Period** | Duration of one circadian cycle |

#### Suppl. Table 4

Used primer in rtPCR candidate genes experiments

| **Gene name, (amplicon size, annealing temperature)** | **Oligonucleotide sequences 5’–3’** <br> **(fw: forward, rev: reverse)** | **Gene bank number** |
|---|---|---|
| ***Ppia* (144 bp, 60°C)** | fw- GCTGGACCAAACACAAAACGG <br> rev- GCCATTCCTGGACCCAAAAC | NM_008907 |
| ***Grn* (171 bp, 60°C)** | fw- CTGCCCGTTCTCTAAGGGTG <br> rev- ATCCCCACGAACCATCAACC | NM_008175 |
| ***Gapdh* (100 bp, 60°C)** | fw- CCTCGTCCCGTAGACAAAATG <br> rev- TCTCCACTTTGCCACTGCAA | NM_001289726 |
| ***Nos2* (127 bp, 60°C)** | fw- GTTCTCAGCCCAACAATACAAGA | NM_010927 |

| | | |
|---|---|---|
| | rev- GTGGACGGGTCGATGTCAC | |
| ***Tgfb1* (133 bp, 60°C)** | fw- CTCCCGTGGCTTCTAGTGC<br>rev- GCCTTAGTTTGGACAGGATCTG | NM_011577 |
| ***P2ry12* (77 bp, 60°C)** | fw- QuantiTect Panel Qiagen<br>rev- QuantiTect Panel Qiagen | NM_027571 |
| ***Ube2d2* (125 bp, 60°C)** | fw- TGTCCATCTGTTCTCTGTTGTGTG<br>rev- ATACTTCTGAGTCCATTCCCGC | NM_019912 |
| ***Csf1r* (96 bp, 60°C)** | fw- QuantiTect Panel Qiagen<br>rev- QuantiTect Panel Qiagen | NM_001037859 |
| ***Cx3cr1* (63 bp, 60°C)** | fw- QuantiTect Panel Qiagen<br>rev- QuantiTect Panel Qiagen | NM_009987 |
| ***P2rx4* (73 bp, 60°C)** | fw- QuantiTect Panel Qiagen<br>rev- QuantiTect Panel Qiagen | NM_011026 |
| ***Trem2* (132 bp, 60°C)** | fw- QuantiTect Panel Qiagen<br>rev- QuantiTect Panel Qiagen | NM_031254 |
| ***Apoe* (135 bp, 60°C)** | fw- QuantiTect Panel Qiagen<br>rev- QuantiTect Panel Qiagen | NM_009696 |
| ***Hexb* (137 bp, 60°C)** | fw- QuantiTect Panel Qiagen<br>rev- QuantiTect Panel Qiagen | NM_010422 |
| ***Il10* (105 bp, 60°C)** | Fw- GCTCTTACTGACTGGCATGAG<br>Rev- CGCAGCTCTAGGAGCATGTG | NM_010548 |
| ***Il1b* (89 bp, 60°C)** | Fw- GCAACTGTTCCTGAACTCAACT<br>Rev- ATCTTTTGGGGTCCGTCAACT | NM_008361 |
| ***Il6* (166 bp, 60°C)** | Fw- CCGGAGAGGAGACTTCACAG<br>Rev- TTCTGCAAGTGCATCATCGT | NM_031168 |
| ***Lamp2* (111 bp, 60°C)** | Fw- ATGTGCCTCTCTCCGGTTAAA<br>Rev- GCAAGTACCCTTTGAATGTCA | NM_01017959 |
| ***Cd68* (182 bp, 60°C)** | Fw- AGGGTGGAAGAAAGGCTTGG<br>Rev- ACTCGGGCTCTGATGTAGGT | NM_001291058 |
| ***Ptgs2* (87 bp, 60°C)** | Fw- AGACACTCAGGTAGACATGATCTACCCT<br>Rev- GGCACCAGACCAAAGACTTCC | NM_011198 |
| ***Hdac1* (100 bp, 60°C)** | Fw- CGAATCCGCATGACTCACAAT<br>Rev- ACTTGGTCATCTCCTCAGCAT | NM_008228 |
| ***Hdac2* (111 bp, 60°C)** | Fw- ACCGGCAACAAACTGATATGG<br>Rev- CGAGGATGGCAAGCACAATAT | NM_008229 |
| ***Hdac3* (138 bp, 60°C)** | Fw- ACCCAGTGTCCAGATTCATGA<br>Rev- TGGTCGCCATCATAGAACTCA | NM_010411 |
| ***Tfeb* (119 bp, 60°C)** | Fw- TATCAGCTCCAACCCCGAGA<br>Rev- TAGACAGGGGCAGCGTGTTA | NM_001161722 |
| ***Trpml1* (175 bp, 60°C)** | Fw- CGGTGTCATTCGCTACCTGA<br>Rev- CAGCGAGCGGAACTTCACAT | NM_053177 |
| ***Lyz2* (163 bp, 60°C)** | Fw- AAGGCATTCGAGCATGGGTG<br>Rev- TCGAGGGAATGTGACCTCTCT | NM_017372.3 |
| ***Psap* (214 bp, 60°C)** | Fw- ATTGCAGCCTGCGAAGTGAA<br>Rev- AGGAAGGGATTTCGCTGTGG | NM_001146120 |
| ***Laptm5* (80 bp, 60°C)** | Fw- CGTGCTCATCCTGAAGGTCTA<br>Rev- CCATGGCCGAATTCATGTGC | NM_010686.4 |
| ***Ctsd* (186 bp, 60°C)** | Fw- AATCCCTCTGCGCAAGTTCA<br>Rev- CGCCATAGTACTGGGCATCC | NM_009983 |
| ***Lamp1* (103 bp, 60°C)** | Fw- CAGCACTCTTTGAGGTGAAAAAC<br>Rev- TGCGAATGGTTCTCAGATCGT | NM_010684 |
| ***Gpnmb* (134 bp, 60°C)** | Fw- TCTATCCCTGGCAAAGACCCAG<br>Rev- ATTGGCTTGTACGCCTTGTG | NM_053110 |
| ***Mt-Nd1* (147 bp, 60°C)** | Fw- GCTTTACGAGCCGTAGCCCA<br>Rev- GGGTCAGGCTGGCAGAAGTAA | |
| ***Cnr1* (126 bp, 60°C)** | Fw- GTGCTGTTGCTGTTCATTGTG<br>Rev- CTTGCCATCTTCTGAGGTGTG | NM_007726 |
| ***Scd1* (122 bp, 60°C)** | Fw- TGGAGACGGGAGTCACAAGA<br>Rev- ACACCCCGATAGCAATATCCAG | NM_009127.4 |
| ***Fnbp1* (190 bp, 60°C)** | Fw- CACCGAAGTCACCCAGAACA<br>Rev- ACTACCATCTGGGCTCTCCC | NM_001038700 |
| ***Fyco1* (217 bp, 60°C)** | Fw- CCCACGAGCGTGTGCTAC<br>Rev- ATCTGTGATTGGCTCTCCGC | NM_001164227.1 |

| | | |
|---|---|---|
| ***Fat1* (215 bp, 60°C)** | Fw- GTCGAGAGCCACAGAGAAGG<br>Rev- TTAACTCCTGGCCACCACAG | NM_001081286 |

#### Suppl. Table 5

Used antibodies in microglia morphology and gliosis and synapse histology experiments

| **Primary antibodies** | | | | |
|---|---|---|---|---|
| **Primary antibody** | Host | IHC, dilution | Manufacturer | RRID |
| **Iba1** | Rabbit | 1:400 | FUJIFILM Wako Chemicals Europe GmbH, Neuss, Germany | AB_839504 |
| **GFAP** | Rat | 1:200 | Invitrogen by ThermoFisher Scientific, Waltham, USA | AB_2532994 |
| **PSD95** | Rabbit | 1:500 | Synaptic Systems GmbH, Goettingen, Germany | AB_887760 |
| **Synaptophysin** | Guinea pig | 1:1000 | Synaptic Systems GmbH, Goettingen, Germany | AB_1210382 |
| **CD11b** | Rat | 1:200 | Bio Rad Laboratories Inc., Hercules, USA | AB_1100616 |
| **CD68** | Mouse | 1:200 | Bio Rad Laboratories Inc., Hercules, USA | AB_322219 |
| **LAMP1** | Rabbit | 1:250 | Sigma-Aldrich, Deisenhofen, Germany | AB_10646194 |
| **Secondary antibodies** | | | | |
| **Fluorophore** | anti- | IHC, dilution | Manufacturer | RRID |
| **Alexa 488** | Rat, guinea pig | 1:500 | Invitrogen/Life Technologies | AB_2535794<br>AB_2534117 |
| **Alexa 647** | Rabbit, mouse | 1:500 | | AB_2535813<br>AB_2535804 |
